# Functional mapping of epigenetic regulators uncovers coordinated tumor suppression by the HBO1 and MLL1 complexes

**DOI:** 10.1101/2024.08.19.607671

**Authors:** Yuning J. Tang, Haiqing Xu, Nicholas W. Hughes, Samuel H. Kim, Paloma Ruiz, Emily G. Shuldiner, Steven S. Lopez, Jess D. Hebert, Saswati Karmakar, Laura Andrejka, D. Nesli Dolcen, Gabor Boross, Pauline Chu, Colin Detrick, Sarah Pierce, Emily L. Ashkin, William J. Greenleaf, Anne K. Voss, Tim Thomas, Matt van de Rijn, Dmitri A. Petrov, Monte M. Winslow

**Affiliations:** Department of Genetics, Stanford University School of Medicine, Stanford, CA 94305; Department of Biology, Stanford University, Stanford, CA 94305; Cancer Biology Program, Stanford University School of Medicine, Stanford, CA 94305; Walter and Eliza Hall Institute of Medical Research, Melbourne, and Department of Medical Biology, University of Melbourne, Melbourne, VIC, Australia, 3052; Department of Pathology, Stanford University School of Medicine, Stanford, CA 94305; Institute of Evolution, Centre for Ecological Research; and Centre for Eco-Epidemiology, National Laboratory for Health Security, Budapest, Hungary

**Author notes:** Corresponding Author: Monte M. Winslow.

## Abstract

Epigenetic dysregulation is widespread in cancer. However, the specific epigenetic regulators and the processes they control to drive cancer phenotypes are poorly understood. Here, we employed a novel, scalable and high-throughput *in vivo* method to perform iterative functional screens of over 250 epigenetic regulatory genes within autochthonous oncogenic KRAS-driven lung tumors. We identified multiple novel epigenetic tumor suppressor and tumor dependency genes. We show that a specific HBO1 complex and the MLL1 complex are among the most impactful tumor suppressive epigenetic regulators in lung. The histone modifications generated by the HBO1 complex are frequently absent or reduced in human lung adenocarcinomas. The HBO1 and MLL1 complexes regulate chromatin accessibility of shared genomic regions, lineage fidelity and the expression of canonical tumor suppressor genes. The HBO1 and MLL1 complexes are epistatic during lung tumorigenesis, and their functional correlation is conserved in human cancer cell lines. Together, these results demonstrate the value of quantitative methods to generate a phenotypic roadmap of epigenetic regulatory genes in tumorigenesis *in vivo*.

## INTRODUCTION

Epigenetic alterations are a hallmark of cancer (Hanahan 2022). Critical features of the epigenome, including DNA methylation (Liang et al. 2023), chromatin accessibility (Corces et al. 2018), histone modifications (Geffen et al. 2023), and spatial genome organization (Deng et al. 2022), are all altered during tumorigenesis. Despite increasingly detailed profiling of the tumor epigenome, these characterizations rarely point to the drivers of epigenetic changes and the functional role of most epigenetic regulatory genes during tumorigenesis remains unknown. Furthermore, the specific epigenetic modifications that influence tumor phenotypes are still unclear. Given the widespread epigenetic alterations in cancer, systematic identification of functional epigenetic regulators and insights into the molecular programs they regulate are essential to better understand tumor biology and develop more effective therapeutics.

The epigenome is controlled by the activity of many different genes and complexes with overlapping, specific, and coordinated functions (Marakulina et al. 2023). Epigenetic regulators are often members of large families with potential functional redundancy (Cenik and Shilatifard 2021). Conversely, individual complexes can have interchangeable subunits that dictate specific functions (Centore et al. 2020; Yokoyama et al. 2024). Finally, epigenetic regulators can interact with one another to control molecular programs and thus cellular phenotypes (Cenik and Shilatifard 2021). Despite these layers of complexity, studies on epigenetic regulators in cancer have largely focused on individual genes, and only a few epigenetic complexes have been extensively studied (Mabe et al. 2024). Moving from investigating single genes to delineating the network of epigenetic regulation in tumorigenesis would lead to a more comprehensive understanding of the mechanisms by which the cancer epigenome determines cancer phenotypes.

A pressing challenge in studying cancer epigenetics is disentangling the impact of every epigenetic family, complex, and complex member on cancer phenotypes. While CRISPR-Cas9-mediated functional genomics screens have provided important insights into cancer driver genes, they primarily utilize cell lines that represent the most advanced cancer cells and systems that are unable to recapitulate the physiological tumor environment (Chow and Chen 2018). In contrast, genetically engineered mouse models of cancer allow functional investigation of defined genetic alterations on phenotypes throughout tumor development (Gengenbacher et al. 2017; Tang et al. 2023b). Tumors in these models are initiated from normal somatic cells within their natural environment, and recapitulate the histological, transcriptional and epigenetic profiles of their corresponding human cancers (LaFave et al. 2020; Marjanovic et al. 2020). However, only a small number of epigenetic regulators have been investigated individually using autochthonous cancer models (Walter et al. 2017; Concepcion et al. 2022).

Multiplexed somatic CRISPR-mediated gene editing has been performed in genetically engineered mouse models (Wang et al. 2018; Cai et al. 2021; Langille et al. 2022). Nonetheless, previous methods have yielded low resolution data analyzing only the largest dissectible tumors (Langille et al. 2022) or combining the growth effects of target genes from all tumors (Wang et al. 2018). Incorporating genetic barcodes into *in vitro* and *in vivo* CRISPR screens provides clonal resolution to phenotypic readouts, which drastically improves both the sensitivity to detect effects and the breadth of phenotypes that can be measured (Michlits et al. 2017; Rogers et al. 2017; Cai et al. 2021; Foggetti et al. 2021; Hughes et al. 2022). However, previous *in vivo* methods were constrained by scale.

In this study, we developed a scalable and quantitative method to perform high-throughput functional screens on large panels of genes in autochthonous cancer models with clonal resolution. Across iterative *in vivo* screens, we systematically dissected the functional impacts of >250 epigenetic regulatory genes on multiple aspects of lung tumorigenesis. Coupling this functional map with molecular analyses highlighted the HBO1 and MLL1 complexes as key interconnected tumor-suppressive epigenetic regulatory complexes in lung tumors.

## RESULTS

### U6-barcoding enables CRISPR screening with clonal resolution

To enable scalable and high-throughput quantification of genetic perturbations on cellular growth, we designed a system in which a clonal barcode is encoded within the 20-nucleotide region at the 3’ end of the U6 promoter directly adjacent to the sgRNA (**Figure 1A**, **Supplemental Figure 1A, Methods**). This system ensures proper coupling of genetic perturbation and clonal barcode, which can otherwise be uncoupled by template switching during lentiviral reverse transcription as well as by PCR during library preparation (Hegde et al. 2018; Hill et al. 2018). U6-barcoding enables pooled cloning and viral packaging, thus drastically increasing the scale and throughput of CRISPR screens with clonal resolution, and has immediate applications in diverse CRISPR screening approaches (**Supplemental Figure 1B**) (Michlits et al. 2017; Rogers et al. 2017; Rogers et al. 2018; Tang et al. 2023b; Hebert et al. 2024a; Hebert et al. 2024b)

**Figure 1.**
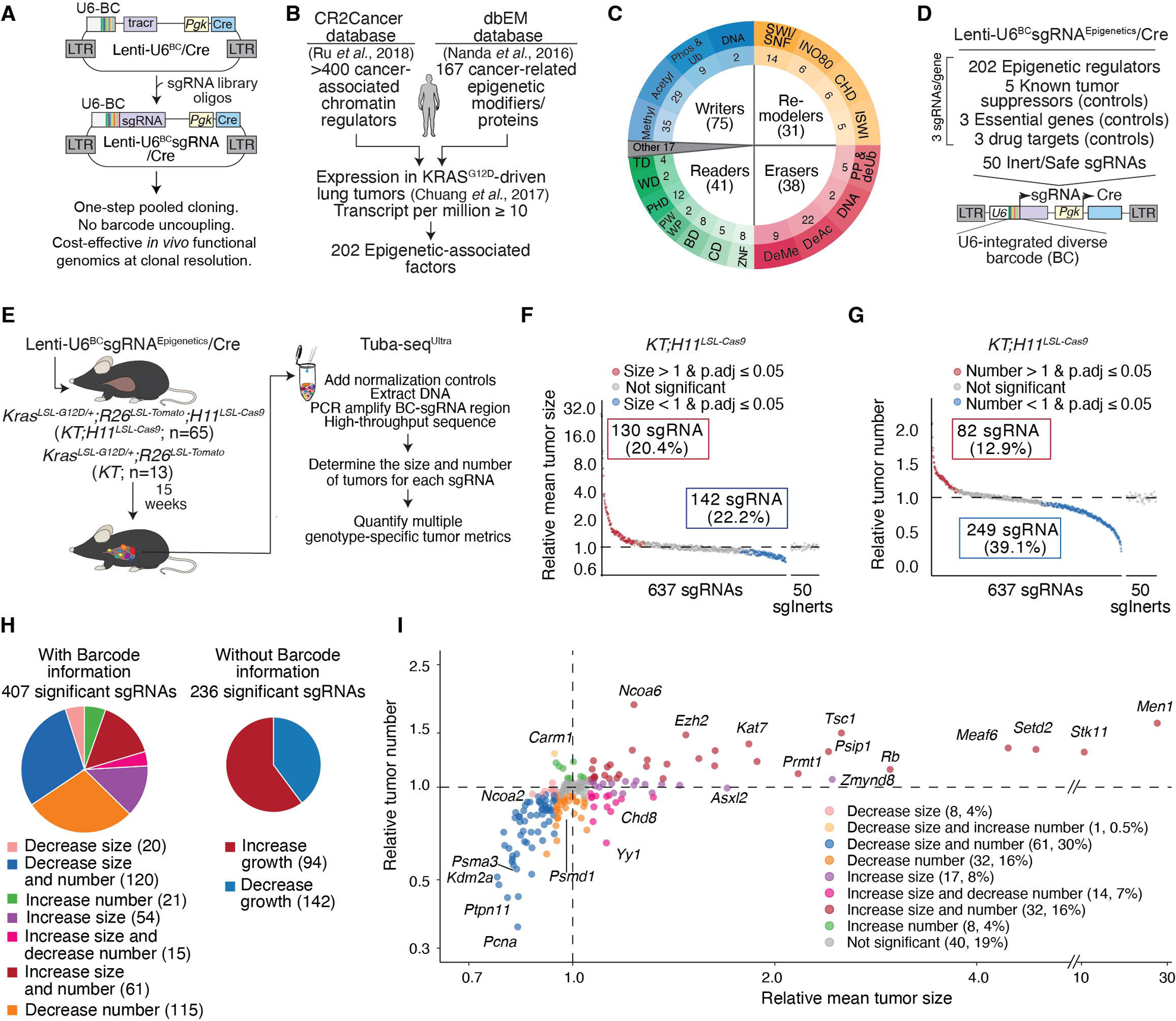
High-throughput analyses uncover the functional landscape of epigenetic regulation during autochthonous lung tumorigenesis. **A.** Genes were prioritized from two published databases of putative cancer-associated epigenetic regulators. To be included in the library, genes needed to be expressed at a transcript per million (TPM) ≥ 10 in KRAS^G12D^-driven lung cancer cells from analogous genetically engineered mouse model. **B.** Two-hundred and two genes from broad classes of epigenetic genes were prioritized for analysis in lung tumorigenesis *in vivo* (**see Supplemental Table 1**). **C.** The Tuba-seq^Ultra^ Lenti-U6^BC^sgRNA^Epigenetics^/Cre library contained sgRNAs targeting 202 epigenetic regulators (3 sgRNAs/gene) as well as sgRNAs targeting known tumor suppressors, drug targets, and essential genes. Additionally, 50 vectors with Inert/Safe targeting sgRNAs were included to generate tumors driven solely by oncogenic KRAS. **D.** Generation of Tuba-seq^Ultra^ (**U**6 barcode **L**abeling with per-**T**umor **R**esolution **A**nalysis) lentiviral libraries with a barcoded U6 promoter enables amplification and sequencing of the barcode-sgRNA region to provide genotype and clonal data. **E.** Tumors were initiated using the Lenti-U6^BC^sgRNA^Epigenetics^/Cre library in the indicated numbers of *Kras^LSL-G12D/+^;R26^LSL-Tomato^;H11^LSL-Cas9^* (*KT;H11^LSL-Cas9^*) and *KT* mice. *KT* mice, which lack Cas9, were used to quantify the relative titer of each vector in the sgRNA library and assess tumor growth effects. Tumors developed for 15 weeks before Tuba-seq^Ultra^ analysis. **F-G.** The impact of each sgRNA on tumor growth (Relative mean tumor size: log-normal (LN) mean tumor size relative to *sgInert* tumors) and tumor initiation/early growth (Relative tumor number) in *KT;H11^LSL-Cas9^*mice. Each dot is a sgRNA. Significant sgRNAs are colored as indicated (**see Methods and Supplementary Figure 2C**). **H.** Clonal-level analyses deconvolute the effect of different alterations on tumor size and number and uncover many more effects than sgRNA-level analyses without barcode-level information (computed here as Relative tumor burden, **see Methods**). **I.** Gene-level analysis of the impact of inactivating each gene on tumor growth (Relative mean tumor size) and tumor initiation/early growth (Relative tumor number). Each dot is a gene. Established tumor suppressors, essential genes, top novel epigenetic tumor suppressors, and representative genes in other categories are labeled. Note the split x-axis.

We previously developed a method that allows parallel analysis of gene function in autochthonous cancer models using tumor barcoding coupled with high-throughput barcode sequencing (Tuba-seq) (Rogers et al. 2017). This method uniquely barcodes each lentiviral-induced clonal tumor enabling barcode sequencing of bulk tumor-bearing tissue to measure the size of each tumor. Integrating barcodes with somatic genome editing allows the impact of different genetic perturbations on tumor size and number to be quantified (**Supplemental Figure 1C-D**) (Rogers et al. 2017; Cai et al. 2021; Li et al. 2021). Introducing U6-barcoding with Tuba-seq dramatically reduces the cost and effort required to perform barcode based CRISPR screens (**Supplemental Figure 1E-F**). This Tuba-seq^Ultra^ (**U**6 barcode **L**abeling with per-**T**umor **R**esolution **A**nalysis) approach provides high-throughput, quantitative, and highly sensitive analysis of large numbers of genetic perturbations on clonal growth in parallel within autochthonous mouse models.

### Chromatin regulators broadly impact lung tumorigenesis *in vivo*

Epigenetic dysregulation is a fundamental hallmark of cancer (Hanahan 2022); thus, we sought to investigate the role of every major category of epigenetic regulators during lung adenocarcinoma initiation and growth *in vivo*. Genes were selected by integrating databases of epigenetic regulators that may be disrupted in human cancers based on criteria such as mutational status, structural alterations, expression levels and/or proteomic data (Singh Nanda et al. 2016; Ru et al. 2018) and further selected based on their expression in mouse KRAS-driven lung cancer cells (**Figure 1B**) (Chuang et al. 2017). We generated a Tuba-seq^Ultra^ library containing 3 sgRNAs targeting each of >200 epigenetic regulators, 5 canonical tumor suppressor genes, 3 known drug targets, and 3 essential genes, as well as 50 inert sgRNAs (Lenti-U6^BC^sgRNA^Epigenetics^/Cre library; **Figure 1C-D** and **Supplemental Table 1**).

We initiated tumors with the Lenti-U6^BC^sgRNA^Epigenetics^/Cre library in *Kras^LSL G12D/+^*;*R26^LSL-Tomato^*;*H11^LSL-Cas9^* (*KT;H11^LSL-Cas9^*; n=65) and Cas9-negative *Kras^LSL-G12D/+^;R26^LSL-Tomato^* (*KT*; n=13) control mice via intratracheal injection (**Figure 1E**). Cas9-negative *KT* mice enable us to determine the representation of each sgRNA in the pool and whether any sgRNAs affect tumorigenesis in the absence of Cas9 (**Supplemental Figure 2**). Fifteen weeks after tumor initiation, all mice had a high tumor burden based on lung weight (**Supplemental Figure 3A**). We extracted DNA from bulk tumor-bearing lungs, PCR-amplified the barcode-sgRNA regions, and used Tuba-seq^Ultra^ to quantify the impact of inactivating each epigenetic regulator on tumor initiation and growth (**Supplemental Figure 2**). We quantified the size of >3×10^6^ clonal tumors, with a median of ∼4,500 tumors per sgRNA (**Supplemental Figure 3B-C**).

Many sgRNAs significantly increased or decreased the size and number of tumors in *KT;H11^LSL-Cas9^* mice, underscoring the broad and bidirectional impact of perturbing epigenetic regulators on lung tumorigenesis (**Figure 1F-G**, **Supplemental Figure 3, Supplemental Table 2, 3**). Inert sgRNAs in *KT;H11^LSL-Cas9^* mice did not affect tumorigenesis, and no sgRNAs had effects in *KT* control mice (**Figure 1F-G**, **Supplemental Figure 3D-E**). With per-tumor resolution, we identified 40% more statistically significant sgRNAs than we would have in a conventional CRISPR screen without barcodes (**Figure 1H**). These data show the fidelity of Tuba-seq^Ultra^ for highly multiplexed somatic genetic analyses.

Analysis at the gene level uncovered that inactivating >70% of epigenetic regulators had significant functional impacts on at least one facet of lung tumorigenesis, demonstrating the importance of epigenetic regulation in lung tumorigenesis (**Figure 1I**). Furthermore, quantifying the size and number of tumors generated by each sgRNA distinguished diverse functional phenotypes of the targeted genes (**Figure 1H-I**). Specifically, inactivating different epigenetic regulators can positively and/or negatively affect tumor size and/or number (**Figure 1I**).

### The HBO1 and MLL1 complexes are novel suppressors of lung cancer

We identified multiple novel tumor-suppressive and tumor dependency genes, many of which have never been implicated in lung tumors (**Figure 2A-D, Supplemental Table 4**). Inactivation of all control canonical tumor suppressor genes, including *Stk11*, *Setd2, Rb1*, *Rasa1*, and *Tsc1*, increased tumor growth (relative tumor size) and/or initiation (relative tumor number) at magnitudes consistent with previous studies (**Figure 2A-B**) (Cai et al. 2021). Meanwhile, inactivation of known essential genes and drug targets significantly decreased tumor initiation and/or growth (**Figure 2C-D**) (Cai et al. 2021; Blair et al. 2023). Most of the top epigenetic tumor suppressor genes increased both tumor size and number. However, inactivating *Epc1* and *Epc2* specifically increased tumor number but not tumor size, suggesting that they primarily affect tumor initiation (**Figure 2B, Supplemental Table 4**). Interestingly, EZH2 and PRMT1, which are oncogenic and candidate drug targets in other cancer types (Duan et al. 2020; Hwang et al. 2021), are tumor suppressors in autochthonous lung tumor (**Figure 2A, Supplemental Figure 4A**). Finally, we identified many tumor dependency genes, several of which have not previously been shown to affect tumor growth, such as *Chd2*, *Dhx30*, and *Gtf3c1* (**Figure 2C-D**).

**Figure 2.**
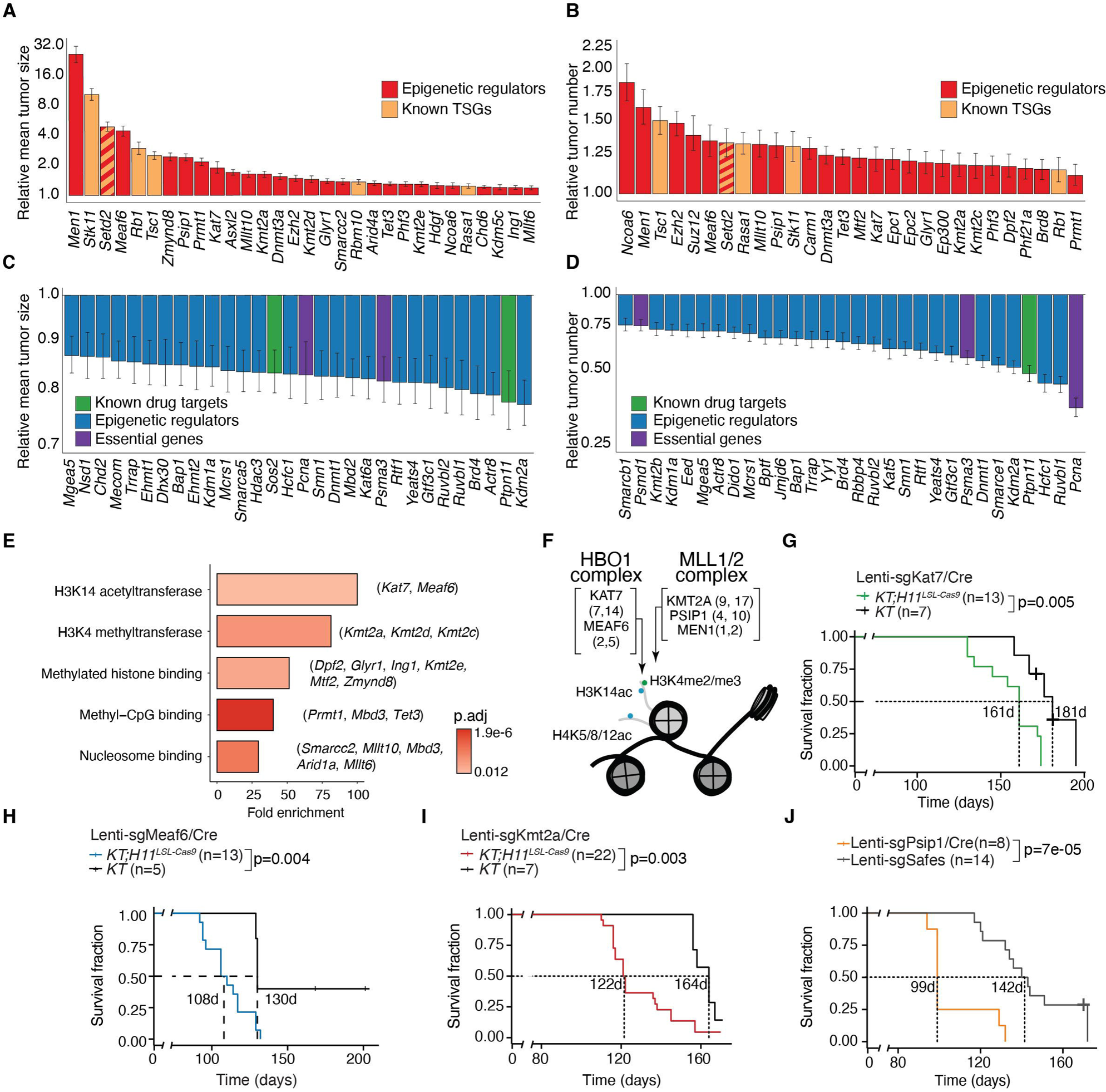
Top tumor-suppressive and tumor-dependency epigenetic regulators in lung tumorigenesis. **A-B.** The top 30 tumor-suppressive genes from the Lenti-U6^BC^sgRNA^Epigenetics^/Cre library that impact tumor size (**A**) and/or tumor number (**B**). Each bar represents the relative log-normal mean tumor size (**A**) or relative tumor number (**B**) of aggregated data from all sgRNAs for that gene (**see Methods**). The error bars are 95% confidence intervals. Note that *Setd2* is both a known tumor suppressor in lung cancer and an epigenetic regulator. **C-D.** The top 30 tumor-dependency genes from the Lenti-U6^BC^sgRNA^Epigenetics^/Cre library that impact tumor size (**C**) and/or tumor number (**D**). Each bar represents the relative log-normal mean tumor size (**C**) or relative tumor number (**D**) of aggregated data from all sgRNAs for that gene (**see Methods**). The error bars are 95% confidence intervals. **E.** Functional annotation from Gene Ontology of tumor-suppressive epigenetic regulators. Gene hits for each functional category are shown in brackets. Statistical significance is determined by Fisher’s exact test with Benjamini-Hochberg correction. **F.** Diagram showing the function of the HBO1 and MLL1/2 complexes and the tumor-suppressive subunits in the Lenti-U6^BC^sgRNA^Epigenetics^/Cre library. The numbers in brackets represent the gene ranking by effect size from largest to smallest for tumor size and tumor number (first value, second value) out of the 202 total genes in the Lenti-U6^BC^sgRNA^Epigenetics^/Cre library. **G-J.** Kaplan-Meier survival curves for inactivating *Kat7* (**G**), *Meaf6* (**H**), *Kmt2a* **(I)**, and *Psip1* (**J**) individually with a single sgRNA in *KT;H11^LSL-Cas9^* and *KT* mice (**G-I**), or in *KT;H11^LSL-Cas9^* mice transduced with *sgSafe* (**J**). The dashed lines and numbers indicate median survival. Statistical significance is determined by logrank test. Note the split x-axis.

Tumor-suppressive epigenetic regulators were enriched for histone 3 lysine 14 acetylation (H3K14ac) and histone 3 lysine 4 methylation (H3K4me) activity (**Figure 2E** and **Supplemental Table 5**). Five of the top ten genes with the largest effects on tumor growth are in protein complexes linked to H3K14ac and H3K4me (**Figure 2F**). The lysine acetyltransferase *Kat7* and *Meaf6* are members of the HBO1 complex that deposits H3K14ac (Kueh et al. 2011), and inactivation of either gene significantly increased tumor size and number (**Figure 2A-B, F**, **Supplemental Figure 4B**). Furthermore, inactivating *Hdac3* or *Sirt1*, which can deacetylate H3K14ac (Suter et al. 2012; Wang et al. 2019), decreased tumor size and number (**Figure 2C**, **Supplemental Figure 4C, Supplemental Table 4**). Inactivation of several genes in the MLL1/2 complexes, including *Kmt2a, Men1*, and *Psip1*, which are primarily involved in H3K4me2/3 (Wang and Helin 2024), also increased lung tumor growth (**Figure 2A-B, F**, **Supplemental Figure 4D**). Tumor dependency genes were involved in H4K16ac and H3K9me, as well as chromatin remodeling, histone deacetylation, and histone demethylation (**Supplemental Figure 4E, Supplemental Table 6**).

By inactivating genes simultaneously in the same animals, our approach removes confounding variability that arises from differences between animals and allows direct comparison of effect sizes between genes. We elected to focus further on novel tumor-suppressive epigenetic regulators because their magnitudes of effect are similar to or exceed those of canonical tumor suppressor genes like *Rb1* and *Tsc1* (**Figure 2A**), and their molecular roles in lung tumorigenesis are entirely unknown.

To validate the tumor-suppressive effects of the catalytic and structural subunits of the HBO1 and MLL1/2 complexes outside of a pooled setting, as well as to generate material for subsequent molecular analyses, we inactivated genes individually in *KT* and *KT;H11^LSL-Cas9^* mice. Inactivation of *Kat7*, *Meaf6*, *Kmt2a* or *Psip1* all significantly reduced survival, consistent with their function as tumor suppressors (**Figure 2G-J**). Immunohistochemistry for NKX2.1 and P63 confirmed that the tumors in which these genes were inactivated were lung adenomas/adenocarcinomas (**Supplemental Figure 4F**). Taken together, the quantitative *in vivo* screening and subsequent single-gene validation demonstrates that the HBO1 and MLL1/2 complexes are two of the most important epigenetic complexes that restrain lung tumorigenesis.

### Saturation analysis defines a distinct tumor-suppressive HBO1 complex variant and underscores the unique impact of the MLL1 complex

The HBO1 complex is a member of the larger, highly conserved MYST family of histone acetyltransferases (Avvakumov and Cote 2007; Voss and Thomas 2009), while the MLL1 complex is a member of the COMPASS family of methyltransferase complexes (Cenik and Shilatifard 2021). To fully elucidate the role of the MYST and COMPASS families in lung tumorigenesis, we generated a Tuba-seq^Ultra^ library that included key genes from our initial analysis, and 47 additional chromatin regulators. This library targeted every COMPASS and MYST family member, histone demethylases, and other related genes of interest (Lenti-U6^BC^sgRNA^Saturation^/Cre library; **Figure 3A-B, Supplemental Table 7**). We initiated tumors with the Lenti-U6^BC^sgRNA^Saturation^/Cre library in *KT;H11^LSL-Cas9^* (n=33) and *KT* (n=7) mice (**Supplemental Figure 5A-D**). Tuba-seq^Ultra^ analysis uncovered many additional epigenetic regulators that impact tumor size and/or tumor number (**Figure 3C-D, Supplemental Figure 5E-J, and Supplemental Table 8, 9**).

**Figure 3.**
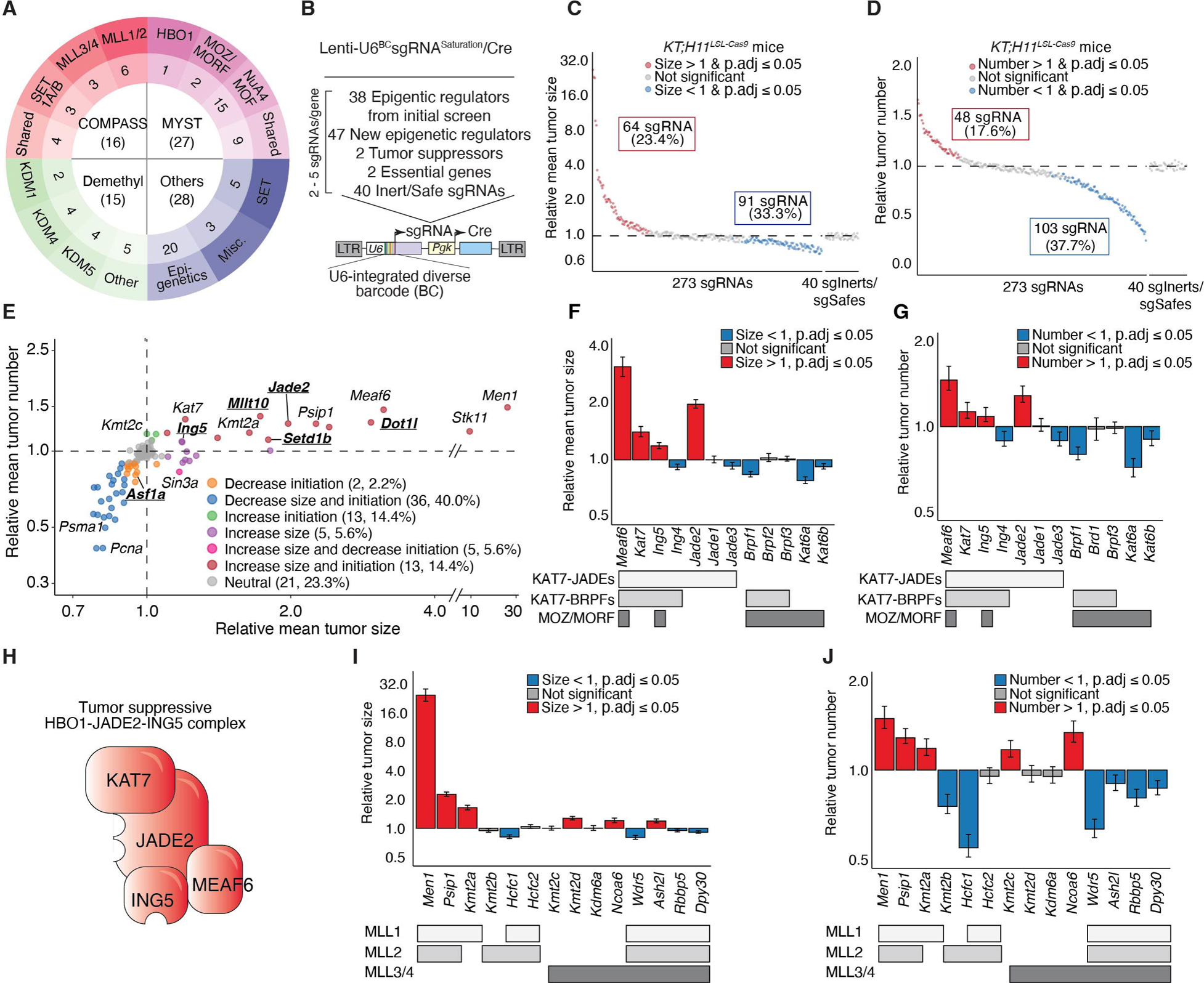
MYST and COMPASS saturation analyses uncover specific tumor-suppressive complexes. **A.** The MYST and COMPASS family genes, as well as other genes of interest that were included for functional analysis in lung tumorigenesis *in vivo* (**see Supplemental Table 7**). **B.** The Lenti-U6^BC^sgRNA^Saturation^/Cre library contains sgRNAs targeting 38 epigenetic regulators (3-4 sgRNAs/gene) from the initial analyses (Figures 1-2), 47 new epigenetic regulators, as well as tumor suppressors, essential genes, and Inert/Safe-targeting sgRNAs. **C-D.** Impact of each sgRNA on tumor size (**C**) and tumor number (**D**) in *KT;H11^LSL-Cas9^* mice. Each dot is a sgRNA. Significant sgRNAs are colored as indicated. **E.** Gene-level analysis of the impact of inactivating each gene in the Lenti-U6^BC^sgRNA^Saturation^/Cre library on tumor size and tumor number. Each dot is a gene. Established tumor suppressors, essential genes, top novel epigenetic tumor suppressors, and representative genes in other classes are labeled. Newly included genes are bolded and underlined. Note the split x-axis. **F-G.** Effects of inactivating genes from the HBO1 or the MOZ/MORF complexes on tumor size **(F)** and tumor number **(G)**. Bars are aggregate data of all sgRNAs for each gene. Error bars are 95% confidence intervals. The complex(es) each subunit belongs to is indicated at the bottom of the graph. **H.** Schematic shows the tumor-suppressive HBO1^JADE2-ING5^ complex, where all subunits are tumor suppressor genes. **I-J.** Effects of inactivating genes from the MLL1/2/3/4 complexes on tumor size **(I)** and number **(J)**. Bars are aggregated data of all sgRNAs for each gene. Error bars are 95% confidence intervals. The complex(es) each subunit belongs to is indicated at the bottom of the graph.

In addition to KAT7 and MEAF6, the HBO1 complex can contain either ING4 or ING5 and one of JADE1/2/3 or BRPF1/2/3 (Avvakumov and Cote 2007). Among these alternative subunits, our Tuba-seq^Ultra^ analysis identified ING5 and JADE2 as tumor-suppressive HBO1 subunits (**Figure 3E-G**, **Supplemental Figure 5K-L, Supplemental Table 10**). Inactivation of ING4, other JADEs and all BRPFs either had no effect or decreased lung tumorigenesis *in vivo* (**Figure 3F-G**). These differences were not driven by differences in the expression level of the subunits (**Supplemental Figure 5M**). Thus, KAT7/MEAF6/JADE2/ING5 comprise the tumor-suppressive variant of the HBO1 complex in lung adenocarcinoma (**Figure 3E-H**). Importantly, none of the other genes in the MYST family were tumor suppressors (**Supplemental Figure 6A-B, Supplemental Table 10**). Inactivating *Kat6a* or *Kat6b*, the catalytic subunits of the MOZ/MORF complexes, or *Kat8*, the catalytic subunit of the male-specific-lethal (MSL) and the non-specific-lethal (NSL) complexes, or *Kat5*, the catalytic subunit of the TIP60 complex, led to modest decreases in tumor growth (**Figure 3F-G**, **Supplemental Figure 6C-D**).

Among the COMPASS family complex components, those within the MLL1/2 complexes had the highest magnitudes of tumor suppression (**Figure 3I-J**, **Supplemental Figure 6A-B, Supplemental Table 10**). However, inactivation of *Kmt2b*, the catalytic subunit of the MLL2 complex and a paralog of *Kmt2a*, did not promote tumor growth and reduced tumor number (**Figure 3I-J**). Tumors initiated with dual-sgRNA lentiviral vectors indicated that coincident inactivation of *Kmt2a* and *Kmt2b* further reduced tumorigenesis (Hebert et al. 2024a) (**Supplemental Figure 6E-F**). Thus, while the MLL1 complex has an unique tumor-suppressive role, MLL1 and MLL2 complexes are partially redundant in controlling an essential function in lung tumors, consistent with data from other cell types (Denissov et al. 2014; Chen et al. 2017).

By systematically targeting entire gene families, we dissected the functional impact of MYST and COMPASS genes in driving lung tumorigenesis *in vivo* and demonstrated that paralogs can have distinct functions. We pinpointed the HBO1^JADE2-ING5^ complex and the MLL1 complex as suppressors of lung tumorigenesis.

### HBO1 complex deficiency disrupts histone acetylation in lung tumors

The HBO1 complex catalyzes acetylation of several lysine residues on the tails of histone 3 and histone 4 (Yokoyama et al. 2024). To assess which histone modifications are regulated by the HBO1 complex in lung cancer, we first assessed H3K14ac (Kueh et al. 2011; MacPherson et al. 2020; Kueh et al. 2023; Yokoyama et al. 2024). Immunohistochemistry showed that CRISPR-mediated *Kat7* or *Meaf6* inactivation greatly reduced H3K14ac (**Figure 4A, Supplemental Figure 7A**). Western blot analysis of FACS isolated neoplastic cells from *KT;H11^LSL-Cas9^* mice with Lenti-sgKat7/Cre-initiated lung tumors (hereafter referred to as *sgKat7* cells) confirmed a significant reduction in KAT7 and H3K14ac. The residual KAT7 protein and H3K14ac could result from incomplete inactivation of *Kat7* by CRISPR in a subset of tumors (**Figure 4B-C**). H4K12ac and H4K5ac are other histone acetylation marks attributed to the HBO1 complex (Doyon et al. 2006). H4K12ac but not H4K5ac was reduced in *sgKat7* cells (**Figure 4B-C, Supplemental Figure 7B-C**). Similarly, H3K14ac and H4K12ac, but not H4K5ac were reduced in *sgMeaf6* cells (**Figure 4D-E, Supplemental Figure 7A, D-E**).

**Figure 4.**
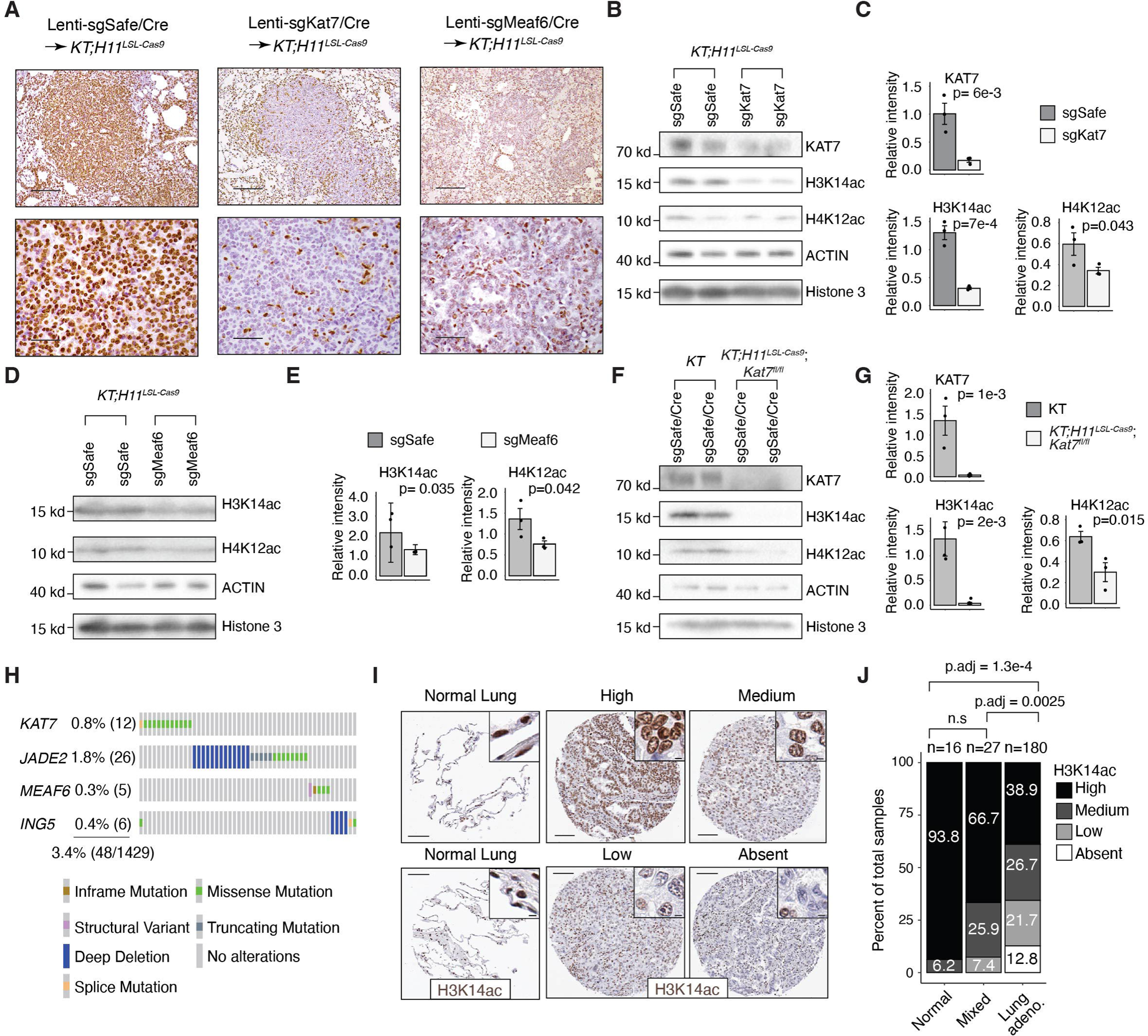
Disruption of the HBO1 complex histone modification targets in mouse and human lung tumors. **A.** Representative immunohistochemistry staining for H3K14ac in mouse lung tumors initiated by different single sgRNAs in *KT;H11^LSL-Cas9^* and *KT* mice. Bottom panels (scale bars, 50 μm) are higher magnification images of areas in the top panel (scale bars, 100 μm). **B.** Representative western blots of FACS-isolated lineage (CD45/CD31/Ter119/F4/80) negative, tdTomato positive (Lin^−^TdTom^+^) neoplastic cells from *KT;H11^LSL-Cas9^* mice transduced with Lenti-sgKat7/Cre or Lenti-sgSafe/Cre vectors. Each lane on the western blot is a different mouse. **C.** Bar plots showing densitometry quantification of protein levels on western blots. Relative intensities are normalized to ACTIN. Error bars are +/− the standard error of the mean, and each dot represents a different mouse. Statistical significance is computed by Student’s t-test. **D.** Representative western blots of FACS-isolated Lin^−^TdTom^+^ neoplastic cells from *KT* and *KT;H11^LSL-Cas9^;Kat7^fl/fl^*mice transduced with Lenti-sgSafe/Cre vectors. Each lane on the western blot is a different mouse. **E.** Bar plots showing densitometry quantification of protein levels on western blots. Relative intensities are normalized to ACTIN. Error bars are +/− the standard error of the mean, and each dot represents a different mouse. Statistical significance is computed by Student’s t-test. **F.** Representative western blots of FACS-isolated Lin^−^TdTom^+^ neoplastic cells from *KT;H11^LSL-Cas9^* mice transduced with Lenti-*sgMeaf6*/Cre or Lenti-*sgSafe*/Cre vectors. Each lane on the western blot is a different mouse. **G.** Bar plots showing densitometry quantification of protein levels on western blots. Relative intensities are normalized to ACTIN. Error bars are +/− the standard error of the mean, and each dot represents a different mouse. Statistical significance is computed by Student’s t-test. **H.** Type and frequency of mutations for different subunits in the HBO1^JADE2-ING5^ tumor-suppressive complex in human lung adenocarcinoma. Numbers indicate the number of patients. **I.** Representative immunohistochemistry images from a human lung cancer tissue microarray (TMA) expressing high, medium, low, or absent levels of H3K14ac (scale bars, 50 μm. Inset scale bars, 5 μm). **J.** Bar plot showing the percentage of samples from human TMAs that express different levels of H3K14ac in normal lung (Normal), mixed lung adenocarcinoma with bronchioalveolar lung carcinoma (Mixed), and lung adenocarcinoma. Statistical significance is computed by Kruskal-Wallis test with Benjamini-Hochberg correction.

To further validate the role of the HBO1 complex in histone modification in an orthogonal model in which all tumors would be *Kat7*-deficient, we generated lung tumors in *KT* and *KT;H11^LSL-Cas9^;Kat7^fl/fl^*mice (**Methods**, **Supplementary Figure 7F**) (Kueh et al. 2011; Kueh et al. 2023). Lung tumors in *KT;H11^LSL-Cas9^;Kat7^fl/fl^*mice were higher grade as evidenced by increased nuclear pleomorphism, reduced nucleus to cytoplasm ratio and increased papillary features compared to control tumors in *KT* mice (**Supplemental Figure 7G-H**). Moreover, neoplastic cells from lung tumors in *KT;H11^LSL-Cas9^;Kat7^fl/fl^*mice lacked KAT7 protein, had extremely low/absent H3K14ac, and had modestly but consistently reduced H4K12ac as assessed by immunohistochemistry and western blotting (**Figure 4F-G** and **Supplemental Figure 7I-K**). Thus, in lung adenocarcinoma, H3K14ac is predominantly if not exclusively generated by the HBO1 complex, with effects on H4K12ac being more modest, consistent with previous reports (Au et al. 2021; Kueh et al. 2023; Yokoyama et al. 2024).

### Human lung adenocarcinomas exhibit genetic alterations in the HBO1^JADE2-ING5^ complex variant and frequent reductions in H3K14ac

While mutation frequency was not an initial criterion to include genes in our broad functional screen of epigenetic regulators, given the identification of the tumor-suppressive HBO1^JADE2-ING5^ complex, we queried published human lung adenocarcinoma genome sequencing data for mutations and DNA copy number changes in these genes (Cerami et al. 2012; Gao et al. 2013; de Bruijn et al. 2023). Mutations in *KAT7* occurred in ∼1% of tumors and are largely mutually exclusive with mutations and deep deletions in *JADE2, ING5* and *MEAF6* (**Figure 4H**). Overall, ∼3% of human lung adenocarcinomas had point mutations or deep deletions in a member of the HBO1^JADE2-ING5^ complex (**Figure 4H**). Tumor suppressor genes in the MLL1 complex, including *KMT2A*, *PSIP1*, and *MEN1*, had point mutations or deep deletions in ∼5% of human lung adenocarcinomas. (**Supplemental Figure 8A**). In addition, genetic alterations in the subunits of the HBO1^JADE2-ING5^ and/or MLL1 tumor suppressor complexes are associated with reduced overall survival of lung adenocarcinoma patients (**Supplemental Figure 8B-D**).

Given that tumor suppressor genes are frequently inactivated by non-mutational mechanisms, we next assessed the levels of H3K14ac, H4K12ac, and H4K5ac in >200 human lung adenocarcinomas by immunohistochemistry (**Figure 4I-J, Supplemental Figure 9A-E**). More than 30% of tumors had low or absent H3K14ac, and > 60% of tumors exhibited a decrease in H3K14ac, consistent with reduced HBO1 complex activity in a large fraction of human lung adenocarcinomas (**Figure 4J**). Lung tumors with mixed bronchioalveolar carcinoma histology, a precursor to lung adenocarcinoma with better clinical outcomes (Raz et al. 2006), were less likely to have low/absent H3K14ac. Similar to our mouse tumors, there was a modest reduction in H4K12ac levels in human lung adenocarcinomas (**Supplemental Figure 9A-B**) and H4K12ac levels significantly correlated with H3K14ac, suggesting that these modifications are coordinately regulated (**Supplemental Figure 9C**). Levels of H4K5ac were not different between human normal lung and cancer samples (**Supplemental Figure 9D-E**).

To contextualize the prevalence of HBO1 inactivation in human lung adenocarcinomas, we assessed H3K36me3, the histone modification catalyzed by SETD2, which is one of the most highly mutated *bona fide* tumor suppressor genes in human lung adenocarcinomas (Govindan et al. 2012; Cancer Genome Atlas Research 2014). Approximately ∼15% of tumors had reduced levels of H3K36me3 (**Supplemental Figure 9F-G**), comparable to previous data (Walter et al. 2017). Thus, the fraction of human lung adenocarcinoma with reduced/absent HBO1 complex activity is greater than or equal to the percent of tumors with reduced/absent SETD2 activity.

### The HBO1 and MLL1 complexes regulate accessibility at shared chromatin regions in lung adenocarcinoma

To further investigate the epigenetic consequences of inactivating the HBO1 complex in lung tumors, we performed Assay for Transposase-accessible chromatin with sequencing (ATAC-seq) on *sgKat7* cells (**Figure 5A, Supplemental Figure 10A-B, Supplemental Table 11**). These cells had significantly altered chromatin accessibility, with more regions becoming less accessible compared to *sgSafe* cells (**Figure 5B, Supplemental Table 12**). Over 35% of the less accessible regions were in promoter regions (**Figure 5C, Supplemental Table 13**). To better understand the epigenetic impacts of other HBO1 and MLL1 complex genes, we performed ATAC-seq on sorted neoplastic cells from lung tumors initiated with *sgMeaf6*, *sgKmt2a*, or *sgPsip1* expressing Lenti-sgRNA/Cre vectors. Similar to *Kat7*, inactivating these genes significantly affected chromatin accessibility, with more regions becoming less accessible in *sgMeaf6* and *sgPsip1* cells **(Supplemental Figure 10C, Supplemental Tables 14-16)**. Furthermore, 27% to 47% of the less accessible regions in *sgMeaf6*, *sgKmt2a*, or *sgPsip1* cancer cells were within promoters (**Figure 5C, Supplemental Figure 10D**). These observations are consistent with H3K14ac and H3K4me3, the primary histone modifications by the HBO1 and MLL1 complexes respectively, as marks that facilitate transcription at promoters (Kueh et al. 2011; MacPherson et al. 2020; Wang and Helin 2024).

**Figure 5.**
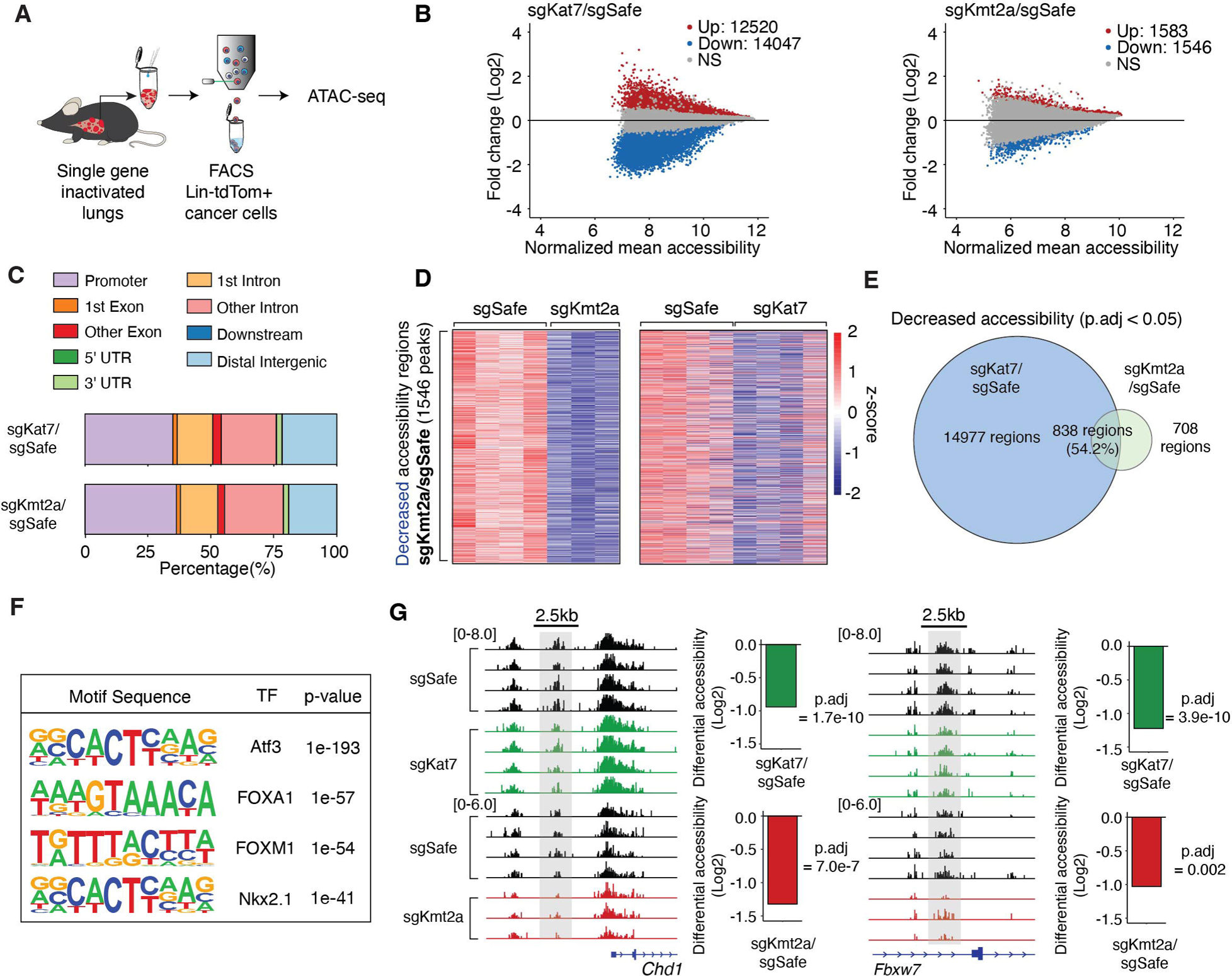
The HBO1 and MLL1 complexes regulate chromatin accessibility at shared regions in the genome. **A.** FACS-isolated lineage (CD45/CD31/Ter119/F4/80) negative, tdTomato positive (Lin^−^TdTom^+^) neoplastic cells from dissociated tumor-bearing mouse lungs are used for bulk ATAC-seq. **B.** Mean average (MA) plots of chromatin accessibility changes for *sgKat7* and *sgKmt2a* cells relative to *sgSafe* cells. Red and blue dots represent statistically significant ATAC-seq peaks (p.adj ≤ 0.05). Note that these data are from two separate ATAC-seq experiments with separate *sgSafe* samples for each experiment. **C.** Annotation and percent representation of genomic regions with significantly decreased chromatin accessibility peaks (cut-off: log2 fold-change < 0 & p.adj ≤ 0.05) for *sgKat7* and *sgKmt2a* cells relative to *sgSafe* cells. **D.** Heatmaps showing all significantly decreased chromatin accessibility regions in *sgKmt2a* cells relative to *sgSafe* cells (cut-off: log2 fold-change < 0 & p.adj ≤ 0.05). The same chromatin regions are shown for *sgKat7* cells relative to *sgSafe* cells, indicating a broad similarity of decreased chromatin accessibility between these two genotypes. **E.** Venn diagram showing the overlap in decreased chromatin accessibility regions between *sgKat7* cells relative to *sgSafe* cells and between *sgKmt2a* cells relative to *sgSafe* cells. **F.** Top transcription factor motifs enriched in shared genomic regions with decreased chromatin accessibility between *sgKat7* cells relative to *sgSafe* cells and between *sgKmt2a* cells relative to *sgSafe* cells. **G.** Genome accessibility tracks showing decreased chromatin accessibility upstream of transcription start sites for tumor suppressor genes *Chd1* and *Fbxw7* in *sgKat7* and *sgKmt2a* cells. Grey regions highlight peaks with significant differential chromatin accessibility. Numbers in brackets are the range of normalized peak counts for the tracks below. Bar plots show fold-change of differential chromatin accessibility for the indicated sgRNA relative to *sgSafe* cells.

Critically, the overall chromatin accessibility landscapes between tumors with inactivated HBO1 and MLL1 complex genes are significantly correlated (**Supplemental Figure 10E**). There were substantial overlaps in regions of reduced chromatin accessibility. For example, >50% of the less accessible regions in sg*Kmt2a* cells were also less accessible in sg*Kat7* cells (**Figure 5D-E, Supplemental Table 17**). Similar correlations and overlaps in chromatin accessibility existed between inactivating other HBO1 or MLL1 complex genes (**Supplemental Figure 10F-I**).

To better characterize the overlapping chromatin regions regulated by the HBO1 and MLL1 complexes, we examined the DNA-binding motifs and the genes that are associated with these regions. The shared regions of reduced chromatin accessibility between *sgKat7* and *sgKmt2a* cells were enriched for transcription factor motifs for FOXA1, FOXA2 and NKX2.1, which regulate lung morphogenesis and epithelial cell differentiation (**Figure 5F**, **Supplemental Table 18**) (Herriges and Morrisey 2014; Orstad et al. 2022). Reduced chromatin accessibility regions for the other inactivated genes enriched for similar transcription factors (**Supplemental Figure 11A, Supplemental Table 18**). In addition, genomic regions near known tumor suppressor genes such as *Chd1*, *Fbwx7*, and *Dusp5* had consistently reduced accessibility (**Figure 5G, Supplemental Figure 11B, Supplemental Tables 4, 12 - 16**) (Xiao et al. 2018; Kidger et al. 2022). Overlapping regions of increased chromatin accessibility were enriched for KLF and TEAD motifs (**Supplemental Figure 11C, Supplemental Table 19**), which are known to promote cell proliferation and regulate lung morphogenesis (McConnell and Yang 2010; Zhong et al. 2024). Taken together, our data suggest that the HBO1 and MLL1 complexes regulate the chromatin accessibility landscape in shared regions of the genome involved in lung development and tumor suppression.

### The HBO1 and MLL1 complexes regulate cell type identity and canonical tumor suppressor gene expression in lung adenocarcinoma

To investigate the gene expression programs regulated by the HBO1 complex in lung tumors, we performed single-cell RNA sequencing (scRNA-seq) on FACS-isolated neoplastic cells from *sgKat7* lung tumors (**Figure 6A, Supplemental Table 20**). The *sgKat7* cells were in clusters distinct from control *sgSafe* cells (**Figure 6B, Supplemental Figure 12A-D**). We compared gene expression between clusters composed predominantly of *sgKat7* cells relative to *sgSafe*, given that a subset of *sgKat7* cells will not have complete gene inactivation resulting from inefficiencies in CRISPR-Cas9-mediated gene inactivation (**Supplemental Figure 12D**). There were significant differences in the overall gene expression between *sgKat7* and *sgSafe* clusters, with a greater number of genes being down-regulated in *sgKat7* clusters, consistent with ATAC-seq profiling (**Figure 6C**, **Figure 5B, Supplemental Table 21**). Up-regulated genes were largely involved in metabolism and developmental pathways, while genes involved in cytoskeleton organization, cell motility, differentiation and apoptosis were down-regulated (**Figure 6D-E, Supplemental Table 22-23**).

**Figure 6.**
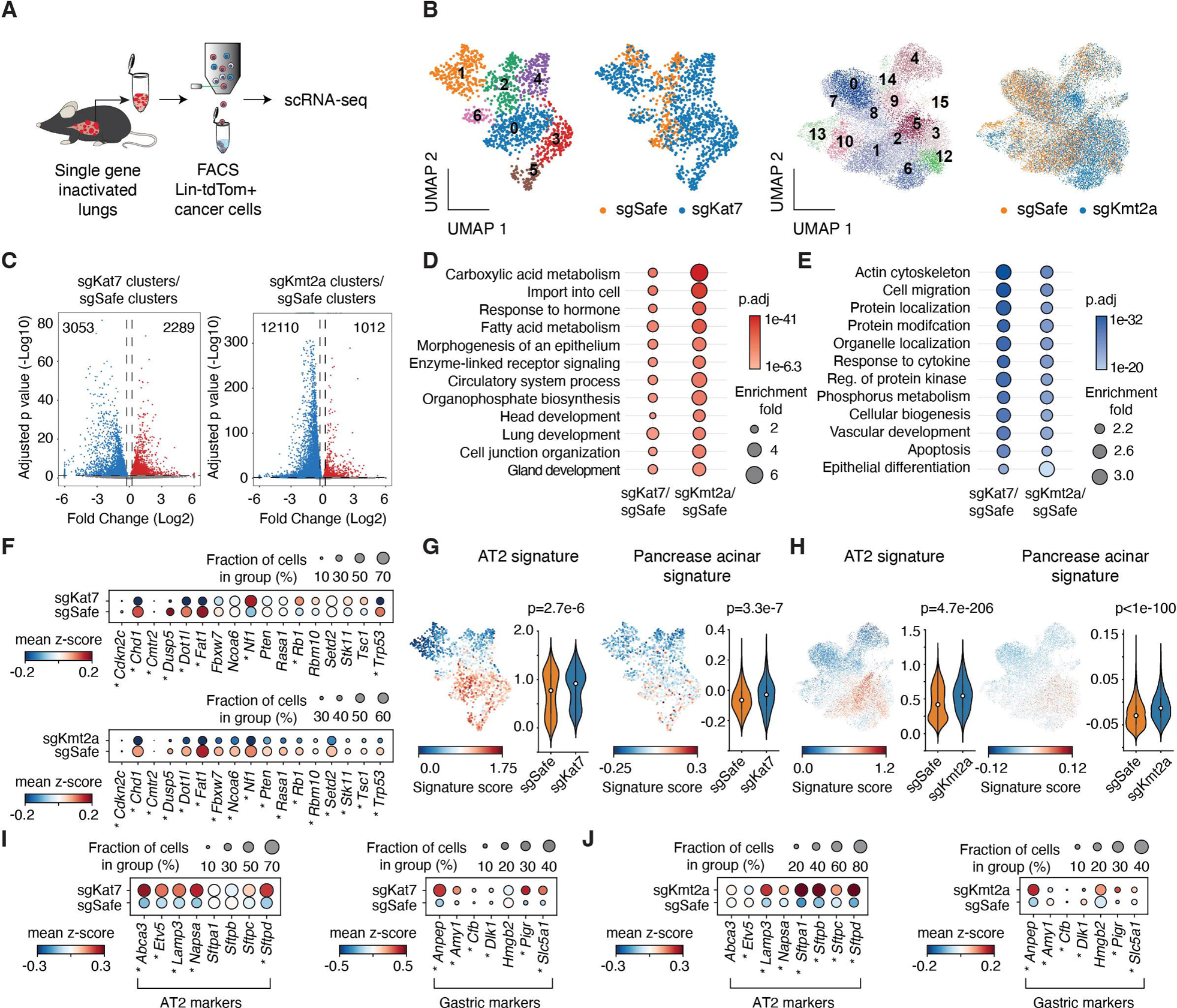
The HBO1 and MLL1 complex genes regulate cell state identity and canonical tumor suppressor gene expression. **A.** FACS-isolated lineage (CD45/CD31/Ter119/F4/80) negative, tdTomato positive (Lin^−^TdTom^+^) neoplastic cells from dissociated tumor-bearing mouse lungs are used for scRNA-seq. **B.** UMAPs, cell clusters and genotypes from scRNA-seq of tumor cells transduced with *sgKat7* or*sgSafe*, and *sgKmt2a* or *sgSafe*. **C.** Volcano plots of differential gene expression between cell clusters that are predominantly *sgKat7* or *sgKmt2a* compared to cell clusters that are predominantly *sgSafe*. The specific clusters for each group are indicated in **Figure S12A-B** (**see Methods**). Red dots are statistically significant up-regulated genes (log2 fold-change > 0 & p.adj ≤ 0.05), and blue dots are statistically significant down-regulated genes (log2 fold-change < 0 & p.adj ≤ 0.05). The total number of statistically significant genes is indicated at the top of the plot on each side. **D-E.** Top enriched gene ontology (GO) terms shared between statistically significant up-regulated genes (**D**) or down-regulated genes (**E**) for the indicated comparisons. The data are from differential gene expression between cell clusters that are predominantly *sgKat7* or *sgKmt2a* compared to cell clusters that are predominantly *sgSafe*. **F.** Dot plots from scRNA-seq showing differential gene expression of canonical tumor suppressor genes between the indicated cluster-based comparisons. * indicates p.adj ≤ 0.05 (Student’s t-test). **G-H.** UMAPs and volcano plots showing cell type signature scores for cell clusters that are predominately *sgKat7* or *sgSafe* (**G**), and *sgKmt2a* or *sgSafe* (**H**). Median cell type signature score is presented by the dot in each violin plot. Statistical significance is computed using Mann-Whitney-U test. **I-J.** Dot plots from scRNA-seq showing differential expression of AT2 and gastric cell markers between cell clusters that are predominantly *sgKat7* (**I**) or *sgKmt2a* (**J**) compared to cell clusters that are predominantly *sgSafe*. * indicates p.adj ≤ 0.05 (Student’s t-test).

To investigate the impact of perturbing other HBO1 complex genes as well as the MLL1 complex on transcriptional programs and cell states, we performed scRNA-seq on *sgMeaf6*, *sgKmt2a*, *sgPsip1* or *sgSafe* cells (**Figure 6B-C, Supplemental Figure 12E-R**). Similar to *sgKat7* cells, clusters enriched for *sgKmt2a* had more down-regulated genes than up-regulated genes, and many of the cellular processes that were enriched in *sgKat7* cell clusters were also enriched in *sgKmt2a* cell clusters (**Figure 6C-E, Supplemental Table 21-23**). Cell clusters that were predominantly *sgMeaf6* or *sgPsip1* also showed greater numbers of down-regulated genes and enrichment of cellular processes enriched in *sgKat7* cell clusters (**Supplemental Figure 13 A-D**, **Supplemental Table 22-23**). Multiple tumor suppressor genes, such as *Trp53*, *Stk11*, *Chd1*, *Dot1l*, *Dusp5*, *Fbxw7* or *Rb1*, were downregulated in cell clusters enriched for inactivated HBO1 or MLL1 complex genes (**Figure 6F**, **Supplemental Figure 13E**). These observations are congruent with the ATAC-seq analysis, further supporting that the HBO1 or MLL1 complexes govern shared molecular programs and the expression of canonical tumor suppressor genes in lung tumors.

We next examined the impact of HBO1 or MLL1 complex deficiency on cellular identity. Inactivating *Kat7*, *Kmt2a* or *Psip1* promoted a more AT2-like state, characterized by increased AT2 cell type signature score and increased expression of known AT2 marker genes (**Figure 6G-I, Supplemental Figure 13F-I, Supplemental Table 24**). Cell clusters of *sgMeaf6* cells were enriched for both AT2 and AT1 signatures (**Supplemental Table 24**). Furthermore, cells in clusters of sgRNAs targeting the HBO1 or MLL1 complex genes had enhanced gene signatures from other tissues such as the pancreas, liver and kidney (**Figure 6G-I, Supplemental Figure 13F-I, Supplemental Table 24**). The emergence of mixed lineage cell states indicates reduced lineage fidelity, where these cells can engage in a more diverse phenotypic space. Taken together, the scRNA-seq analyses revealed that inactivation of the HBO1 or MLL1 complexes consistently altered transcriptional programs associated with lung development, repressed the expression of canonical tumor suppressor genes and weakened lineage fidelity.

### KAT7 exhibits epistasis with the MLL1 complex and canonical tumor suppressor genes

Understanding genetic interactions can uncover redundant functions within and between complexes and pathways, as well as reveal important insights into cancer evolution and cellular fitness (Phillips 2008; Dixon et al. 2009; Wang et al. 2014; Hebert et al. 2024a). Given the shared tumor-suppressive effects of the HBO1 and MLL1 complexes and their considerable similarity in molecular features, we examined the fitness relationship between the HBO1 and MLL1 complexes in human cancer cells. Analysis of the Cancer Dependency Map Project (DepMap) (Tsherniak et al. 2017) showed that the tumor-suppressive HBO1^JADE2-ING5^ complex members had highly correlated fitness effects with one another, and with the tumor-suppressive genes from the MLL1 complex, but not with *KMT2B* (**Figure 7A, Supplemental Figure 14A-B, Supplemental Table 25**). Importantly, the correlation coefficients between genes across the HBO1 and MLL1 complexes were comparable to genes within each complex. In addition, *ING4* and *JADE1/3* were less correlated than *ING5* and *JADE2* with tumor-suppressive MLL1 genes (**Supplemental Figure 14B, Supplemental Table 25**). These fitness correlations from >1,000 human cancer cell lines support a functionally distinct HBO1^JADE2-ING5^ complex that interacts with the MLL1 complex.

**Figure 7.**
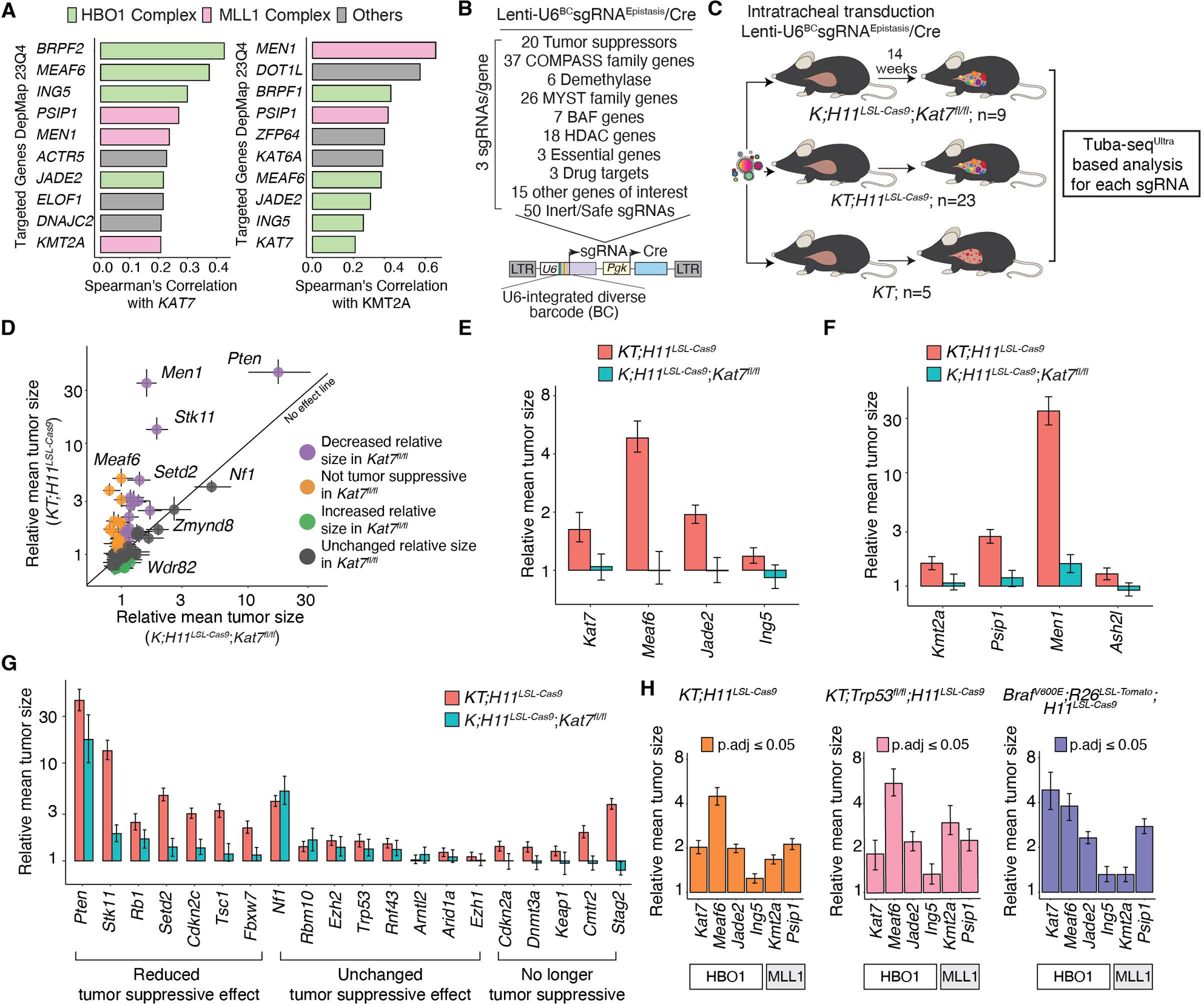
*Kat7* is epistatic with the MLL1 complex and canonical tumor suppressor genes. **A.** Correlation of cancer dependency scores from DepMap showing the top genes correlated with *KAT7* include genes within the HBO1 complex and the MLL1 complex. Similarly, highly correlated genes with *KMT2A* include genes within the MLL1 complex and the HBO1 complex. **B.** Genes included in the Lenti-U6^BC^sgRNA^Epistasis^/Cre library. **C.** Lenti-U6^BC^sgRNA^Epistasis^/Cre library is transduced intratracheally to 3 groups of mice to initiate lung tumors. The genotype and the number of mice for each group is indicated. **D.** The mean tumor size relative to inert sgRNAs for the genes targeted in the Lenti-U6^BC^sgRNA^Epistasis^/Cre library are shown for the *K;H11^LSL-Cas9^;Kat7^fl/fl^* mice (x-axis) and the *KT;H11^LSL-Cas9^* mice (y-axis). Each dot is a gene. The diagonal line (y=x) represents no change in relative mean tumor size between the two mouse genotypes. The error bars are 95% confidence intervals. **E.** Mean tumor size relative to inert sgRNAs for the HBO1^JADE2-ING5^ complex genes in *KT;H11^LSL-Cas9^* and *K;H11^LSL-Cas9^;Kat7^fl/fl^*mice. **F.** Mean tumor size relative to inert sgRNAs for the MLL complex genes in *KT;H11^LSL-Cas9^* and *K;H11^LSL-Cas9^;Kat7^fl/fl^* mice. **G.** Mean tumor size relative to inert sgRNAs for canonical tumor suppressor genes in *KT;H11^LSL-Cas9^* and *K;H11^LSL-Cas9^;Kat7^fl/fl^*mice. Genes are considered to have reduced tumor suppressive effects in *K;H11^LSL-Cas9^;Kat7^fl/fl^* mice if their 95% confidence intervals are outside the lower bound of the 95% confidence interval in *KT;H11^LSL-Cas9^* mice. Genes are considered to be no longer tumor suppressive if their relative mean size is ≤ 1 in *K;H11^LSL-Cas9^;Kat7^fl/fl^* mice but ≥ 1 in *KT;H11^LSL-Cas9^* mice (**see Methods**). **H.** The mean tumor size relative to inert sgRNAs when targeting genes in the HBO1^JADE2-ING5^ or MLL1 complex in the indicated mouse genotypes.

To dissect the functional relationship between the HBO1 and MLL1 complexes, and with other tumor suppressor genes, we generated a Tuba-seq^Ultra^ library targeting the MYST family associated genes, the COMPASS family associated genes and 20 canonical tumor-suppressive genes in *KT, KT;H11^LSL-Cas9^*, and *K;H11^LSL-Cas9^;Kat7^fl/fl^*mice (**Figure 7B-C, Supplemental Figure 14C-E, Supplemental Table 26**). As expected, Cre-mediated inactivation of *Kat7* increased tumor growth (**Supplemental Figure 14F-G**). Importantly, there were widespread epistatic interactions between the genes we targeted and *Kat7* (**Figure 7D**). Cas9-mediated targeting of *Kat7* increased tumor growth in *KT;H11^LSL-Cas9^*mice while having no effect in *K;H11^LSL-Cas9^*;*Kat7^fl/fl^* mice (**Figure 7E**). Likewise, inactivation of *Ing5, Meaf6,* or *Jade2* increased tumor growth in *KT;H11^LSL-Cas9^* mice but not in *K;H11^LSL-Cas9^; Kat7^fl/fl^* mice, demonstrating that these genes function as part of the tumor suppressive HBO1^JADE2-ING5^ complex. (**Figure 7E, Supplemental Table 27**).

To investigate the apparent genetic interaction between the HBO1 and MLL1 complexes in tumor suppression, we assessed the impact of coincident inactivation of *Kat7* and key members of the MLL1 complex. The increased tumor growth driven by *Kmt2a, Men1,* or *Psip1* inactivation was almost completely abrogated in the context of *Kat7* deficiency (**Figure 7F**). This dramatic reduction of tumor suppression by MLL1 complex genes in the absence of *Kat7* demonstrates the epistatic interaction between these two complexes, where the MLL1 and HBO1 complexes functionally interact to suppress lung tumor growth, consistent with their molecular similarities.

Since HBO1 has a broad impact on neoplastic cell state and the expression of canonical tumor suppressor genes (**Figure 6**), we hypothesized that inactivation of HBO1 could influence the functional outcome of other tumor suppressor pathways. We quantified the effects of inactivating 20 tumor suppressor genes on *Kat7*-proficient and *Kat7*-deficient lung tumorigenesis. Inactivation of several tumor suppressors, such as *Nf1*, *Rbm10*, and *Ezh2*, promoted tumor growth to a similar extent regardless of *Kat7* status. However, inactivating many others, such as *Pten*, *Stk11*, and *Setd2*, increased the growth of *Kat7-*deficient tumors less than *Kat7*-proficient tumors, indicating they are epistatic with *Kat7* (**Figure 7G**). Most dramatically, inactivating *Stag2* and other tumor suppressor genes associated with the STAG2-cohesin complex (Ashkin et al., 2024) increased the growth of *Kat7*-proficient tumors but decreased the growth of *Kat7*-deficient lung tumors, suggesting that STAG2-cohesin is required for *Kat7*-deficient lung tumors (**Figure 7G, Supplemental Figure 14H**) (Cai et al. 2021; Mayayo-Peralta et al. 2023). We further confirmed this synthetic lethal interaction by inactivating *Kat7* in *KT;H11^LSL-Cas9^;Stag2^fl/fl^* mice, which significantly reduced tumor size relative to *sgInert* tumors (**Supplemental Figure 15A-C**) (Ashkin et al., 2024). Thus, *Kat7* displays a range of epistatic interaction with a subset of tumor suppressor genes, further supporting its role as a pivotal regulator of lung tumorigenesis.

Finally, we assessed the generalizability of the HBO1 and MLL1 complex genes in suppressing lung tumorigenesis across different oncogenic contexts and in the absence of the critical p53 tumor suppressor gene (Blair et al. 2023; Tang et al. 2023a). We included sgRNAs targeting genes in the HBO1^JADE2-ING5^ complex and the MLL1 complex in a Tuba-seq^Ultra^ library and initiated lung tumors in *KT;H11^LSL-Cas9^*, *Braf^V600E^T;H11^LSL-Cas9^*, and *KT;p53^fl/fl^;H11^LSL-Cas9^* mice (**Methods,** and **Supplemental Figure 15D**). Inactivating these genes greatly increased tumor growth across all genetic backgrounds (**Figure 7H, Supplemental Figure 16, Supplemental Table 28**). Thus, tumor suppression by the HBO1 and MLL1 complex extends broadly across multiple genetic driver contexts in lung tumors.

## DISCUSSION

Enabled by the simplicity and versatility of Tuba-seq^Ultra^, we performed iterative functional screens *in vivo* to systematically investigate epigenetic regulatory genes that control lung tumorigenesis. This experimental paradigm of discovery, saturation, and epistasis screens within autochthonous cancer models, coupled with molecular analysis identified previously unappreciated HBO1^JADE2-ING5^ and MLL1 complexes as among the most impactful tumor suppressive epigenetic regulators in lung tumorigenesis.

An important implication of this work is the capacity to move beyond studying individual driver genes to investigating all functional units such as protein complexes that can impact tumorigenesis. Epigenetic regulatory genes can affect a wide range of cellular functions and participate in multiple complexes. By examining their functional impacts in parallel, we deconvoluted the myriad potential functions and pathways associated with each gene and pinpointed the specific molecular machinery important in tumorigenesis. Through targeting all the genes in the larger MYST family and paralogs within the HBO1 complex, we clearly delineated the HBO1^JADE2-ING5^ complex as the specific tumor suppressive complex.

The HBO1 and MLL1 complexes maintain chromatin accessibility at regions enriched for transcription factor motifs involved in lung differentiation, and inactivation of these complexes promotes a shift toward cell states of lineages from different tissues. Increased cellular plasticity and lineage infidelity are frequently observed during tumor progression. Chromatin remodeling is fundamental to the emergence of cellular plasticity and lineage infidelity (Marjanovic et al. 2020; Kaiser et al. 2023). Cells with enhanced plasticity and lineage infidelity play critical roles in promoting many aspects of carcinogenesis from therapy resistance to metastatic ability (LaFave et al. 2020; Torborg et al. 2022) In lung and pancreatic cancer, cells with enhanced plasticity are defined by specific chromatin accessibility modules (LaFave et al. 2020; Burdziak et al. 2023). Thus, the ability of the HBO1 and MLL1 complexes to safeguard lineage fidelity and prevent dedifferentiation places them as an essential nexus of tumor suppression.

Our work demonstrates the crucial role of non-mutational mechanisms in determining tumor phenotypes. Not only are the HBO1 and MLL1 complexes crucial tumor suppressive epigenetic regulators, but the disruption of the HBO1 complex in human tumors likely occurs primarily through non-mutational mechanisms. The relatively low mutational rates for individual genes in the HBO1 or MLL1 complex may be partially attributed to mutually exclusive mutations in the genes in these interrelated complexes. Notably, more than half of human lung adenocarcinomas have absent or diminished levels of H3K14ac. Thus, HBO1 activity in most of these tumors is reduced through mutation-independent mechanisms.

We anticipate that the relationship between the HBO1 and MLL1 complexes in cancer is multifaceted. Biochemical analyses have shown that the HBO1 complex can bind the MLL1 complex (Gaurav et al. 2024). Additionally, the functional interaction between the HBO1 and MLL1 complexes could be mediated through cross recruitment to their histone modifications. The PHD fingers in ING4/5 bind to H3K4me3 generated by the MLL1 complex (Hung et al. 2009), thereby recruiting the HBO1 complex. Conversely, the MLL1 complex can interact with H3K14ac generated by the HBO1 complex leading to the generation of H3K4me3 (Nakanishi et al. 2008).

The interaction between the HBO1 and MLL1 complex is highly conserved and critical to their function in the cancer context. There is broad fitness correlation between the HBO1 and MLL1 complexes across diverse human cancer cell lines, particularly for the HBO1 complex containing JADE2. However, the functional output of this interaction is linked to cellular and genetic context. In MLL1 fusion-driven leukemia, the HBO1 and MLL1 complex interaction is essential for cell survival, and KAT7 has been suggested as a potential therapeutic target (MacPherson et al. 2020). Moreover, fusions containing KAT7 or JADE2 potently transforms hematopoietic progenitors into leukemic cells while other JADEs fusion has limited transforming capacity (Gaurav et al. 2024). These data demonstrate the unique role of the HBO1^JADE2-ING5^ complex in regulating cancer phenotypes. In contrast, the HBO1 and MLL1 complexes interact to suppress lung tumorigenesis in multiple oncogenic backgrounds. *Kat7* inactivation changes the impact of several canonical tumor suppressors, suggesting that the HBO1 complex interfaces with multiple tumor-suppressive pathways. Interestingly, *Kat7* is a genetic vulnerability in *Stag2*-deficient lung tumors, similar to its role in leukemia. These context-dependent functional outputs underscore the importance of using diverse *in vivo* models to investigate the impact of cancer type and genotype on epigenetic regulation in cancer. Furthermore, the success of therapeutic targeting of KAT7 will require thorough consideration of both the tumor type and mutational landscape.

Although this study primarily focused on tumor-suppressive epigenetic regulators, we identified multiple genes that are required for autochthonous lung tumor growth. Inactivation of several genes reduced tumor growth to an extent comparable to known drug targets including SOS2 and PTPN11/SHP2. Inhibitors already exist for some of the novel epigenetic tumor dependency genes that we identified, such as *Kdm1a* (Fang et al. 2019) and *Kat6a* (Mukohara et al. 2024). Future *in vivo* studies on these epigenetic tumor dependency genes, considering cellular and genetic contexts, may uncover and validate novel druggable targets for lung cancer.

Tuba-seq^Ultra^ is cost-effective, highly accessible and has virtually unlimited scalability compared to previous methods for barcode-based CRISPR screens with clonal tracing. We anticipate that Tuba-seq^Ultra^ will enable rapid and comprehensive *in vivo* functional genomics analyses, including studies that focus on broad biological processes, identifying therapeutic targets, and dissecting the intricate genetic mechanisms involved in a wide range of biological questions and disease models.

## METHODS

### Experimental Model and Subject Details

#### Mouse models

All animal experiments in this study were approved by the Institutional Animal Care and Use Committee (IACUC) at Stanford University, protocol number 26696. *Kras^LSL-G12D/+^* (RRID:IMSR_JAX:008179) (Jackson et al. 2001), *Trp53^fl/fl^* (RRID:IMSR_JAX008462) (Marino et al. 2000), *R26^LSL-tdTomato^* (RRID:IMSR_JAX:007914) (Madisen et al. 2010), and *H11^LSL-Cas9^* (RRID:IMSR_JAX:027632) (Chiou et al. 2015), *Kat7^fl/fl^* (Kueh et al. 2011), *Braf^V600E^*(RRID:IMSR_JAX: 017837) (Dankort et al. 2007) mice have been previously described. All mice were on a C57BL/6 background and kept in standard condition with 12hour light/dark cycle and ad libitum access to food and water.

### Human lung adenocarcinoma tissue microarray

Human patient samples were obtained from Sandford University School of Medicine Department of Pathology. All clinical specimens in this study were collected from patients with written informed consent for research use in accordance with the Declaration of Helsinki and approved by the Stanford University Institutional Review Board. In this study we used formalin fixed paraffin embedded (FFPE) lung cancer tissue microarray containing up to n = 220 lung adenocarcinoma samples, n = 34 mixed bronchioalveolar carcinoma samples, and n = 16 normal lung samples. Other cancer and tissue samples on the slide and samples that failed to stain due to TMA preparation were excluded from this study.

## METHOD DETAILS

### Tuba-seq^Ultra^ vector design and barcoding

Proper functioning of the U6 promoter requires sequence motifs including the octamer, proximal sequence element, distal sequence element, and the TATA box. Previous work has shown that the U6 promoter can be engineered if these critical motifs and the distance between them are intact (Ventura et al. 2004). Thus, we hypothesized that the regions immediately after the TATA box is amendable for barcode incorporation. To generate the Tuba-seq^Ultra^ vector, we replaced the human U6 promoter in the previously described Lenti-sgRNA/Cre vector (Cai et al. 2021) with a gBlock (Integrated DNA Technologies, IA) synthesized bovine U6 (bU6) promoter via Gibson assembly. The bU6 promoter allows effective sgRNA expression (Adamson et al. 2016) and expands the potential barcode space from 15 nucleotides in hU6 to 23 nucleotides, providing up to 4^23^ barcode diversity (Lambeth et al. 2005). In our design, the last 3 nucleotides in the bU6 vector were left unchanged to serve as sticky ends to clone sgRNAs and we included fixed nucleotides in the barcode region to distinguish between different Tuba-seq^Ultra^ libraries.

To barcode the Tuba-seq^Ultra^ vector, it was linearized by PCR using high fidelity Q5 polymerase (NEB, MO0492L) with primers flanking the U6-barcode-sgRNA region.

Forward primer: 5’ GTTTAAGAGCTAAGCTGGAAACA 3’.

Reverse primer: 5’ AACTATATAAAGCTAAGAATAGAAAAAATTTTACAGTTATGGT 3’.

The PCR product was then treated with DpnI restriction endonuclease (NEB, R0176S) to digest the original vector thereby reducing background bacteria colonies. This linearized backbone was purified using Qiagen PCR purification with QIAquick Mini Spin columns following manufacturer’s protocol (Qiagen, #28106). Barcoding primers were synthesized by Stanford Protein and Nucleic Acid Facility (Stanford, CA) as a ∼100 nucleotide single stranded sequence with the following format: actgtaaaattttttctattcttagctttatatagttNNNNNNNNNNNNNNNNNatggaccaggatgggcaccacccgtttaagagctaagctggaaac

The Ns in the primer denote the degenerate sequences that serve as barcodes. This primer consists of homology regions with the Tuba-seq^Ultra^ vector backbone (lower case sequences) and two Esp3I sites in opposite orientation for library insertion. Gibson assembly was performed to anneal the barcoding primer with the purified linearized Tuba-seq^Ultra^ vector using NEBuilder HiFi DNA Assembly Master Mix (NEB, ES2621L) with 150ng/µL of vector and 1µL of 10uM barcoding primer per 20µL reaction at 50 °C for 4 hours. The assembled product was purified with standard isopropanol precipitation protocol and re-suspended in TE. The resuspended barcoded vector was electroporated into 10-beta electrocompetent *E. coli* (NEB, C3020K) following manufacturer’s protocol. The bacteria colonies are pooled, and the plasmids were extracted using Qiagen Midi-prep kit (Qiagen, #12941). Depending on the library size ∼ 150,000 – 2,000,000 colonies were used per library.

### sgRNA design, synthesis and cloning of Lenti-Tuba-seq^Ultra^ library

sgRNAs in the Lenti-U6^BC^sgRNA^Epigenetics^/Cre and Lenti-U6^BC^sgRNA^Saturation^/Cre library were primarily taken from the Brie mouse CRISPR libraries (Doench et al. 2016). The sequences were accessed from Addgene (https://www.addgene.org/pooled-library/broadgpp-mouse-knockout-brie/). For each gene, we selected the first three sgRNAs, which typically had the highest Rule 2 scores (Doench et al. 2016). Non-targeting sgRNAs (*sgNT*) were taken from Cai *et al*., 2021. Safe targeting sgRNAs (*sgSafes*) were taken from Bassik Mouse CRISPR Knockout Library accessed from Addgene (https://www.addgene.org/pooled-library/bassik-mouse-crispr-knockout/) (Morgens et al. 2017). For complete sequence of sgRNAs in each library, see Supplemental Tables 2, 8, and 27.

The sgRNAs for each library were synthesized on oligo chips (Agilent Technologies, CA, and Twist Biosciences, CA) in the format TAAGGTGCGTACTAGCTGACCGTCTCTatggXXXXXXXXXXXXXXXXXXXXgtttAGAG ACGGCGCCTACACACCTATCTAA Upper case sequences denote PCR handles for amplifying the oligo libraries, lower case sequence denote Esp3I restriction site overhangs and Xs denote sgRNA sequences. The synthesized sgRNA library is PCR amplified with Q5 polymerase (NEB, MO0492L) for 11 – 12 cycles using the following primers.

Forward primer: 5’ TAAGGTGCGTACTAGCTGAC 3’.

Reverse primer: 5’ TTAGATAGGTGTGTAGGCGC 3’. The amplified sgRNA library is purified with QIAquick Mini Spin columns (Qiagen, #28106) following manufacturer’s protocol.

The barcoded Tuba-seq^Ultra^ vector was digested with BsmBI-v2 (NEB R0379L) for 2 hours at 55°C and purified with QIAquick Mini Spin columns (Qiagen, #28106) following the manufacturer’s protocol. Golden Gate Assembly was used to ligate the PCR amplified sgRNA library with the barcoded Tuba-seq^Ultra^ vector. The molar ratio of insert (sgRNA library) to backbone (barcoded vector) was approximately 2:1. The Golden Gate assembly mix included 10X T4 Ligase buffer (NEB, B0202S), T4 DNA ligase (NEB, M0202S), Esp3I (NEB, R0734S), and molecular grade water (Cytiva, SH30538). The reaction was set up overnight for 70 cycles (37°C, 5 min → 20°C, 5 min). The assembled product was purified with isopropanol precipitation following standard protocol and electroporated into 10-beta electrocompetent *E. coli* (NEB, C3020K) as described by the manufacturer. The bacterial colonies were pooled (up to 10 million colonies for a library) and the plasmids were extracted via Qiagen Midi-prep (Qiagen, #12941) with endotoxin removal.

To check library integrity, barcode diversity and sgRNA representation, the U6-barcode-sgRNA region of the plasmid library was PCR amplified using primers compatible with Illumina sequencing (Illumina, CA). Forward primer:

AATGATACGGCGACCACCGAGATCTACACxxxxxxxxxACACTCTTTCCCTACACGACG CTCTTCCGATCTNNNNNNNAGGGTTACAGTTTAGTCACCATA

Reverse primer: CAAGCAGAAGACGGCATACGAGxxxxxxxxGTGACTGGAGTTCAGACGTGTGCTCTTC CGATCTNNNNNNNNCGACTCGGTGCCACTTTTTC

Ns denote degenerate nucleotides to increase read diversity at the beginning of the amplicon, which facilitate cluster identification. Xs denote the i5 and i7 indexes. The PCR amplified plasmid library was validated by deep sequencing on MiSeq-Nano (Illumina, CA) or Novaseq6000 (Illumina, CA).

### Single sgRNA cloning

The single sgRNAs used for initiating *sgKat7*, *sgMeaf6*, *sgPsip1* and *sgKmt2a* tumors were selected from the Lenti-U6^BC^sgRNA^Epigenetics^/Cre library, using the sgRNA with the largest effect size. Their sequences were:

*sgKat7*: gCCTATCCTGAGGAATATGCG

*sgMeaf6*: gCCAATAGCAAAAACGACCGG

*sgPsip1*: gAGAGAGGACGACCTGCAGGT

*sgKmt2a*: GCAGCCGTTAGACCTCGAAG

*sgSafe14*: gCAAAATTGCATTCCCTCGTT

*sgSafe23*: gTATACCATTCCAGAATGTCG

These sgRNAs were individually cloned into the unbarcoded Lenti-Tuba-seq^Ultra^ vector by site-directed mutagenesis (SDM). The primers were designed using NEBaseChanger (https://nebasechanger.neb.com/). The unbarcoded vector was amplified with SDM primers using Q5 polymerase (NEB, M0492L). The amplified products were phosphorylated and circularized with KLD enzyme mix (NEB, M0554S). The KLD-treated products were chemically transformed into chemically competent *E. coli* (NEB, C2987H) following the manufacturer’s protocol. Individual colonies were selected and inoculated in LB broth with ampicillin (100 mg/L). The plasmids were extracted using the QIAprep Spin Miniprep Kit (Qiagen, 27106) and verified by Sanger sequencing (Elim Biopharm, CA). Correct colonies were expanded again in LB broth with ampicillin (100 mg/L) and the plasmids were extracted using the Qiagen Midi-prep kit (Qiagen, 12941) with endotoxin removal.

### Lentivirus packaging

Each Lenti-U6^BC^sgRNA/Cre library was co-transfected into HEK293T cells with pCMV601 VSV-G (Addgene, 8454) envelope plasmid and pCMV-dR8.2 dvpr (Addgene, 8455) packaging plasmid using Opti-MEM (Gibco, 31985070) in 150 mm cell culture plates. The HEK293T cells were cultured in DMEM (Gibco, 11-960-051) with 10% FBS (Phoenix Scientific, PS-100-00-500), 1% Pen/Strep/glutamine (Invitrogen, 10378-016), and 0.1% amphotericin B (Life Technologies, 15290-018). Sodium butyrate (Sigma-Aldrich, B5887) was added 8 h after transfection to achieve a final concentration of 20 mM. Cell culture media was refreshed 24 h after transfection. Supernatants were collected at 36 and 48 hours after transfection, filtered through a 0.45 μm syringe filter (Millipore, SLHP033RB) to remove cells and debris, concentrated by ultracentrifugation (25,000g for 1.5 hours at 4°C), and resuspended overnight in PBS and stored at −80°C until use.

Lentiviral packaging for single Lenti-sgRNA/Cre vectors was performed in a similar manner, except the vectors were transfected individually into separate plates of HEK293T cells.

To determine the titer of the concentrated lentiviral particles, we transduced LSL-YFP cells (a gift from Dr. Alejandro Sweet-Cordero/UCSF) with the virus and measured the percent YFP-positive cells by flow cytometry. The titer was calculated by comparing the percent YFP-positive cells to a lentiviral preparation of known titer.

### Dual-sgRNA barcoding and cloning

Cloning and barcoding of dual-sgRNA was performed as described elsewhere (Hebert et al. 2024a). Briefly, the Tuba-seq^Ultra^ vector was first barcoded as described above. The sgRNAs were ordered as a single-stranded DNA pool (Twist Biosciences, CA) containing two spacers (guides) flanked by BsmBI Type IIS restriction sites. The oligo sequences were amplified with low cycle PCR and ligated into a donor sequence containing one tracrRNA and the mouse U6 promoter by Golden Gate Assembly. The ligated product was then cloned into the barcoded Tuba-seq^Ultra^ vector by Golden Gate Assembly.

### Spike-in construction and cell line generation

Three Lenti-U6^BC^sgSpike-in/Cre vectors were generated to create the spike-in cell lines that were used as benchmarks to calculate cell number during Tuba-seq^Ultra^ analysis (**see Tuba-seq^Ultra^ Analysis**). Each vector contained non-targeting sgRNAs. The spike-in vectors were cloned by Gibson assembly following the same method as used for barcoding the Tuba-seqUltra vector described above (**see Tuba-seq^Ultra^ Vector Design and Barcoding**). The primer sequences are as follows:

Spike-in 1

ACTGTAAAATTTTTTCTATTCTTAGCTTTATATAGTTCNNNNNNNNCNNNNNNTATGggagttctgcctcaagcaagtGTTTAAGAGCTAAGCTGGAAAC

Spike-in 2

ACTGTAAAATTTTTTCTATTCTTAGCTTTATATAGTTCNNNNNNNNCNNNNNNTATGggttgaatcgtccgtacatgtGTTTAAGAGCTAAGCTGGAAAC

Spike-in 3

CTGTAAAATTTTTTCTATTCTTAGCTTTATATAGTTCNNNNNNNNCNNNNNNTATG gtatacttgcaccatgccataGTTTAAGAGCTAAGCTGGAAAC

The cloned vectors were electroporated into 10-beta electrocompetent *E. coli* (NEB, C3020K) and the bacteria colonies were pooled. The plasmids were extracted via Qiagen Midi-prep (Qiagen, 12941) with endotoxin removal and the U6^BC^ -sgRNA region was verified by Sanger sequencing (Elim Biopharm, CA).

The Lenti-U6^BC^sgSpike-in/Cre vectors were packaged individually into lentivirus by co-transfecting with pCMV601 VSV-G and pCMV-dR8.2 dvpr plasmid into HEK293T cells as described above. The viruses were collected at 36 hours after transfection and added to separate plates of LSL-YFP cells at low titer to reduce multiple transductions to generate the three spike-in cell lines. The transduced cells were sorted by FACS and the percent YFP positive cell for each spike-in cell line was less than 1% of the total population. The sorted YFP positive cells were expanded *in vitro* and aliquoted into tubes containing either 50,000 or 100,000 cells from each spike-in line per vial. The aliquoted spike-in vials were stored at −80 °C until use.

### Tumor initiation and monitoring

Lung tumors were initiated by intratracheal intubation of isoflurane-anesthetized mice. Animal cohorts aged between 8 to 16 weeks with a balanced sex ratio were transduced. Each mouse received 60 μL of virus in PBS. For the initial cohort of Lenti-U6^BC^sgRNA^Epigenetics^/Cre library, the *KT;H11^LSL-Cas9^* mice (n=26/66) received varying titers of virus to determine the most optimal titer for transduction. Specifically, n=8 mice received 50,000 ifu per mouse, n=6 mice received 150,000 ifu per mouse, and n=8 mice received 300,000 ifu per mouse. After this experiment, the remaining 40 *KT;H11^LSL-Cas9^* mice in the second cohort each received 150,000 ifu. Since analyses were performed at the level of individual tumors relative to inert sgRNAs in the same animals, the mice transduced with different titers were pooled together during analyses (**see Tuba-seq^Ultra^ analysis**). For the Lenti-U6^BC^sgRNA^Saturation^/Cre library, a single cohort of *KT;H11^LSL-Cas9^* and *KT* mice were transduced with 150,000 ifu per mouse and 300,000 ifu per mouse, respectively. For the Lenti-U6^BC^sgRNA^Epistasis^/Cre library, the *K;H11^LSL-Cas9^;Kat7^fl/fl^* mice were transduced with 150,000 ifu per mouse, the *KT;H11^LSL-Cas9^* mice were transduced with 200,000 ifu per mouse and the *KT* mice were transduced with 400,000 ifu per mouse. Note that in the *K;H11^LSL-Cas9^;Kat7^fl/fl^* mice, 5/9 mice carried the *R26^LSL-tdTomato^* allele, whereas 4/9 mice were wildtype. Animals transduced with Tuba-seq^Ultra^ libraries were euthanized after 15 weeks and the lungs were collected.

For initiating single gene inactivated lung tumors, mice transduced with Lenti-sgMeaf6/Cre, Lenti-sgKmt2a/Cre, Lenti-sgPsip1/Cre, Lenti-sgSafe14/Cre and Lenti-sgSafe23/Cre vectors at 200,000 ifu per mouse regardless of mouse genotype. The number of animals for each experiment was indicated in the figures. For transduction with the Lenti-sgKat7/Cre vector, the *KT;H11^LSL-Cas9^* (n = 10) and *KT* (n = 4) mice received 200,000 ifu per mouse and a separate cohort of *KT;H11^LSL-Cas9^* (n = 3) and *KT* (n = 3) mice received 120,000 ifu per mouse. The animals were monitored regularly and euthanized when they displayed morbidity signs including weight loss, heavy breathing, and reduced mobility.

For assessing tumor grades in *KT* and *KT;H11^LSL-Cas9^;Kat7^fl/fl^*mice, the tumors were initiated intratracheally with Lenti-sgSafe23/Cre vector at 200,000 ifu per mouse. The mice were harvested 27 weeks post viral transduction, and the lungs were collected.

### Tissue processing

For all experiments, the lungs were weighed at the time of collection. Lungs from mice transduced with Tuba-seq^Ultra^ libraries were frozen at −20 °C for genomic DNA extraction. Lungs transduced with single Lenti-sgRNA/Cre vectors were harvested, and one lobe dissected and fixed in 10% formalin for histology. The remaining lobes were immediately minced into fine pieces and enzymatically dissociated with solution containing collagenase IV, dispase, and trypsin at 37 °C for 30 minutes following a previously described method (Chuang et al. 2017). The single cell suspensions were frozen in Bambanker cryopreservation media (Wako Chemicals, 302-14681) and stored in liquid nitrogen.

### Tuba-seq^Ultra^ library preparation

The aliquoted benchmark spike-in cell lines were added to the frozen whole lung tissue from each mouse. Thus, each frozen lung tissue had three independent benchmark spike-in cell lines. For experiments using Lenti-U6^BC^sgRNA^Epigenetics^/Cre and Lenti-U6^BC^sgRNA^Saturation^/Cre libraries, 100,000 cells from each of the three spike-in cell lines were added to the lungs for a total of 300,0000 spike-in cells per mouse. For experiments involving the Lenti-U6^BC^sgRNA^Epistasis^/Cre library, 50,000 cells from each of the three spike-in cell lines were added to the lungs, for a total of 150,000 spike-in cells per mouse. Genomic DNA from whole tumor bearing lungs with the spike-in cells added were extracted as previously described (Rogers et al. 2017). Briefly, the tissues were homogenized in 20 ml lysis buffer (100mM NaCl, 20mM Tris, 10mM EDTA, 0.5% SDS) with 200 μl of 20 mg/ml Proteinase K (Life Technologies, AM2544). Homogenized tissues were incubated at 55°C overnight. Genomic DNA was extracted with phenol-chloroform (ThermoFisher Scientific, 15593049) or the Qiagen Puregene Kit (Qiagen, 158063) and precipitated with ethanol.

Next generation sequencing libraries for experiments involving the Lenti-U6^BC^sgRNA^Epigenetics^/Cre library were prepared by amplifying the U6-barcode-sgRNA in a two-step nested PCR with NEBNext Ultra II Q5 Master Mix (NEB, M0544S). For each mouse, 32μg of genomic DNA was first amplified with primers (forward: 5’ TCGATTAGTGAACGGATCGGC 3’, reverse: 5’ CGAACCTCATCACTCGTTGC 3’) for 15 cycles to enrich the U6-barcode-sgRNA region. The first PCR products were then amplified with Illumina sequencing compatible primers in the format described above for plasmid sequencing with unique dual i5 and i7 indexes for each sample (**see sgRNA design, synthesis and cloning of Lenti-Tuba-seq^Ultra^ library**) for 15 cycles using NEBNext Ultra II Q5 Master Mix. Library preparation for experiments involving the Lenti-U6^BC^sgRNA^Saturation^/Cre and Lenti-U6^BC^sgRNA^Epistasis^/Cre vectors, 32μg of genomic DNA from each mouse was amplified in a single step using the Illumina sequencing compatible primers for up to 35 PCR cycles. The final PCR products underwent double-sided purification (0.70X left side and 0.95X right side) using SPRIselect beads (BD Life Sciences, B23317) and were pooled. The purified libraries were quantified and checked for quality using Agilent TapeStation system (Agilent, CA).

### Deep sequencing

Unequal and/or insufficient sequencing depth between lung samples can introduce noise in the data in computing tumor size and number. To ensure all lung samples were sequenced with adequate depth, the Lenti-U6^BC^sgRNA^Epigenetics^/Cre library was sequenced in a two-step process. First, the deep sequencing library for each lung sample was pooled evenly and was sequenced using 150bp pair-end reads on Illumina MiSeq Nano (Illumina, CA). Since a fixed number of benchmark spike-in cells were present in each lung sample, the relative sequencing depth for each sample was determined based on the number of reads for each spike-in cell benchmark in the sample. The samples were then re-pooled by adjusting the quantity of the prepared library for that sample based on the relative spike-in counts. The re-pooled library was sequenced with Illumina NextSeq 500 (performed by Admera Health, NJ) or Illumina Novaseq 6000 (performed by Novogene, CA). For the Lenti-U6^BC^sgRNA^Saturation^/Cre and Lenti-U6^BC^sgRNA^Epistatsis^/Cre libraries, individual samples were pooled based on lung weights, a proxy for total tumor burden. All samples were sequenced with 150bp pair-end sequencing to a minimum depth of 2 reads per 150 cells.

### Immunohistochemistry

Lung tissues fixed in 10% formalin were embedded in paraffin and sectioned by Stanford Pathology/Histology Service Center (Stanford, CA) or Histo-Tec Laboratories (Hayward, CA). Hematoxylin and eosin stains were performed by Stanford Pathology/Histology Service Center (Stanford, CA) or Histo-Tec Laboratories (Hayward, CA). Immunohistochemistry was performed on 4μm sections. Antigen retrieval was performed with 10mM citrate buffer in a pressurized decloaking chamber. The slides were washed with 1X PBST and prepared with VECTASTAINABC-HRP Kit (Vector Laboratories, PK-4000). Antibody dilutions were 1: 250 for NKX2.1 (Abcam, ab76013, RRID:AB_1310784), 1: 100 for p63 (Cell Signaling, 13109S, RRID:AB_2637091), 1: 200 for H3K14ac (Abcam, ab52946, RRID:AB_880442), 1: 1000 for H4K12ac (Abcam, ab177793, RRID:AB_2651187), and 1: 200 for H4K5ac (Abcam, ab51997, RRID:AB_2264109). Sections were developed with DAB (Vector Laboratories, SK-4100) and counterstained with hematoxylin.

Immunohistochemistry for human lung adenocarcinoma TMAs was performed by Stanford Pathology/Histology Service Center. The dilutions were 1: 100 for H3K14ac (Abcam, ab52946, RRID:AB_880442), 1: 200 for H4K12ac (Abcam, ab177793, RRID:AB_2651187), 1: 200 for H4K5ac (Abcam, ab51997, RRID:AB_2264109), and 1:100 for H3K36me3 (Cell Signaling, 4909, RRID:AB_1950412).

### FACS of neoplastic cells

Dissociated tumor cells in single cell suspension were retrieved from liquid nitrogen storage and thawed at 37°C. The cells were washed in 1X PBS and stained with antibodies targeting CD45 (BioLegend, 103112), CD31 (BioLegend, 303116), F4/80 (BioLegend, 123116) and Ter119 (BioLegend, 116212) at 1:800 dilution for each antibody. DAPI (BD Biosciences, 564907) was used as live/dead indicator. The cells were sorted by flow cytometry on FACSymphony S6 (BD Biosciences) or FACSAria II (BD Biosciences) for CD45-negative, CD31-negative, F4/80-negative, Ter119-negative cells and tdTomato-positive (Lin^−^ TdTom^+^) live neoplastic cells.

### Western blotting

FACS-isolated cells neoplastic cells were lysed by RIPA buffer (Fisher Scientific, PI89900) supplemented with protease inhibitor cocktail (Millipore, 11836170001). Denatured samples were run on a 4%–12% Bis-Tris gradient gel (Thermo Scientific, NP0323BOX) and transferred onto PVDF membranes. Membranes were immunoblotted using primary antibodies at 1: 1000 for H3K14ac (Abcam, ab52946, RRID:AB_880442), 1: 2000 for H4K12ac (Abcam, ab177793, RRID:AB_2651187), and 1: 2000 for H4K5ac (Abcam, ab51997, RRID:AB_2264109), 1:750 for β-ACTIN (Cell Signaling, 4970, RRID:AB_10691808), 1:1000 for KAT7 (Proteintech, 13751-1-AP, RRID:AB_2266703), and 1:2000 for Histone 3 (Abcam, ab1791, RRID:AB_302613). Histone 3 staining was performed on membranes stripped with Restore PLUS Western Blot Stripping Buffer (ThermoFisher, 46430). HRP-conjugated goat-anti-rabbit antibody was used as the secondary antibody at 1:5000 dilution (Santa Cruz Biotechnology, sc-2004).

### Bulk ATAC-seq library preparation

Two separate ATAC-seq experiments were performed following the Omni-ATAC protocol as previously described (Corces et al. 2017; Marinov et al. 2023b; Kim et al. 2024). One experiment was on FACS sorted Lin^−^TdTomato^+^ neoplastic cells from *KT;H11^LSL-Cas9^* mice transduced with *sgKat7* (n = 4) or *sgSafe23* (n = 4). The other ATAC-seq experiment was on FACS sorted Lin^−^TdTomato^+^ neoplastic cells from *KT;H11^LSL-Cas9^* mice transduced with *sgMeaf6* (n = 3), *sgKmt2a* (n = 3), *sgPsip1* (n = 3), *sgSafe23* (n = 3), or *sgSafe14* (n = 1)). Briefly, approximately 50,000 cells were washed once in 1*×* PBS and resuspended in 50 µL ATAC-RSB-Lysis buffer (10 mM Tris-HCl pH 7.4, 10 mM NaCl, 3 mM MgCl_2_, 0.1% IGEPAL CA-630, 0.1% Tween-20, 0.01% Digitonin). The cells in lysis buffer were incubated on ice for 3 minutes. Then 1 mL ATAC-RSB-Wash buffer (10 mM Tris-HCl pH 7.4, 10 mM NaCl, 3 mM MgCl_2_, 0.1% Tween20, 0.01% Digitonin) was added, and the nuclei were centrifuged at 500 *g* for 5 min at 4 °C. To transpose the nuclei, a solution of 25 µL 2*×* TD buffer (20 mM Tris-HCl pH 7.6, 10 mM MgCl_2_, 20% Dimethyl Formamide), 2.5 µL Tn5 transposase (custom produced) and 22.5 *µ*L nuclease-free H_2_O was added, followed by incubation for 30 minutes at 37 °C while agitating at 1000 RPM. The transposed DNA was isolated using the MinElute PCR Purification Kit (Qiagen, 28004/28006), and PCR amplified as previously described (72 °C for 3 minutes, 98 °C for 30 seconds, 10 cycles of 98 °C for 10 seconds, 63 °C for 30 seconds, 72 °C for 30 seconds) (Corces et al. 2017; Marinov et al. 2023b; Kim et al. 2024). Libraries were purified using the Qiagen MinElute PCR Purification kit (Qiagen, 28004), then sequenced on an Illumina NextSeq 550 as 2*×*38mers (Illumina, CA).

### Single-cell RNA-seq library preparation

Two separate scRNA-seq experiments were performed using the split-seq approach from Parse Biosciences (Rosenberg et al. 2018; Tran et al. 2022). One experiment was on FACS sorted Lin^−^TdTomato^+^ neoplastic cells from *KT;H11^LSL-Cas9^*mice transduced with *sgKat7* (n = 3), *sgSafe23* (n = 2), or *sgSafe14* (n = 1). The other scRNA-seq experiment was on FACS sorted Lin^−^ TdTomato^+^ neoplastic cells from *KT;H11^LSL-Cas9^* mice transduced with *sgMeaf6* (n = 3), *sgKmt2a* (n = 3), *sgPsip1* (n = 3), or *sgSafe14* (n = 4). Immediately after FACS, the sorted neoplastic cells were fixed with Evercode Cell Fixation kit (Parse Biosciences, ECF2101). For the *sgKat7* versus *sgSafe14* experiment, the fixed cells were prepared using Evercode WT Mini v2 kit (Parse Biosciences, ECW02110) according to the manufacturer’s instructions. For the other scRNA-seq experiment, the fixed cells were prepared using Evercode WT v2 kit (Parse Biosciences, ECW02030) according to manufacturer’s instructions. The Parse Mini v2 kit generated two sub-libraries and the Parse WT v2 kit generated 8 sub-libraries. The sub-libraries were inspected by the Agilent TapeStation system (Agilent, CA) and sequenced by Illumina NovaSeq S4 flow cell.

## QUANTIFICATION and STATISTICAL ANALYSIS

### Tuba-seq^Ultra^ analysis

#### A. Raw reads parsing and cleaning

Paired-end reads were first merged using AdapterRemoval (Schubert et al. 2016), and were parsed using regular expressions to identify the sgRNA sequence and U6 barcodes. For identifying sgRNA sequences, we required a complete match with the designed sequences, as mutated sgRNA could suffer from reduced efficiency and off-target effects. For different libraries involved in the study, we designed specific barcode patterns to distinguish each library, thereby allowing us to identify any cross-library contamination. Depending on the specific library, the clonal barcode consists of either a 14-nucleotide (Lenti-U6^BC^sgRNA^Epigenetics^ library) or a 12-nucleotide (Lenti-U6^BC^sgRNA^Epistasis^ and Lenti-U6^BC^sgRNA^Saturation^ libraries) random barcode. This high theoretical diversity of clonal barcode (> 4^12^) relative to the total number of tumors ensured that each clonal tumor is uniquely barcoded. Given the throughput of our method, which targets hundreds of sgRNA in parallel in the same animal, usually less than 100 tumors were generated for each sgRNA per mouse. Hence, the likelihood of two clonal tumors in the same animal carrying a barcode-sgRNA sequence within 1-hamming distance is exceedingly low. Therefore, when we encountered low-frequency clonal barcodes within a 1-hamming distance of high-frequency clonal barcodes, we attributed them to sequencing or PCR errors (spurious reads) (Li et al. 2021). These low-frequency barcodes within 1-hamming distance were merged with barcodes of higher frequencies. Finally, total reads for each barcode-sgRNA combination were calculated.

#### B. Clonal tumor size estimation

To enable the conversion of read counts associated with clonal tumors into absolute neoplastic cell numbers, we added three “spike-in” cell lines with defined cell numbers to each sample prior to lung lysis and DNA extraction (**see Spike-in generation**). This conversion was accomplished by normalizing the read counts of the tumor cells to the reads of the “spike-in” cells. For each sample, we checked the read ratio among the three “spike-in” cell lines. Occasionally, if one “spike-in” cell line was under-amplified or over-sampled compared to the other two, we only used the other two to calculate the cell number conversion. Otherwise, all three “spike-in” cell lines were used for the conversion.

To ensure our data were derived only from high-quality samples, any animals with fewer than 10^5^ reads and/or that had more than 50% of the reads come from “spike-in” cells were discarded. We also checked the effect of the positive control and negative control sgRNA in each sample and discarded samples with apparent mislabeled mouse genotypes. Three mice were removed from the Lenti-U6^BC^sgRNA^Epigenetics^/Cre library analysis, 4 mice were removed from the Lenti-U6^BC^sgRNA^Epigenetics^/Cre library analysis, and 2 mice were removed from the Lenti-U6^BC^sgRNA^Epistasis^/Cre library analysis.

The median sequencing depth across all experiments was approximately between 24 - 123 cells per read. To perform statistical comparisons of tumor genotypes, we imposed a minimum tumor size cutoff of 300 or 800 cells, depending on the corresponding sequencing depth of the experiments. Specifically, the tumor size cutoff for the Lenti-U6^BC^sgRNA^Epigenetics^/Cre library and the Lenti-U6^BC^sgRNA^Epistasis^ library analyses was 300 cells, and for the Lenti-U6^BC^sgRNA^Saturation^/Cre library, it was 800 cells. Given that these were independent experiments, and the cut-off was applied uniformly to all the sgRNA and all the mice in those experiments, these differences in cut-offs do not impact overall data interpretation. Furthermore, we confirmed that our results were robust to using different cell number cutoffs.

#### C. Summary statistics for overall growth rate

To quantify the impact of each sgRNA on tumor growth, we normalized tumor statistics for a given sgX tumors against tumors initiated by control sgRNAs, including safe targeting sgRNAs and non-targeting sgRNAs (together referred to as *sgInerts*). We used two key measures: the size of tumors at defined percentiles of the distribution and the log-normal mean (LN mean) size. Percentile sizes provided a nonparametric summary, focusing on the larger tumors to avoid confounding factors from guide efficiency variations. The LN mean represented the average tumor size, assuming a log-normal distribution. The choice of log-normal distribution was described previously (Rogers et al. 2017). By normalizing these measures to the corresponding sg*Inert* statistics, we derived ratios that indicated the growth advantage or disadvantage of each genetic perturbation relative to *sgInert* tumors.

For example, the relative i^th^ percentile size for tumors of genotype X was calculated as:

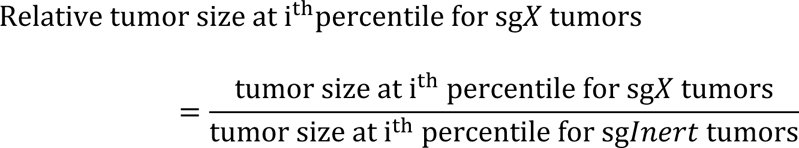

Likewise, the relative LN mean size for tumors of genotype X was calculated as:

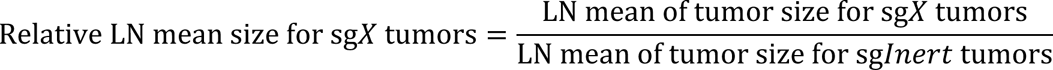

#### D. Summary statistics for relative tumor number and relative tumor burden

We analyzed the effects of gene inactivation on tumorigenesis by computing the number of tumors (“tumor number”) and total number of neoplastic cells (“tumor burden”) associated with each sgRNA. Unlike tumor size metrics, tumor number and burden were linearly influenced by lentiviral titer and affected by differences in the percent representation of each sgRNA in the viral pool. To account for these variables, each Tuba-seq^Ultra^ experiment included a cohort of *KT* control mice. *KT* mice lack expression of Cas9, render all sgRNAs functionally equivalent to inert sgRNAs. Consequently, the observed tumor number and burden associated with each sgRNA in these mice reflected their respective representation within the viral pool. Using KT mice as controls, we were able to accurately assess the effects of gene inactivation on tumor initiation and/or early clonal growth, while accounting for variabilities introduced by the viral vector composition.

To evaluate the impact of a given gene (*X)* on tumor number, we first normalized the number of sg*X* tumors in *KT;H11^LSL-Cas9^* mice (also *KT; p53^fl/fl^;H11^LSL-Cas9^* and *Braf^LSL-V600E/+^T; H11^LSL-Cas9^* mice) by the number of sg*X* tumors in the *KT* mice:

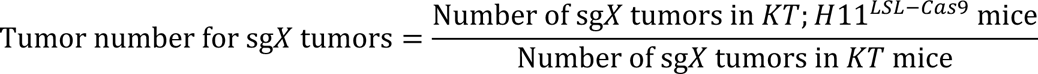

We then calculated a relative tumor number by normalizing this statistic to the corresponding statistic calculated using *sgInert* tumors:

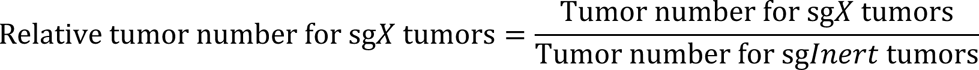

Prior work suggest that genes affect tumor initiation and/or the early clonal growth are imperfectly correlated with tumor growth at later stages (Cai et al. 2021). Relative tumor number thus captures an additional and potentially important aspect of gene function.

Analogous to the calculation of relative tumor number, we calculated the effect of each sgRNA on tumor burden by first normalizing the sgX tumor burden in Cas9-expressing mice to the burden in *KT* mice:

Tumor burden for sg*X* tumors

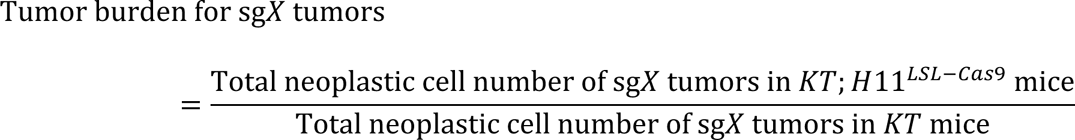

We then calculated a relative tumor burden by normalizing this number to the corresponding statistic calculated using *sgInert* tumors:

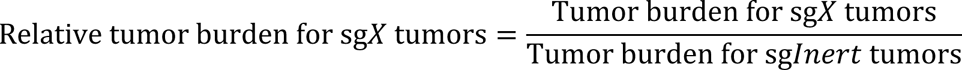

Tumor burden combines size and number effects, providing a measure of the total neoplastic load per mouse. Overall tumor burden tends to have higher variability due to the lack of clonal data and is heavily influenced by the size of the largest tumors present.

#### E. Calculation of confidence intervals and p-values for tumor growth and number metrics

Confidence intervals and p-values were calculated using bootstrap resampling for each growth metric. To account for variability across mice and within-mouse, we used a two-step, nested bootstrap approach: first resampling mice, then resampling tumors within each mouse. This process was repeated 10,000 times to generate 10,000 values for each growth metric. The 95% confidence intervals were determined using the 2.5^th^ and 97.5^th^ percentiles of the bootstrapped statistics. Since we normalize tumor growth metrics to the same metrics in sg*Inert* tumors, the test statistic under the null hypothesis (no genotype effect on tumor growth) is equal to 1. Two-sided *P*-values were thus calculated as followed:

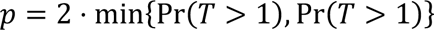

Where T is the test statistic and Pr(*T*>1) and Pr(*T*<1) were calculated empirically as the proportion of bootstrapped statistics that were more extreme than the baseline of 1. To account for multiple hypothesis testing, *P*-values were FDR-adjusted using the Benjamini-Hochberg procedure (Benjamini and Hochberg 1995). Summarized statistics of all experiments in this study can be found in **Supplementary Tables 3, 8 and 27**.

#### F. Computing gene level metrics

The gene-level metrics (M_gene_) are estimated using a weighted average of sgRNA-level metrics (M_sgRNA_):

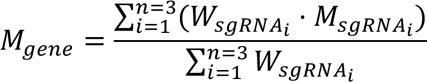

where *W_sgRNAi_* is the relative tumor number of sgRNA_i_ for the focal gene.

### Analyses of dual-sgRNA tumor number

Raw reads processing and tumor number were described previously (Hebert et al. 2024a).

Relative tumor numbers (TN) for each dual-sgRNA are calculated by:

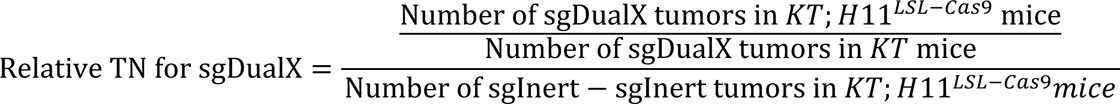

### Epistasis Analyses

Genes were categorized as negatively epistatic with *Kat7* (i.e., reduced growth effects) if their log-normal mean tumor size in *KT;H^11LSL-Cas9^* mice was greater than the upper bound of the 95% confidence interval of the log-normal mean tumor size in *K;H^11LSL-Cas9^;Kat7^fl/fl^*mice. Conversely, genes were considered positively epistatic with *Kat7* (i.e., increased growth effects) if their log-normal mean tumor size in *KT;H^11LSL-Cas9^* mice was less than the lower bound of the 95% confidence interval of the log-normal mean tumor size in *K;H^11LSL-Cas9^;Kat7^fl/fl^* mice. Genes that did not meet either of these criteria were classified as non-epistatic with *Kat7*.

### Plasmid library quality control

To ensure the quality of plasmid library, the plasmid library was deep sequenced (**see Tuba-seq^Ultra^ library preparation**). The raw reads were processed as described above (**see Raw reads parsing and cleaning**). We then examined the fraction of vector containing the expected sgRNAs and clonal barcode patterns. More than 98% of the vectors matched the expected sgRNA and barcode patterns across all plasmid libraries, indicating that a large usable fraction of the virus would be transduced into the mouse lung.

We further ensured that (1) each designed sgRNA was present in the library, (2) the number of clonal barcodes associated with each sgRNA was sufficiently large, and (3) the distribution different sgRNAs was relatively even in the library. In the Lenti-U6^BC^sgRNA^Epigenetics^/Cre library, one of the sgRNA targeting *Setd2* was absent because its sequence contained an Esp3I digestion site. Finally, we confirmed that the total number of sgRNA and clonal barcode combinations was sufficiently large (>10^6^) to greatly surpass the expected tumor initiation in each mouse lung (<50,000 tumors).

### Tuba-seq^Ultra^ cost and labor estimation

Cost analysis comparing Tuba-seq and Tuba-seq^Ultra^ assumes the experiments were being performed by an experienced personnel familiar with the protocols working. The time required for each experiment was based on the Winslow lab protocols. The costs were estimated from prices in the year 2020, with each viral prep costing approximately $250 USD per day in labor and $30 USD in lab consumables. For Tuba-seq, where cloning and viral packaging must be done individually for each sgRNA, we assumed that an experienced person can at most produce 15-18 Lenti-sgRNA/Cre viral preps at a time. Therefore, the labor cost would scale linearly when the library size increases by 15-18 sgRNAs. Other variables that may influence the cost of labor or consumables such as technical variability, logistical delays, or market fluctuations were not included.

### Kaplan-Meier survival analysis

Survival analyses for mice transduced with Lenti-sgRNA/Cre vector were conducted in R version 4.4.1 using packages “survival” (v3.7-0) and “survminer” (v0.4.9). Days to death were determined from the day of transduction to the day the animal was harvested. Separate cohorts of mice initiated with the same sgRNA were combined. The comparison groups and the number of animals for each group are indicated in the figures. For the comparison involving Lenti-sgPsip1/Cre, the Lenti-sgSafe/Cre used combined data from mice transduced with *sgSafe14* (n=7) or *sgSafe23* (n=7) vectors. For experiments involving Lenti-sgKat7/Cre, the survival data for the cohort of *KT;H11^LSL-Cas9^* (n = 3) and *KT* (n = 3) mice that received 120,000 ifu per mouse were combined with the cohort of *KT;H11^LSL-Cas9^* (n = 10) and *KT* (n = 4) mice that received 200,000 ifu per mouse for survival analysis. Statistical significance was determined by log-rank test using the survfit function in the “survival” package.

### Western blot quantification

Western blots were performed on FACS-sorted Lin^−^TdTomato^+^ neoplastic cells from 3 independent mice for each target gene. Quantification was performed using ImageJ (FIJI, v2.14) by selecting each lane and quantifying with the Analyze → Plot lanes function. The data for the target band were normalized to β-ACTIN from the same lane. We reasoned that inactivation of the HBO1 complex, a histone writer complex, would not increase the levels of target histone marks. Therefore, statistical significance was determined by a one-sided Student’s t-test in R v4.4.1.

### DepMap data analysis

Correlation data for each gene was accessed from the DepMap Consortium portal website (https://depmap.org/portal/) (Tsherniak et al. 2017). The correlation coefficients were obtained from the gene’s top 100 co-dependencies from DepMap CRISPR screen data (DepMap 2023 Q4 public data release). For JADE1, JADE3, ING4, BRPF1, and BRPF2, where their top 100 co-dependencies did not include genes from the HBO1 or MLL1 complexes, the correlation coefficients were computed in R (version 4.4.1) using Spearman’s correlation.

### Human mutation data

Mutation frequency and mutation types in human lung adenocarcinoma were accessed from cBioportal (https://www.cbioportal.org/) (Cerami et al. 2012; Gao et al. 2013; de Bruijn et al. 2023). Our analyses can be retrieved from the following URLs: https://bit.ly/3Rgt5YG (HBO-JADE2 genes), https://bit.ly/3A2SQ8I (MLL1 tumor suppressive genes) and https://bit.ly/3VHViKB (combined genes). Because this study focuses on gene loss-of-function, samples with copy number amplifications were not included in **Figure 4H** and **Supplemental Figure 8A**.

### Histopathology evaluation

Defining the stain intensity of the human lung adenocarcinoma tissue microarray samples as absent, low, medium, or high was established by examining the normal lung tissues as the reference. All histone markers including H3K14ac, H4K12ac, H4K5ac and H3K36me3 showed very dark and uniform stains in the nuclei of normal lung tissues, which was defined as “high” intensity. The other intensity levels were determined in reference to the normal lung tissues and to the normal cells in the tumor tissue. Cancer and stromal cells were identified based on cell and nuclear morphology. “Absent” was defined as samples with high-level staining in non-cancerous stromal cells but no stain in the cancer cells. Low and medium staining levels were defined similarly, where the staining intensity was less relative to normal cells in the same sample and to normal lung samples. To ensure reproducibility in staining intensity assessment, randomly selected samples from the tissue microarray slide stained with H3K14ac were independently assessed by P.R and M.V.R with high agreement with Y.J.T.

Statistical significance was calculated with the FSA package in R (Ogle DH, Doll JC, Wheeler AP, Dinno A (2023). *FSA: Simple Fisheries Stock Assessment Methods*. R package version 0.9.5, https://fishr-core-team.github.io/FSA/.) using the Kruskal-Wallis test followed by Dunn’s test to correct for multiple comparisons. Correlation between histone markers was calculated by Spearman’s correlation in R version 4.4.1.

Tumor grade assessment was performed on H&E slides of tumor-bearing lung lobes harvested from *KT* and *KT;H11^LSL-Cas9^;Kat7^fl/fl^*mice transduced with the Lenti-sgSafe23/Cre vector. The assessment was performed by a U.S. board-certified clinical pathologist (M.V.R). To ensure reproducibility, the slides were assessed in a blinded manner twice, approximately one month apart. Statistical significance was determined by Chi-squared test in R version 4.4.1.

### Gene Ontology analysis

To identify molecular processes that were enriched by functionally significant epigenetic regulatory genes, we defined genes as tumor suppressive if relative mean tumor size ≥ 1.1 and p.adj. ≤ 0.05, or relative tumor number ≥ 1.1 and p.adj. ≤ 0.05. We defined genes as tumor dependent if relative mean tumor size ≤ 0.9 and p.adj. ≤ 0.05, or relative mean tumor number ≤ 0.9 and p.adj. ≤ 0.05. We applied the ± 10% criteria to focus on biologically and statistically significant epigenetic regulatory genes in lung tumorigenesis. We further filtered for genes that have at least 2 sgRNAs that are statistically significant. Genes *Vsp72*, *Suz12*, and *Chd8* were filtered from the tumor suppressive gene list because one of the 3 sgRNAs had a negative size effect. Genes that have opposing growth effects such as increasing tumor size and decreasing tumor number were also filtered to avoid ambiguity in gene ontology enrichment. The gene lists were analyzed using https://geneontology.org/ set for human Molecular Functions.

### Bulk-ATAC Analysis

For each ATAC-seq experiment, computational processing was carried out as previously described (Marinov and Shipony 2021; Marinov et al. 2023a). Demultiplexed FASTQ files were mapped to the GRCm38/mm10 assembly of the *Mus musculus* genome as 2×36mers using Bowtie (Langmead et al. 2009) with the following settings: -v 2 -k 2 -m 1 --best --strata. Mitochondrial reads were filtered, and duplicate reads were removed using picard-tools (version 1.99). Browser tracks generation, fragment length estimation, TSS enrichment calculations, and other analyses were carried out using custom-written Python scripts https://github.com/georgimarinov/GeorgiScripts. The refSeq annotation was used for evaluation of enrichment around TSSs. Peak calling was carried out using MACS2 (Feng et al. 2012), with the following settings: -g mm -f BAMPE --to-large --keep-dup all --nomodel. Peaks were compared against the ENCODE set of “blacklisted” regions to filter out likely artifacts.

The processed peaks were merged via iterative summits analysis to identify high-confidence peaks (https://github.com/corceslab/ATAC_IterativeOverlapPeakMerging, R version 4.2.2) (Corces et al. 2018; Granja et al. 2021). The GRCm38/mm10 ENCODE blacklist regions were obtained from (https://github.com/Boyle-Lab/Blacklist/blob/master/lists/mm10-blacklist.v2.bed.gz). The iterative summits were converted to .saf files using bedops (v2.4.41), and peak counts were generated by the featureCounts function in subread (v2.0.3).

Differential peak analysis was performed separately using DESeq2 (v1.4.4) (Love et al. 2014) for each ATAC-seq experiment. Relative chromatin accessibility of each genotype was determined by using *sgSafe* samples as the reference within each ATAC-seq experiment. Peak annotation was carried out using annotatepeaks.pl in HOMER (v4.1.1) (Heinz et al. 2010) and ChIPSeeker (v1.4.0) (Yu et al. 2015). Since we have two sets of ATAC-seq experiments, to identify overlapping genomic regions between the *sgKat7* cells and the other genotypes, another set of peak counts was generated for the *sgKat7* cells using genomic coordinates of the iterative summits from the other ATAC-seq experiment involving *sgKmt2a*, *sgPsip1,* and *sgMeaf6* with subread (v2.0.3). The statistically significant peaks (p.adj ≤ 0.05) were used to generate the heatmaps. Motif enrichment was performed using findmotifsgenome.pl from HOMER (v4.1.1).

### Single-cell RNA-seq data analysis

The FASTQ files were preprocessed with the data analysis pipeline from Parse Biosciences (split-pipe v1.0.3). Default settings were used for aligning to GRCm38/mm10 mouse reference genome. Each sub-library was processed individually using the command split-pipe-mode all, and the outputs were combined with the command split-pipe-mode combine. The metrics generated by the split-pipeline were used for **Supplemental Table 20**.

Gene expression analysis was performed using Scanpy (v1.9.3) (Wolf et al. 2018). Each genotype was analyzed separately with the *sgSafe* cells from the same experiment as the reference. Cells meeting the following criteria: 1) the number of genes expressed was less than 6000; 2) total read count was less than 40,000; and 3) the percent of mitochondria genes was less than 10% were included in the analysis. Scrublet was used to assign doublet scores with an expected doublet rate of 7.5% (Wolock et al. 2019). Doublets detected by Scrublet were removed.

Read count normalization was performed with scanpy.pp.normalize_total using default parameters and log-transformed with scanpy.pp.log1p. Feature selection was performed with scanpy.pp.highly_variable_genes on the normalized and scaled data to identify highly variable genes with the parameters: min_mean=0.0125, max_mean=3, min_disp=0.5. The data was further filtered with scanpy.pp.regress_out by regressing out total read count and percent mitochondrial genes. Then each gene was scaled to unit variance and values exceeding 10 standard deviations were clipped. UMAP embedding with Leiden clustering at resolutions between 0.5-1.5 was used to visualize the data.

Non-neoplastic cells were identified based on clustering patterns and expression of commonly accepted marker genes: Vim*/Pdgfra/S100a4/Acta2* (mesenchymal cells), *Ptprc/Ccr5* (lymphocytes), *Mcam/Pcam1/Vcam1* (perivascular cells), and *Cd34/Tek* (endothelial cells). These cells were filtered from downstream analysis.

Assignment of genotype specific clusters on UMAP was guided by three general principles after visual inspection and calculation of the percent contribution of each sgRNA to a cell cluster. First, CRISPR-Cas9 mediated gene editing is not 100% efficient, thus some cells from mice transduced with gene-targeting sgRNA will be present in clusters of mice transduced with *sgSafe* vectors. Second, scRNA-seq captures a gradient of cell states including transitional states, and selecting cell clusters that are 100% of a genotype is overtly exclusionary. Third, different gene inactivation may have different effects on cell states and the transcriptome, therefore may require different cut-offs for cell cluster assignment. For *sgKat7*, cell clusters with more than 90% of cells from Lenti-sgKat7/Cre transduced mice were considered *sgKat7* clusters, and the rest were *sgSafe* clusters. For *sgMeaf6*, cell clusters with more than 85% of cells from Lenti-sgMeaf6/Cre transduced mice were considered *sgMeaf6* clusters, and the rest were *sgSafe* clusters. For *sgKmt2a*, cell clusters with more than 65% of cells from Lenti-sgKmt2a/Cre transduced mice were considered *sgKmt2a* clusters, and the rest were *sgSafe* clusters. For *sgPsip1*, the same cut-off as *sgKmt2a* was used.

Differential gene expression was conducted using default parameters from scanpy.tl.rank_gene_groups comparing genotype specific cell clusters with *sgSafe* cell clusters set as the reference. Genes with FDR corrected p-value ≤ 0.05 were considered statistically significant.

### Gene set over-representation and cell type signature analysis for scRNA-seq

To perform enrichment analysis of molecular processes, we separated the statistically significant genes into up-regulated genes (log2 fold-change > 0 and FDR corrected p-value ≤ 0.05) or down-regulated genes (log2 fold-change < 0 and FDR corrected p-value ≤ 0.05). The up-regulated and down-regulated gene lists were ranked based on their absolute log2 fold-change and the top 3,000 genes were selected for inputs into Metascape (Zhou et al. 2019) due to its input limit ((metascape.org). The GO_AllList output from each genotype was used to identify commonly enriched molecular processes. We filtered for Gene Ontology processes to reduce redundancy in curated databases.

Cell type signatures were imported from the GO_Cell_Type_Signatures output for each genotype from Metascape. Cell type signatures with FDR adjusted p ≤ 0.05 were considered. Since the cell type signatures from Metascape are from human data (https://www.gsea-msigdb.org/gsea/msigdb/human/genesets.jsp?collection=C8), we used mouse marker gene signatures from PanglaoDB (Franzen et al. 2019) for the same cell types as the Metascape human data to generate the gene signature scores in Scanpy with the function scanpy.tl.score_genes. The gene signature scores were visualized on UMAPs. The mouse genes for each gene signature are included in **Supplemental Table 24**. Statistical significance was determined with Mann-Whitney U test comparing the mean gene signature scores between cells in gene inactivated clusters and cells in the *sgSafe* clusters. Marker genes used in dot plots were also cross-checked on Mouse Cell Atlas (Fei et al. 2022) to ensure that they represent the top genes expressed in the specified cell type.

### Data availability

All Tuba-seq^Ultra^ datasets will be made publicly available through the NCBI’s Sequence Read Archive Database and accession numbers will be provided prior to publication of the manuscript. All ATAC-seq and scRNA-seq data will be made publicly available through the Gene Expression Omnibus and accession numbers provided prior to publication of the manuscript.

### Code availability

The code used for data analysis in this study will be made available on GitHub prior to publication of the manuscript.

## ACKNOWLEDGEMENTS

We thank the Stanford Shared FACS Facility for FACS related services, the Stanford Veterinary Service Center staff for animal care, and Stanford Protein and Nucleic Acidy Facility. We thank Sushama Varma from Stanford Pathology Department for preparing the TMA slides. We thank members of the Winslow, Petrov laboratories and Belal Yasin for experimental support. Y.J.T was supported by a Canadian Institute of Health Research (CIHR) postdoctoral fellowship (MFE-176568). H.X is supported by a Tobacco-Related Disease Research Program (TRDRP) fellowship (T34FT8013). J.D.H is supported by an American Cancer Society postdoctoral fellowship (PF-21-112-01-MM) and a TRDRP fellowship (T31FT1619). S.K is supported by National Cancer Institute’s F99/K00 CA234962. E.G.S is supported by a TRDRP predoctoral fellowship (T33DT6556). D.N.D is supported by the National Cancer Institute Predoctoral to Postdoctoral Fellow Transition Award (K00-CA245784). E.L.A. was supported by the HHMI Gilliam Fellowship for Advanced Study (GT14928) and a National Cancer Institute Predoctoral to Postdoctoral Fellow Transition Award (F99CA284289). G.B is supported by the National Research, Development and Innovation Office (RRF-2.3.1-21-2022-00006) and by the HUN-REN Welcome Home and Foreign Researcher Recruitment Program. M.V.R is supported by the D.K. Ludwig Fund for Cancer Research. This work is supported by R01-CA234349 (to D.A.P and M.M.W), R01-CA230025 (to M.M.W), U01-1146AG077922 (to M.M.W), and in part by the Stanford Cancer institute support grant (NIH P30-1147CA124435).

## CONTRIBUTIONS

Y.J.T and M.M.W conceptualized and designed the study. Y.J.T and M.M.W designed the Tuba-seq^Ultra^ methodology. YJ.T led experimental data collection with contributions from S.L, S.K, J.D.H, L.A, P.R, P.C, C.D, D.N.D, G.B, and E.L.A. L.A. and Y.J.T prepared Tuba-seq^Ultra^ libraries. H.X and Y.J.T designed the computational pipelines for Tuba-seq^Ultra^ analysis with input from N.W.H and E.G.S. H.X performed the Tuba-seq^Ultra^ analysis with help from Y.J.T and N.W.H. G.B. performed dual-sgRNA screen analysis. Y.J.T. and S.H.K performed the ATAC-seq experiments and analyzed the data. S.P. contributed intellectually to the study. A.K.V and T.T contributed intellectually and provided the *Kat7^fl/fl^* mice. Y.J.T and M.M.W wrote the manuscript with comments and contributions from all authors. M.V.R provided TMAs and assessed tumor grades and IHC scores. M.M.W, D.A.P and W.J.G provided resources. M.M.W supervised the study.

## Supplemental Figure Legends

**Supplemental Figure 1.**
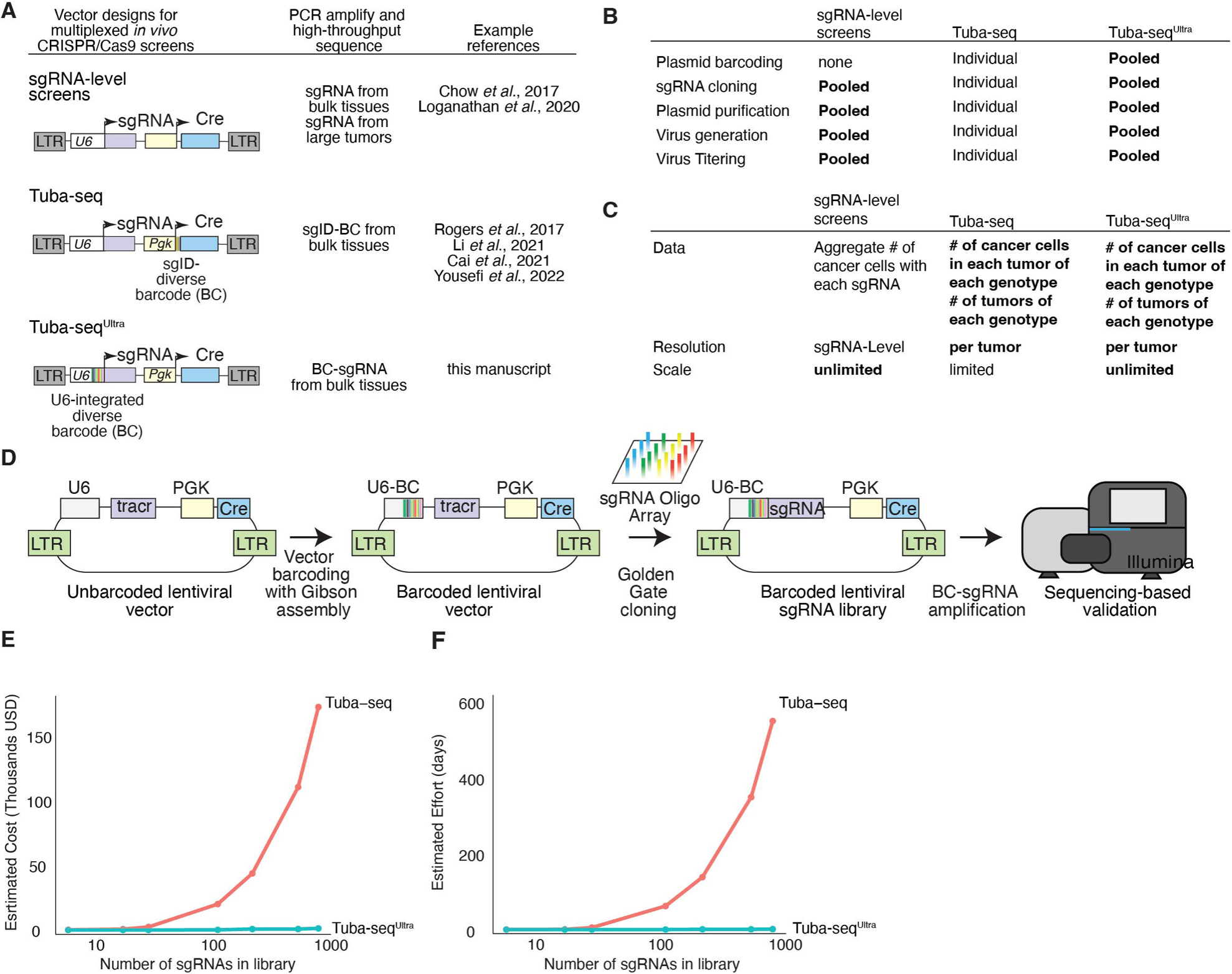
Tuba-seq^Ultra^ enables scalable and cost-effective functional genomics *in vivo* with per-tumor resolution in autochthonous cancer models. **A.** Vectors for somatic cell gene editing and *in vivo* CRISPR/Cas9-mediated screening in cancer models. sgRNA-level screens rely on amplification and sequencing of the sgRNA or targeted indels and generate only aggregate data for each sgRNA, without any ability to identify the impact on tumor number and tumor size. Our original Tuba-seq approach has an sgRNA identifier and random barcode (sgID-BC) distal to the sgRNA, while Tuba-seq^Ultra^ places the barcode immediately adjacent to the sgRNA. **B.** Features of sgRNA-level, Tuba-seq, and Tuba-seq^Ultra^ vector pools have important implications for cloning and virus generation. While the sgRNA-level screen lacks clonal information, Tuba-seq screens require the sgRNA and sgID-BC region to be cloned individually into each vector. Furthermore, for Tuba-seq, each lentiviral preparation needs to be generated individually to eliminate lentiviral template switching, which would uncouple the sgID-BC from the sgRNA. The Tuba-seq^Ultra^ design allows all steps to be performed in a pooled manner. **C.** Scale and resolution of sgRNA-level, Tuba-seq, and Tuba-seq^Ultra^ screens. **D.** Cloning of Tuba-seq^Ultra^ libraries follows a similar protocol to the generation of most sgRNA libraries, except that the sgRNA pool is cloned into a vector backbone with a barcoded U6 promoter. **E-F.** Given the features described in **Supplementary Figure 1B**, the estimated cost **(E)** or required effort **(F)** of Tuba-seq^Ultra^ libraries stay nearly unchanged when increasing the number of sgRNAs in the library, while Tuba-seq library generation becomes cost- and effort-prohibitive as library size increases.

**Supplemental Fig 2.**
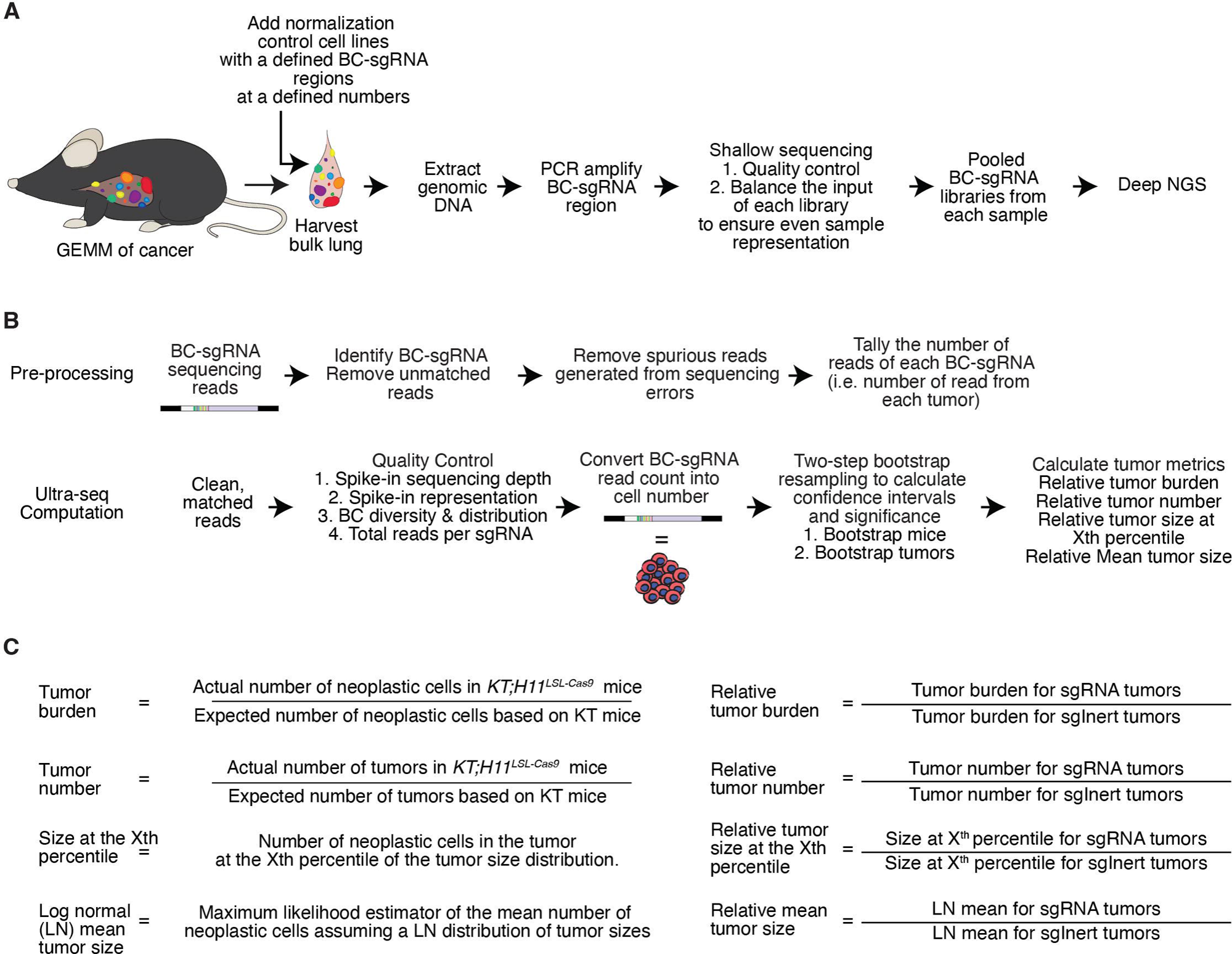
Outline of Tuba-seq^Ultra^ sample processing, analysis of high-throughput barcode-sgRNA sequencing data, and calculation of tumor metrics. **A.** Outline of sample processing for sequencing-based quantification of the number of tumors generated by each sgRNA and the number of neoplastic cells (tumor size) of each clonal tumor for every sgRNA from bulk tumor-bearing lungs. **B.** Processing of barcode-sgRNA reads to accurately quantify tumor growth metrics and calculate the statistical significance of effects. **C.** Calculation of tumor metrics. “Tumor burden” and “Tumor number” are relative to the data from *KT* mice (as they need to be normalized to the precise representation of each vector in the library). In contrast, “Size at the Xth percentile” and “Log-normal (LN) mean tumor size” are calculated for each sgRNA relative to all *sgInert* tumors for within each *KT;H11^LSL-Cas9^* cohort. Inall cases, the “Relative” metrics represent the metric for the sgRNA of interest normalized to that metric for all *sgInert* tumors.

**Supplemental Figure 3.**
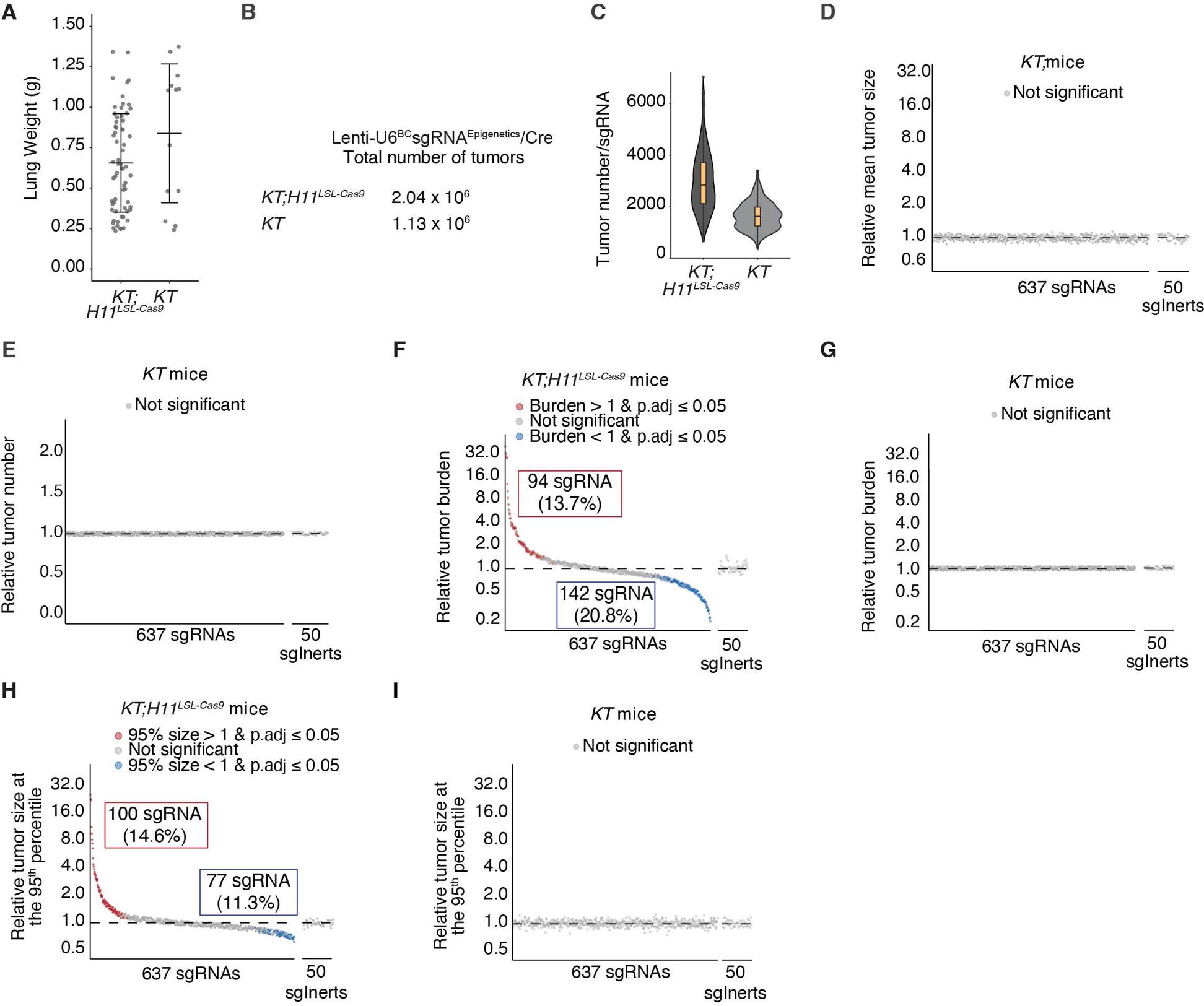
Tuba-seq^Ultra^ enables the analysis of multiple metrics of tumorigenesis from millions of clonal autochthonous lung tumors. **A.** Weight of tumor-bearing lungs from the indicated mouse genotypes. Each dot represents a mouse. Error bars are mean +/− standard deviation. **B.** Total number of tumors analyzed in the indicated mouse genotypes. **C.** The number of tumors analyzed for each sgRNA for the indicated mouse genotypes. Boxes are the first to third quartile, and the median is indicated by the horizontal line. The whiskers are Interquartile Range (IQR) multiplied by 1.5 in each direction. **D-E.** Impact of each sgRNA on tumor growth (Relative mean tumor size) (**D**) and tumor initiation/early growth (Relative tumor number) (**E**) in Cas9-negative *KT* mice. Each dot is a sgRNA. No sgRNAs have significant effects in the absence of Cas9. **F-G.** Impact of each sgRNA on tumor burden (Relative tumor burden) in *KT;H11^LSL-Cas9^*(**F**) and Cas9-negative *KT* mice (**G**). Each dot is a sgRNA. Significant sgRNAs are colored as indicated. Note that tumor burden does not incorporate barcode information, and hence, identified many fewer sgRNAs as having significant effects. **H-I.** Impact of each sgRNA on tumor sizes at the 95th percentile of the size distribution (Relative tumor size at the 95th percentile) in *KT;H11^LSL-Cas9^* **(H)** and Cas9-negative *KT* mice **(I)**. Each dot is a sgRNA. Significant sgRNAs are colored as indicated.

**Supplemental Figure 4.**
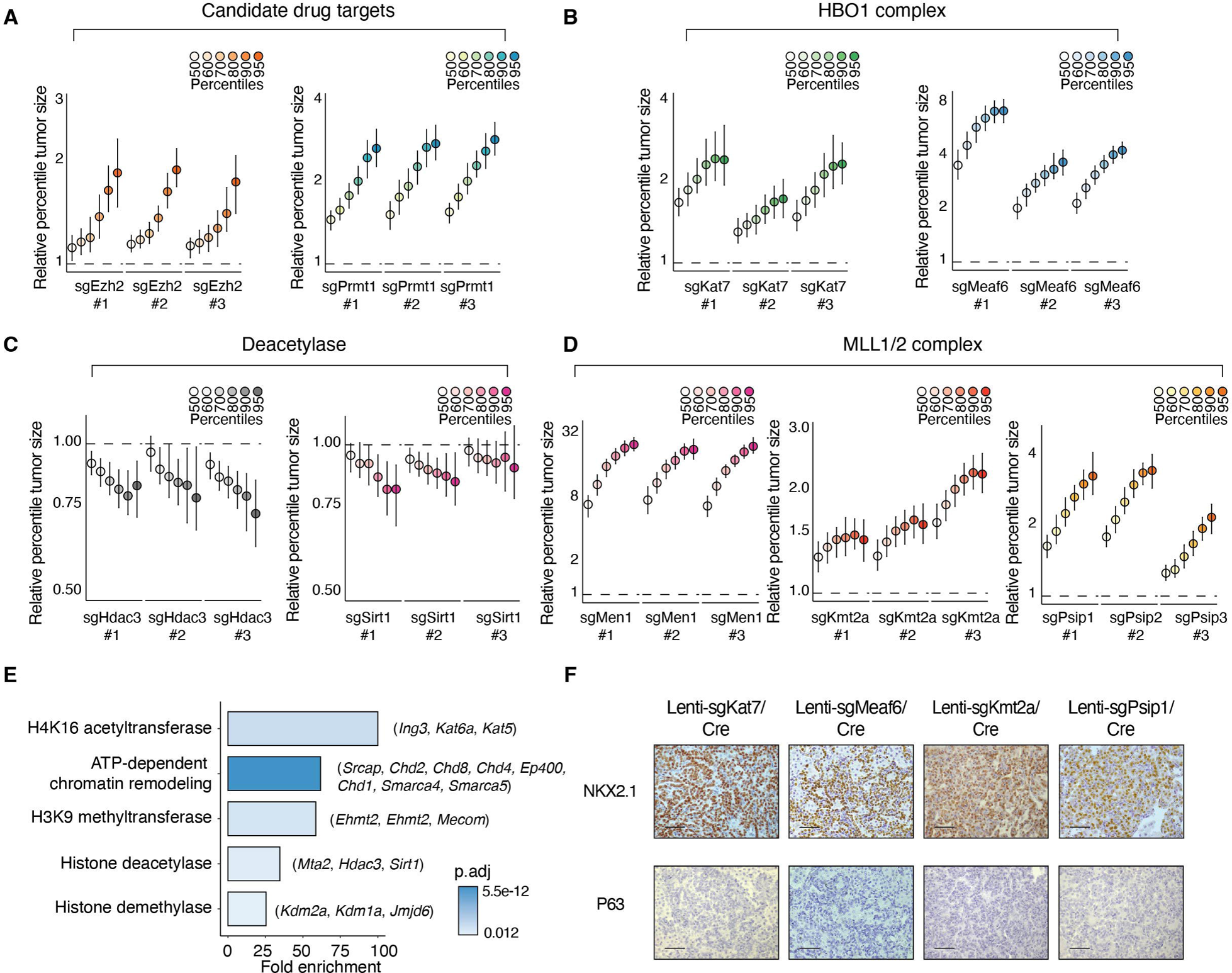
Tumor-suppressive and tumor-dependency epigenetic regulators in lung tumorigenesis. **A-D.** Tumor sizes at the indicated percentiles of sgRNAs targeting *Ezh2* and *Prmt1* **(A)**; *Kat7* and *Meaf6* **(B)**; *Hdac3* and *Sirt1* **(C)**; and *Men1*, *Kmt2a* and *Psip1* **(D)** in *KT;H11^LSL-Cas9^* mice. Each column indicates a different sgRNA. Error bars are 95% confidence intervals. The dotted lines indicate no effect. **E.** Functional annotation from Gene Ontology of tumor dependent epigenetic regulators. Gene hits for each functional category are shown in brackets. Statistical significance is determined by Fisher’s exact test with Benjamini-Hochberg correction. **F.** Immunohistochemistry images of NKX2.1 and p63 on lung tumors from the indicated genotypes (scale bar, 50 μm).

**Supplemental Figure 5.**
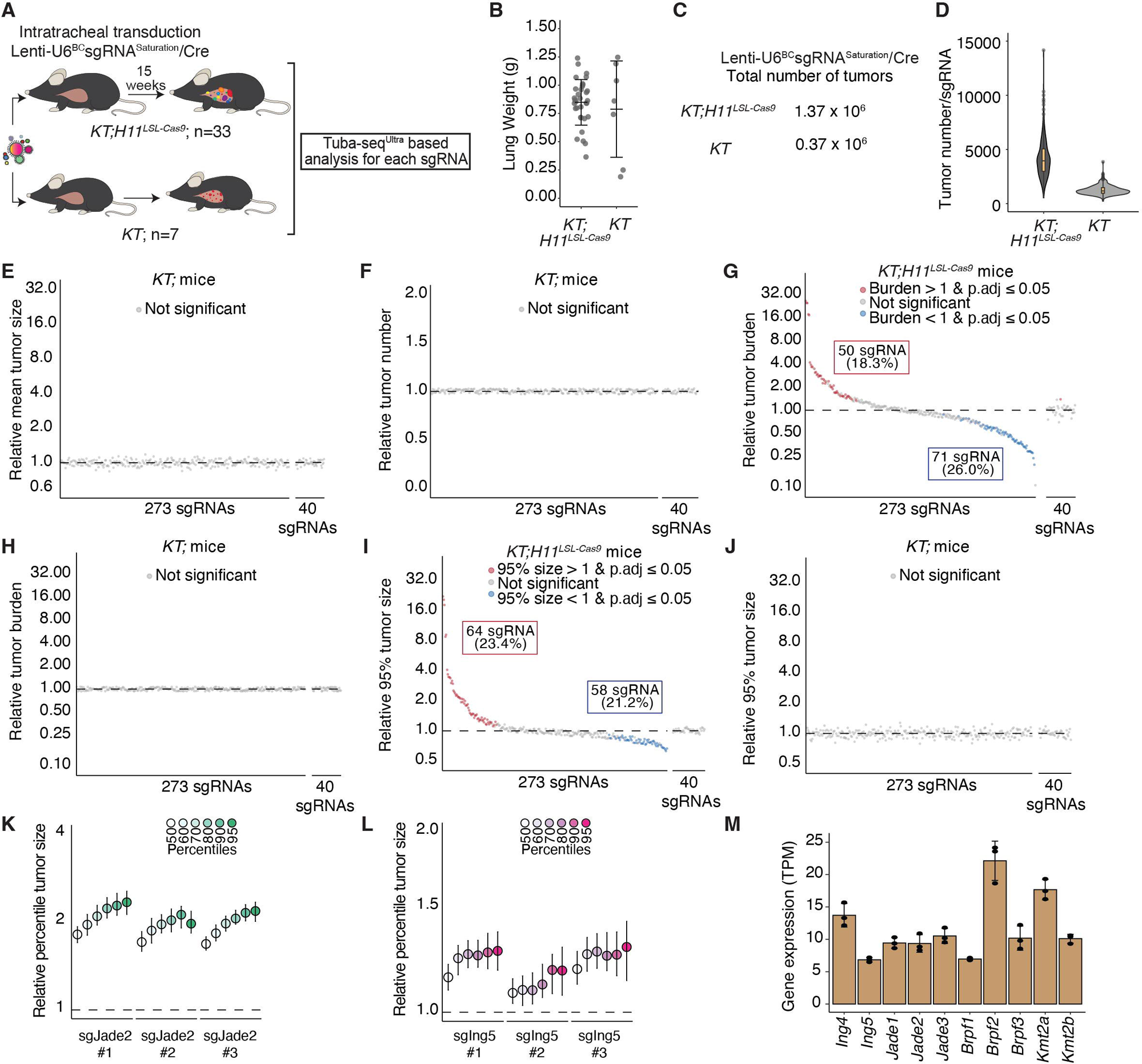
High-throughput MYST and COMPASS saturation analyses identify additional epigenetic regulators driving lung tumorigenesis. **A.** Initiating tumors with Lenti-U6^BC^sgRNA^Saturation^/Cre in the indicated numbers of *KT;H11^LSL-Cas9^* and *KT* mice. Tumors developed for 15 weeks before Tuba-seq^Ultra^ analysis. **B.** Weight of tumor-bearing lungs from the indicated genotypes of mice. Each dot represents a mouse. Error bars are mean +/− standard deviation. **C.** Total number of tumors analyzed in the indicated mouse genotypes. **D.** The number of tumors analyzed for each sgRNA in the indicated mouse genotypes. Boxes are the first to third quartile, and the median is indicated by the horizontal line. The whiskers are Interquartile Range (IQR) multiplied by 1.5 in each direction. **E-F.** Impact of each sgRNA on tumor size (**E**) and tumor number (**F**) in *KT* mice. Each dot is a sgRNA. **G-H.** Impact of each sgRNA on tumor burden in *KT;H11^LSL-Cas9^* (**G**) and *KT* mice (**H**). Each dot is a sgRNA. Significant sgRNAs are colored as indicated. **I-J.** Impact of each sgRNA on tumor sizes at the 95th percentile of the size distribution in *KT;H11^LSL-Cas9^* (**I**) and *KT* mice (**J**). Each dot is a sgRNA. Significant sgRNAs are colored as indicated. **K-L.** Tumor sizes at the indicated percentiles for sgRNAs targeting *Jade2* **(K)** and *Ing5* **(L)**. Error bars are 95% confidence intervals. The dotted line indicates no effect. **M.** Expression in transcripts per million (TPM) of gene paralogs in the HBO1 and MLL1/2 complexes in *KT* lung tumors (Chuang et al., 2017).

**Supplemental Figure 6.**
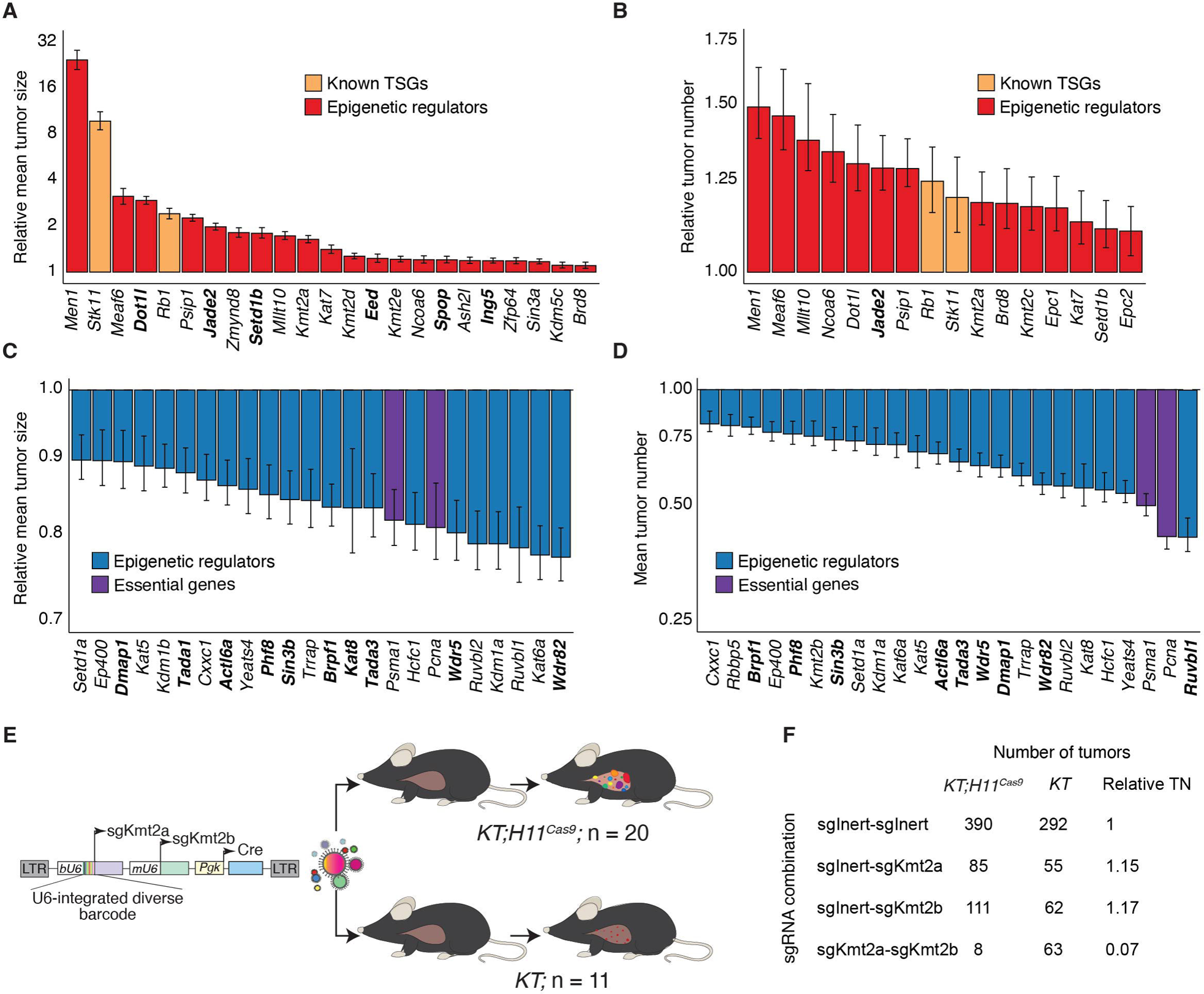
Tumor-suppressive and tumor-dependency genes in the Lenti-U6^BC^sgRNA^Saturation^/Cre library. **A-B.** Tumor-suppressive genes from the Lenti-U6^BC^sgRNA^Saturation^/Cre library that impact tumor size (**A**) and tumor number (**B**). Each bar represents the log-normal mean size of aggregated data from all sgRNAs for that gene. The error bars are 95% confidence intervals. **C-D.** Tumor-dependent genes from the Lenti-U6^BC^sgRNA^Saturation^/Cre library that impact tumor size (**C**) and tumor number (**D**). Each bar represents the log-normal mean size of aggregated data from all sgRNAs for that gene. The error bars are 95% confidence intervals. Newly included genes in the Lenti-U6^BC^sgRNA^Saturation^/Cre library are bolded. **E.** Initiating tumors with a barcoded dual-sgRNA vector targeting *Kmt2a* and *Kmt2b* in *KT;H11^LSL-Cas9^*and *KT* mice. N is the number of mice. **F.** The number of tumors for each sgRNA combination in *KT;H11^LSL-Cas9^* and *KT* mice. Relative TN (Tumor Number) is calculated as the number of tumors in *KT;H11^LSL-Cas9^* mice normalized to *KT* mice and then to *sgInert-sgInert*.

**Supplemental Figure 7.**
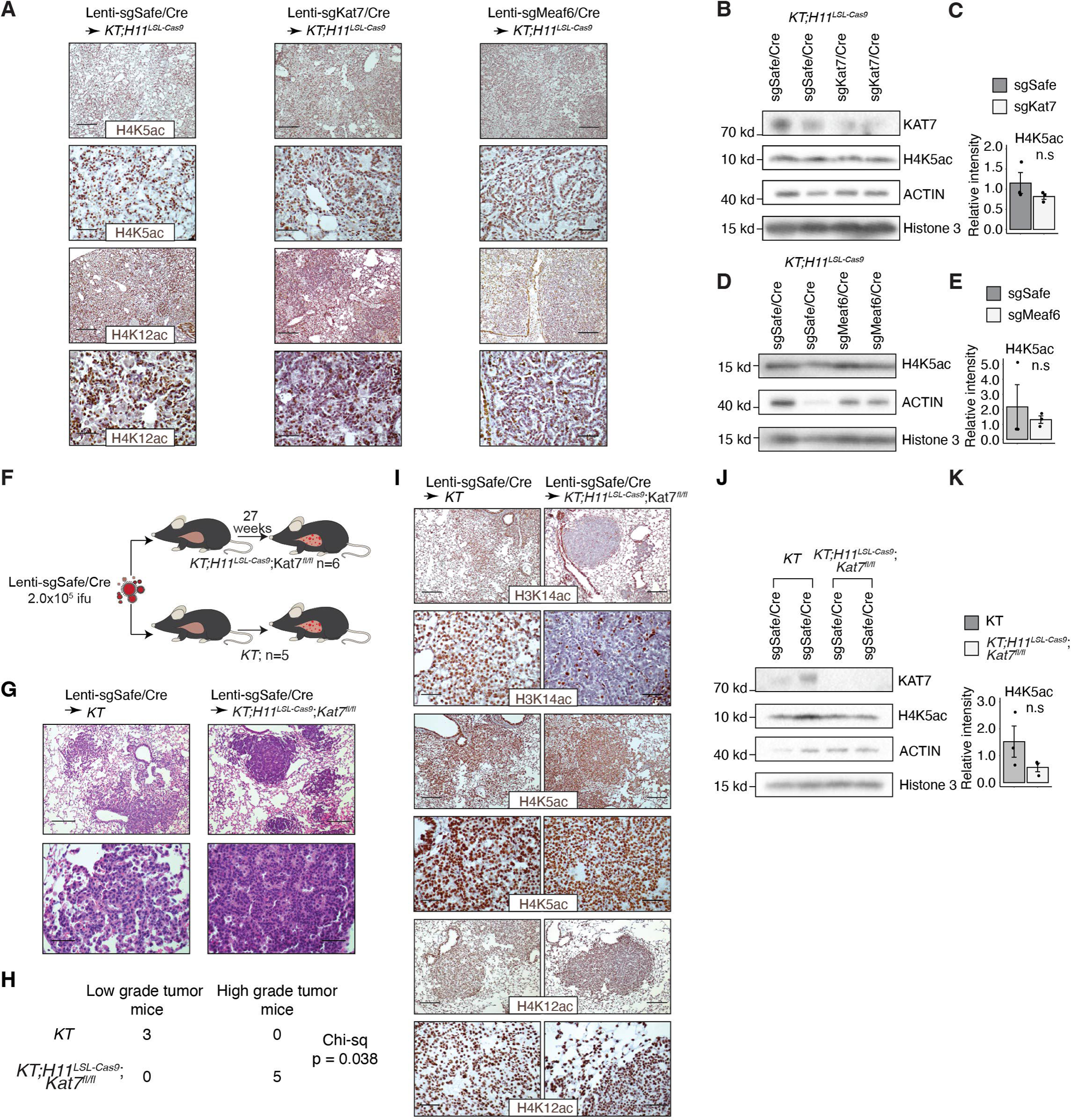
Inactivation of the HBO1 complex genes reduces H3K14ac and H4K12ac levels in lung tumors and worsens tumor grade. **A.** Representative immunohistochemistry images of H4K12ac and H4K5ac staining in mouse lung tumors initiated by different single sgRNAs in *KT;H11^LSL-Cas9^* mice. Bottom panels (scale bars, 50 μm) are higher magnification images of areas in the top panel (scale bars, 100 μm). **B.** Representative western blots of FACS-isolated lineage (CD45/CD31/Ter119/F4/80) negative, tdTomato positive (Lin^−^TdTom^+^) neoplastic cells from *KT;H11^LSL-Cas9^* mice transduced with Lenti-sgKat7/Cre or Lenti-sgSafe/Cre vector. Each lane on the western blot comes from a different mouse. **C.** Bar plots showing densitometry quantification of protein levels on western blots. Relative intensities are normalized to ACTIN. Error bars are +/− standard error of the mean, and each dot represents a different mouse. Statistical significance is determined by Student’s t-test. **D.** Representative western blots of FACS-isolated Lin^−^TdTom^+^ neoplastic cells from *KT;H11^LSL-Cas9^* mice transduced with Lenti-sgMeaf6/Cre or Lenti-sgSafe/Cre vector. Each lane on the western blot comes from a different mouse. **E.** Bar plots showing densitometry quantification of protein levels on western blots. Relative intensities are normalized to ACTIN. Error bars are +/− standard error of the mean, and each dot represents a different mouse. Statistical significance is determined by Student’s t-test. **F.** Intratracheal transduction of Lenti-sgSafe/Cre vector to initiate tumors in *KT* and *KT;H11^LSL-Cas9^;Kat7^fl/fl^* mice. **G.** Representative H&E images of lung tumors from *KT* and *KT;H11^LSL-Cas9^;Kat7^fl/fl^* mice. Bottom panels (scale bars, 50 μm) are higher magnification images of areas in the top panel (scale bars, 100 μm). **H.** Chi-squared test of the number of mice of the indicated genotypes with low or high grades. Tumor grades were assessed blindly by a clinical pathologist (**see Methods**). **I.** Representative immunohistochemistry images of H3K14ac, H4K12ac, and H4K5ac staining in *KT* and *KT;H11^LSL-Cas9^;Kat7^fl/fl^* mouse lung tumors. Bottom panels (scale bars, 50 μm) are higher magnification images of areas in the top panel (scale bars, 100 μm). **J.** Representative western blots of FACS-isolated Lin^−^TdTom^+^ neoplastic cells from *KT* and *KT;H11^LSL-Cas9^;Kat7^fl/fl^* mice transduced with Lenti-sgSafe/Cre vector. Each lane on the western blot comes from a different mouse. **K.** Bar plots showing densitometry quantification of protein levels on western blots. Relative intensities are normalized to ACTIN. Error bars are +/− standard error of the mean, and each dot represents a different mouse. Statistical significance is determined by Student’s t-test.

**Supplemental Figure 8.**
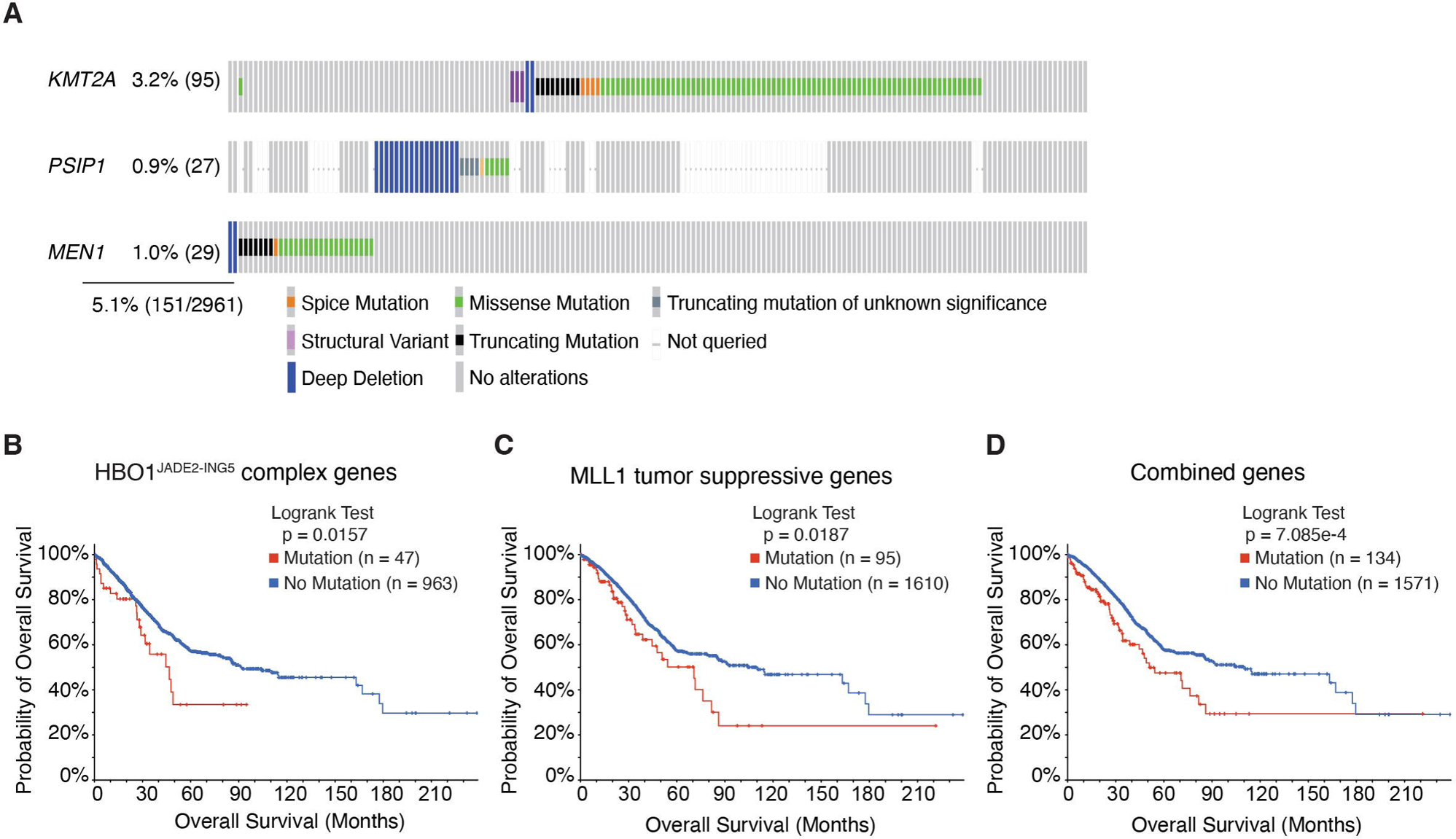
The tumor-suppressive genes in the MLL1 complex are mutated, and genetic alterations in the HBO1 and MLL1 complexes are associated with survival in human lung adenocarcinoma. **A.** Type and frequency of genetic alterations in MLL1 tumor-suppressive genes in human lung adenocarcinoma. Numbers indicate the number of patients. **B-D.** Kaplan-Meier survival curve of human lung adenocarcinoma patients with genetic alterations in *KAT7*, *JADE2*, *ING5*, or *MEAF6* (**B**), MLL1 tumor-suppressive genes *KMT2A*, *MEN1*, *PSIP1* (**C**), or in any HBO1^JADE2-ING5^ subunits or MLL1 tumor-suppressive genes compared to patients without mutations (**D**). The data is accessed from cBioPortal.

**Supplemental Figure 9.**
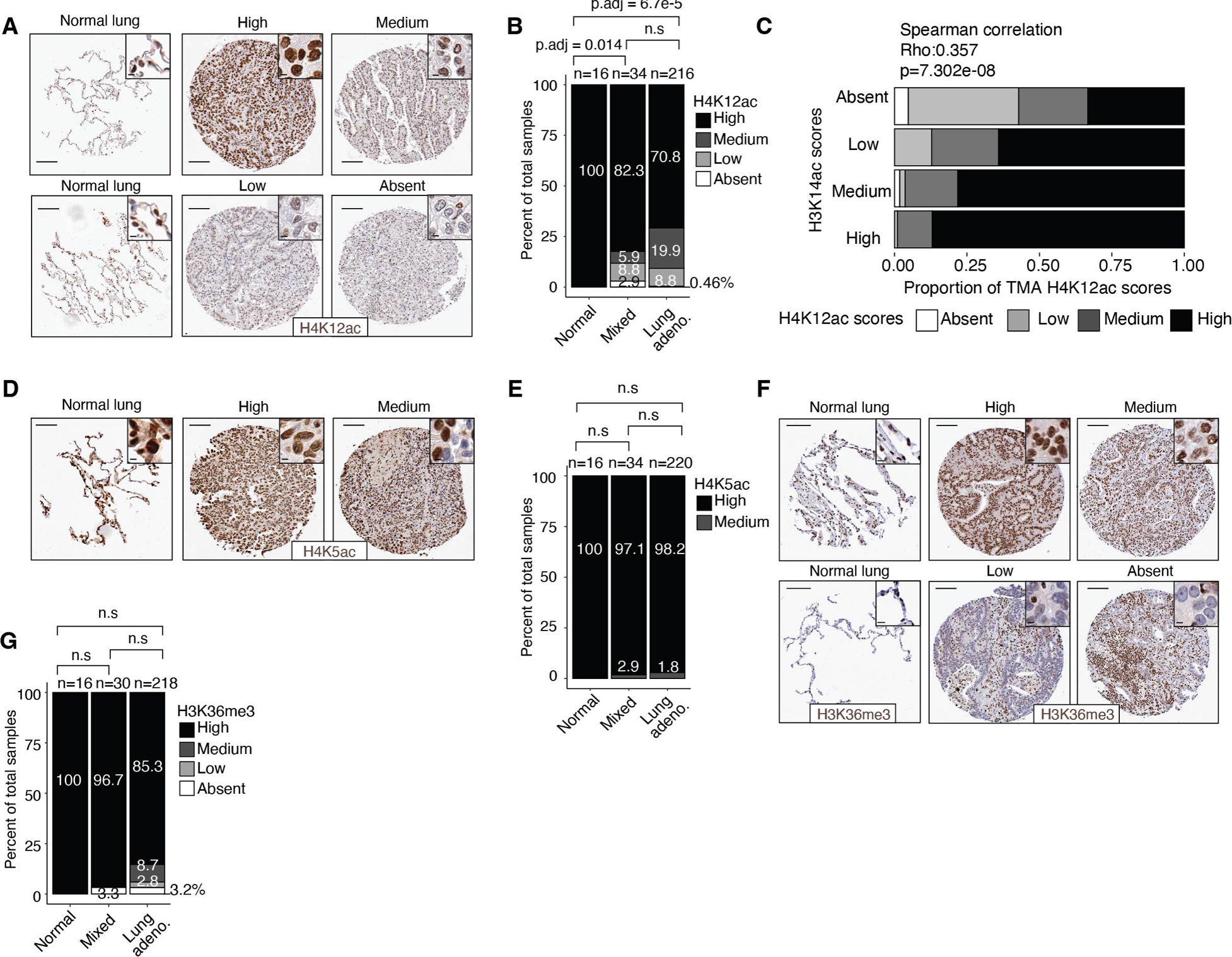
The HBO1 complex histone modification targets and H3K36me3 are disrupted in human lung adenocarcinoma. **A.** Representative immunohistochemistry images from a human lung cancer tissue microarray (TMA) expressing high, medium, low, or absent levels of H4K12ac (scale bars, 100 μm; inset scale bars, 5 μm). **B.** Bar plot showing the percentage of human lung cancer TMA expressing different levels of H4K12ac in normal lung (Normal), mixed lung adenocarcinoma with bronchioalveolar lung carcinoma (Mixed), and lung adenocarcinoma. Statistical significance is determined by Kruskal-Wallis test with Benjamini-Hochberg correction. **C.** Bar plot showing the correlation between H3K14ac and H4K12ac scores in the human lung cancer TMA. For each category of H3K14ac scores, the percentage of TMA with the specified H4K12ac scores is shown. The overall correlation between H3K14ac and H4K12ac in human lung cancer TMA is determined by Spearman’s correlation. **D.** Representative immunohistochemistry images from a human lung cancer TMA expressing H4K5ac (scale bars, 100 μm; inset scale bar, 5 μm). **E.** Bar plot showing the percentage of human TMA expressing different levels of H4K5ac in normal lung (Normal), mixed lung adenocarcinoma with bronchioalveolar lung carcinoma (Mixed), and lung adenocarcinoma. Statistical significance is determined by Kruskal-Wallis test with Benjamini-Hochberg correction. **F.** Representative immunohistochemistry images from a human lung cancer TMA expressing H3K36me3 (scale bars, 100 μm; inset scale bars, 5 μm). **G.** Bar plot showing the percentage of human lung cancer TMA expressing different levels of H3K36me3 in normal lung (Normal), mixed lung adenocarcinoma with bronchioalveolar lung carcinoma (Mixed), and lung adenocarcinoma. Statistical significance is determined by Kruskal-Wallis test with Benjamini-Hochberg correction.

**Supplemental Figure 10.**
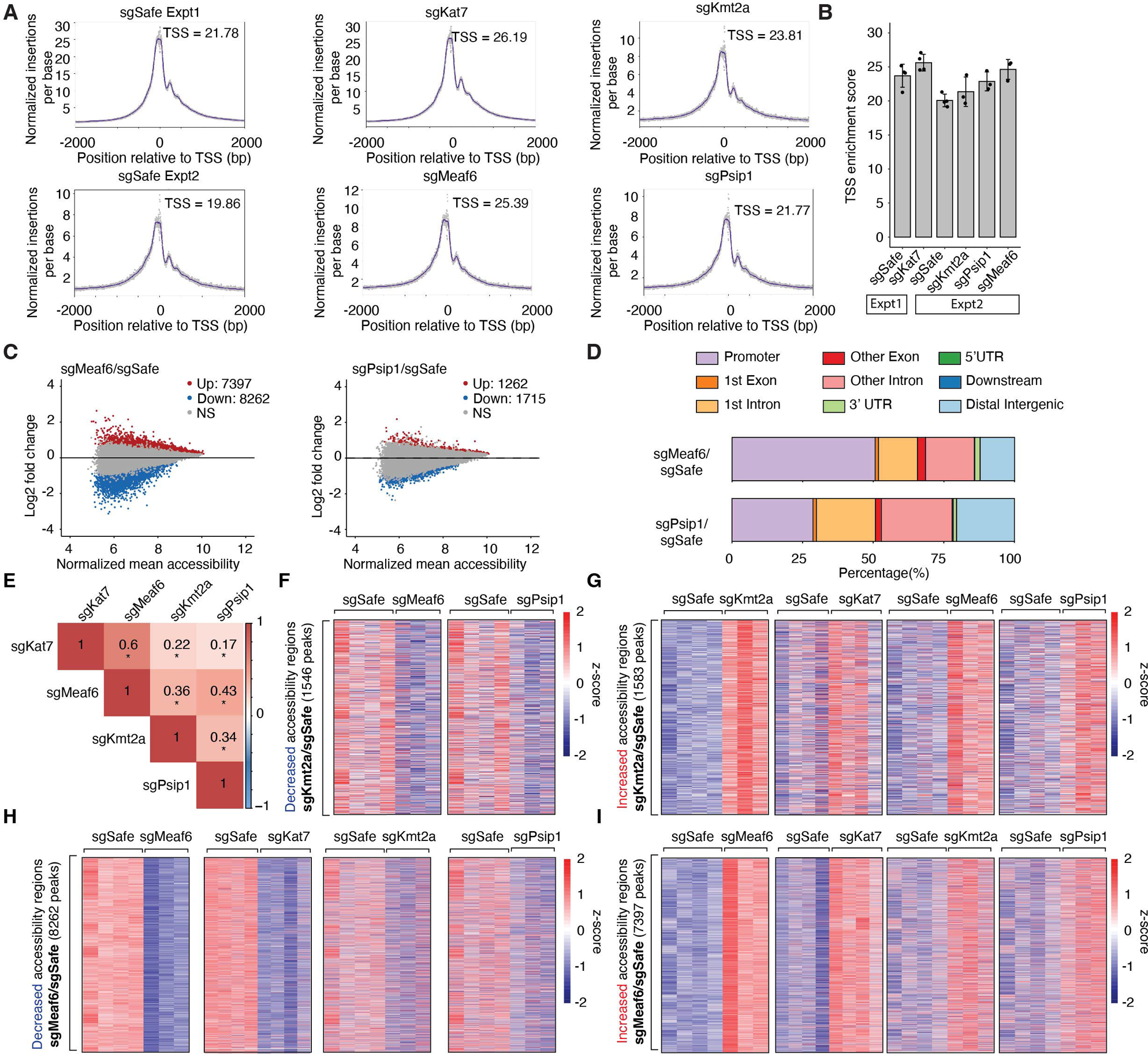
Inactivation of HBO1 or MLL1 complex genes results in highly correlated chromatin accessibility landscapes. **A.** Representative transcription start site (TSS) enrichment profiles for each gene perturbation in the ATAC-seq experiments. The TSS enrichment score is indicated on each plot. **B.** Bar plot showing the average TSS enrichment score for each gene perturbation. The dots represent the TSS enrichment score from an individual mouse. Error bars are +/− standard deviations. Horizontal bars underneath the x-axis indicate separate experiments for the different genotypes. **C.** Mean average (MA) plots for *sgMeaf6* or *sgPsip1* cells relative to *sgSafe* cells. Red and blue dots are statistically significant ATAC-seq peaks (cut-off: p.adj ≤ 0.05). **D.** Annotation and percentage of genomic regions with significantly reduced chromatin accessibility peaks for *sgMeaf6* or *sgPsip1* cells relative to *sgSafe* cells. **E.** Correlation matrix of the chromatin accessibility profile of each genotype relative to *sgSafe* cells. The numbers are Pearson correlation coefficients and * indicates p ≤ 0.05. **F.** Heatmaps showing all significantly decreased chromatin accessibility peaks in *sgKmt2a* cells relative to *sgSafe* cells (cut-off: log2 fold-change < 0 & p.adj ≤ 0.05) in the indicated genotypes. **G.** Heatmaps showing all significantly increased chromatin accessibility peaks in *sgKmt2a* cells relative to *sgSafe* cells (cut-off: log2 fold-change > 0 & p.adj ≤ 0.05) in the indicated genotypes. **H-I**. Heatmaps showing all significantly decreased (cut-off: log2 fold-change < 0 & p.adj ≤ 0.05) (**H**) or increased (cut-off: log2 fold-change > 0 & p.adj ≤ 0.05) (**I**) chromatin accessibility peaks in *sgMeaf6* cells relative to *sgSafe* cells in the indicated genotypes.

**Supplemental Figure 11.**
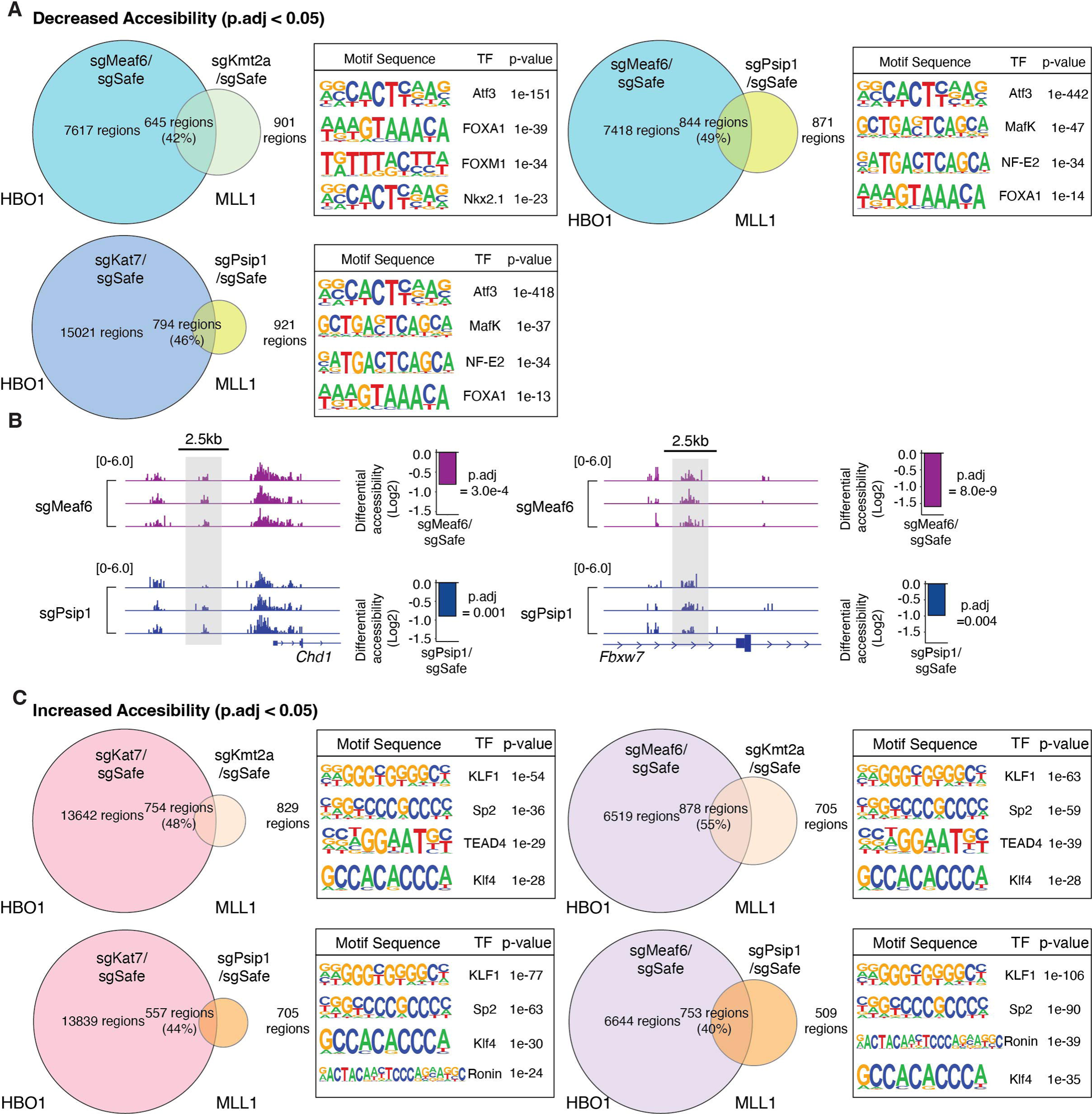
Inactivating genes in the HBO1 or MLL1 complex reduces chromatin accessibility at shared transcription factor motifs and known tumor suppressor genes. **A.** Top transcription factor motifs enriched in overlapping regions of decreased chromatin accessibility upon inactivating genes in the HBO1 or MLL1 complex. **B.** Genome accessibility tracks showing reduced chromatin accessibility upstream of transcription start sites for tumor suppressor genes *Chd1* and *Fbxw7* in *sgMeaf6* and *sgPsip1* cells. Grey regions highlight peaks with significant differential chromatin accessibility. Numbers in brackets are the range of normalized peak counts for the tracks below. Bar plots show fold-change of differential chromatin accessibility in the indicated target relative to *sgSafe* cells. **C.** Top transcription factor motifs enriched in the overlapping increased chromatin accessibility peaks between inactivating HBO1 or MLL1 complex genes.

**Supplemental Figure 12.**
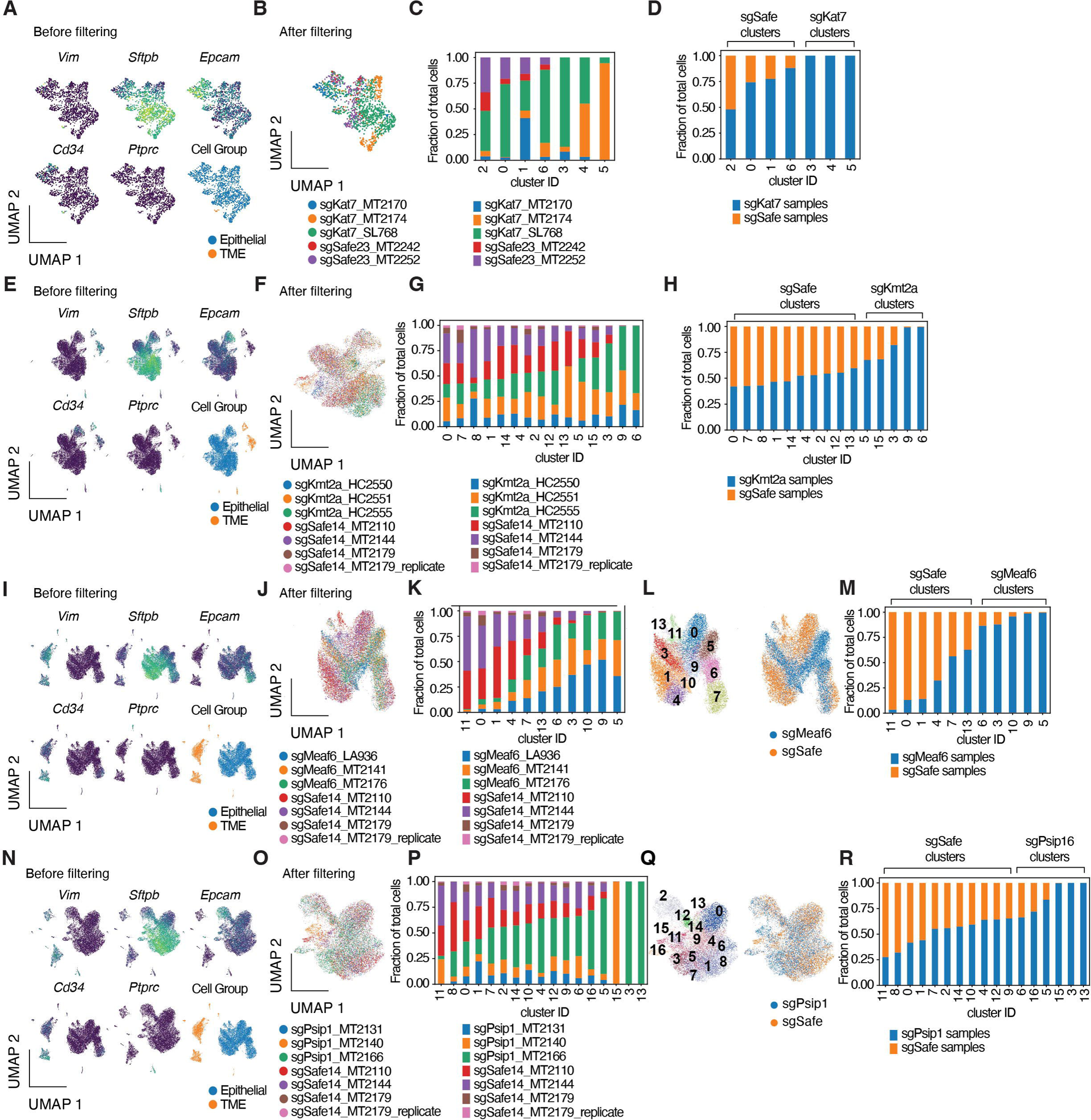
Inactivation of genes in the HBO1 or MLL1 complex alters the transcriptional state in lung tumor cells. **A.** UMAPs for *sgKat7* and *sgSafe* cells from scRNA-seq showing marker genes used to distinguish neoplastic cells and contaminating tumor microenvironment cells that were not fully eliminated during FACS. Cell clusters that express markers of non-cancerous cells are filtered (**see Methods**). Mesenchymal marker *Vim*, endothelial marker *Cd34*, and immune marker *Ptprc* are shown. **B.** UMAP of *sgKat7* and *sgSafe* cells after filtering out the tumor microenvironment cells. Different colors indicate different mice from which the cells are isolated. **C.** Bar plot showing the percentage contribution of each mouse in the UMAP clusters. Different colors indicate different mice from which the cells are isolated. **D.** Bar plot showing the percentage contribution of each genotype in the UMAP clusters for *sgKat7* and *sgSafe* cells. Clusters 3 - 5 are designated as *sgKat7* clusters (**see Methods**). **E-H.** UMAPs and bar plots showing similar information as (**A-D**) for *sgKmt2a* and *sgSafe* cells. **I-K.** UMAPs and bar plot showing similar information as (**A-C**) for *sgMeaf6* and *sgSafe* cells. **L.** UMAPs showing cell clusters and genotype of *sgMeaf6* and *sgSafe* cells. **M.** Bar plot showing the percentage contribution of each genotype in the UMAP clusters for *sgMeaf6* and *sgSafe* cells. Clusters 6,3,10,9,5 are designated as *sgMeaf6* clusters (**see Methods**). **N-R.** UMAPs and bar plots showing the similar information as (**I-M**) for *sgPsip1* and *sgSafe* cells.

**Supplemental Figure 13.**
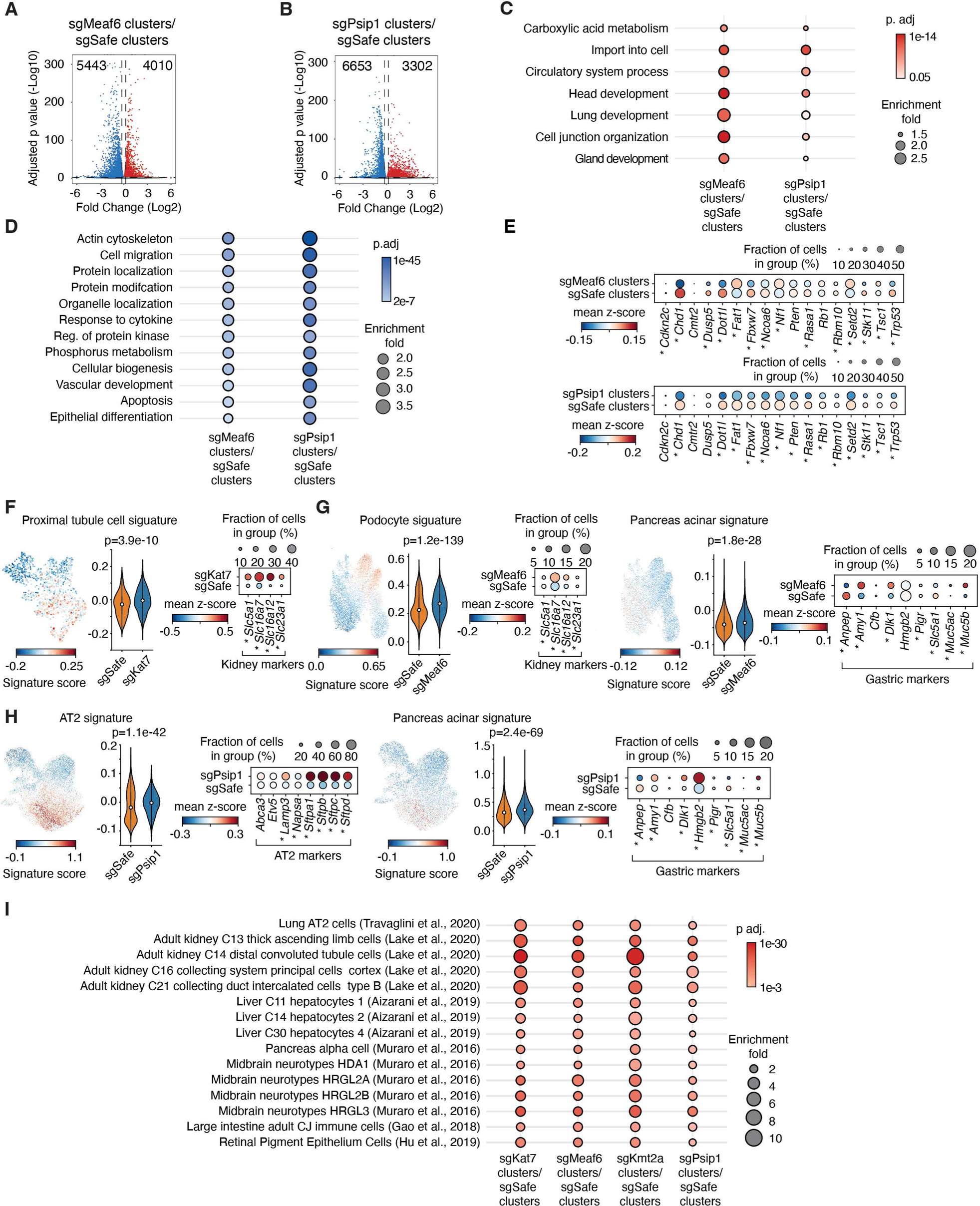
Inactivation of HBO1 or MLL1 complex genes affects shared cellular processes and disrupts lineage fidelity. **A-B.** Volcano plots of differential gene expression between cell clusters that are predominantly *sgMeaf6* (**A**) or *sgPsip1* (**B**) compared to cell clusters that are predominantly *sgSafe*. The specific clusters for each group are indicated in **Figure S12A-B** (**see Methods**). Red dots are statistically significant up-regulated genes (cut-off: log2 fold-change > 0 & p.adj ≤ 0.05), and blue dots are statistically significant down-regulated genes ((cut-off: log2 fold-change < 0 & p.adj ≤ 0.05). The total number of statistically significant genes is indicated at the top of the plot on each side. **C-D.** Enriched Gene Ontology terms of significantly up-regulated genes (**D**) or down-regulated genes (**E**) for indicated comparisons. **E.** Dot plots from scRNA-seq showing differential gene expression of canonical tumor suppressor genes for indicated comparisons. * indicates p.adj ≤ 0.05 (Student’s t-test). **F.** UMAPs showing cell type signature scores for *sgKat7* and *sgSafe* transduced tumor cells. Violin plots showing cell type signature scores of cells in *sgKat7* and *sgSafe* predominate clusters. Median cell type signature score is represented by the dot in each violin plot. Statistical significance is computed using Mann-Whitney-U test. Dot plots show relevant marker genes for cell clusters that are predominately *sgKat7* or *sgSafe*. **G.** UMAPs, violin plots and dot plots showing similar information as (**F**) for *sgMeaf6* and *sgSafe* cells. **H.** UMAPs, violin plots and dot plots showing similar information as (**F**) for *sgPsip1* and *sgSafe* cells. **I.** Dot plot showing common cell type signatures enriched for the indicated comparisons.

**Supplemental Figure 14.**
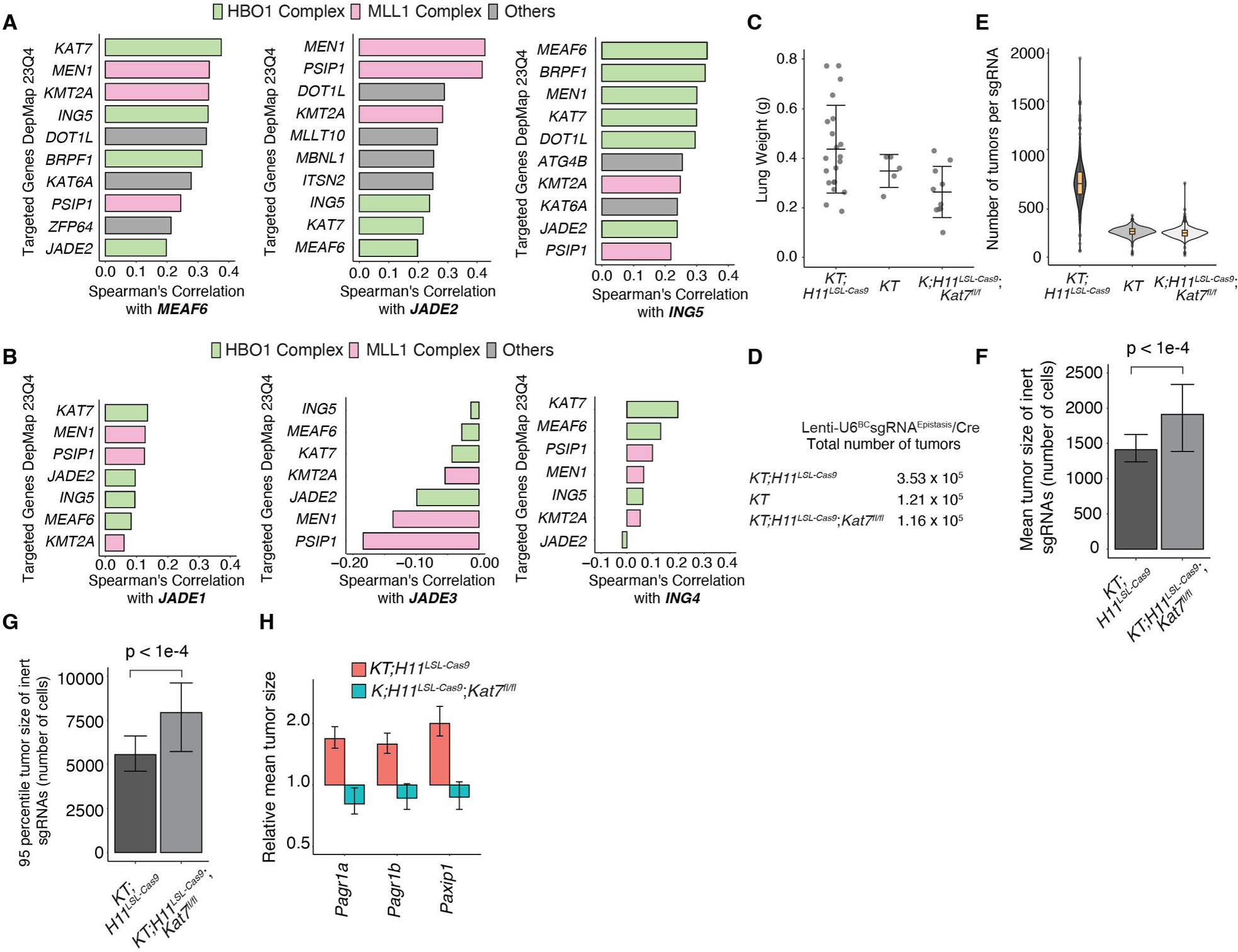
Cancer dependency score correlation and tumor growth metrics from Lenti-U6^BC^sgRNA^Epistasis^/Cre library. **A.** Correlation of DepMap cancer dependency scores showing the top genes that correlate with genes in the HBO1^JADE2-ING5^ complex. **B.** Reduced correlation of DepMap cancer dependency scores between genes from the HBO1 and MLL1 complexes with *JADE1/3* and *ING4*. **C.** Weight of tumor-bearing lungs for the indicated mouse genotypes. Each dot represents a mouse. Error bars are mean +/− standard deviation **D.** Total number of tumors analyzed in the indicated mouse genotypes. **E.** Number of tumors analyzed for each sgRNA in the indicated mouse genotypes. Boxes are the first to third quartile, and the median is indicated by the horizontal line. The whiskers are Interquartile Range (IQR) multiplied by 1.5 in each direction. **F-G.** Bar plots showing the mean (**F**) and 95th percentile size of tumors (**G**) transduced with inert sgRNAs in *KT;H11^LSL-Cas9^* and *K;H11^LSL-Cas9^;Kat7^fl/fl^*mice. Error bars show 95% confidence intervals, and p-values are calculated by two-sided Wilcoxon rank-sum test. **H.** Mean tumor size relative to inert sgRNAs for genes that interact with *Stag2* to regulate cohesin function in *KT;H11^LSL-Cas9^*and *K;H11^LSL-Cas9^;Kat7^fl/fl^* mice. Error bars are 95% confidence intervals.

**Supplemental Figure 15.**
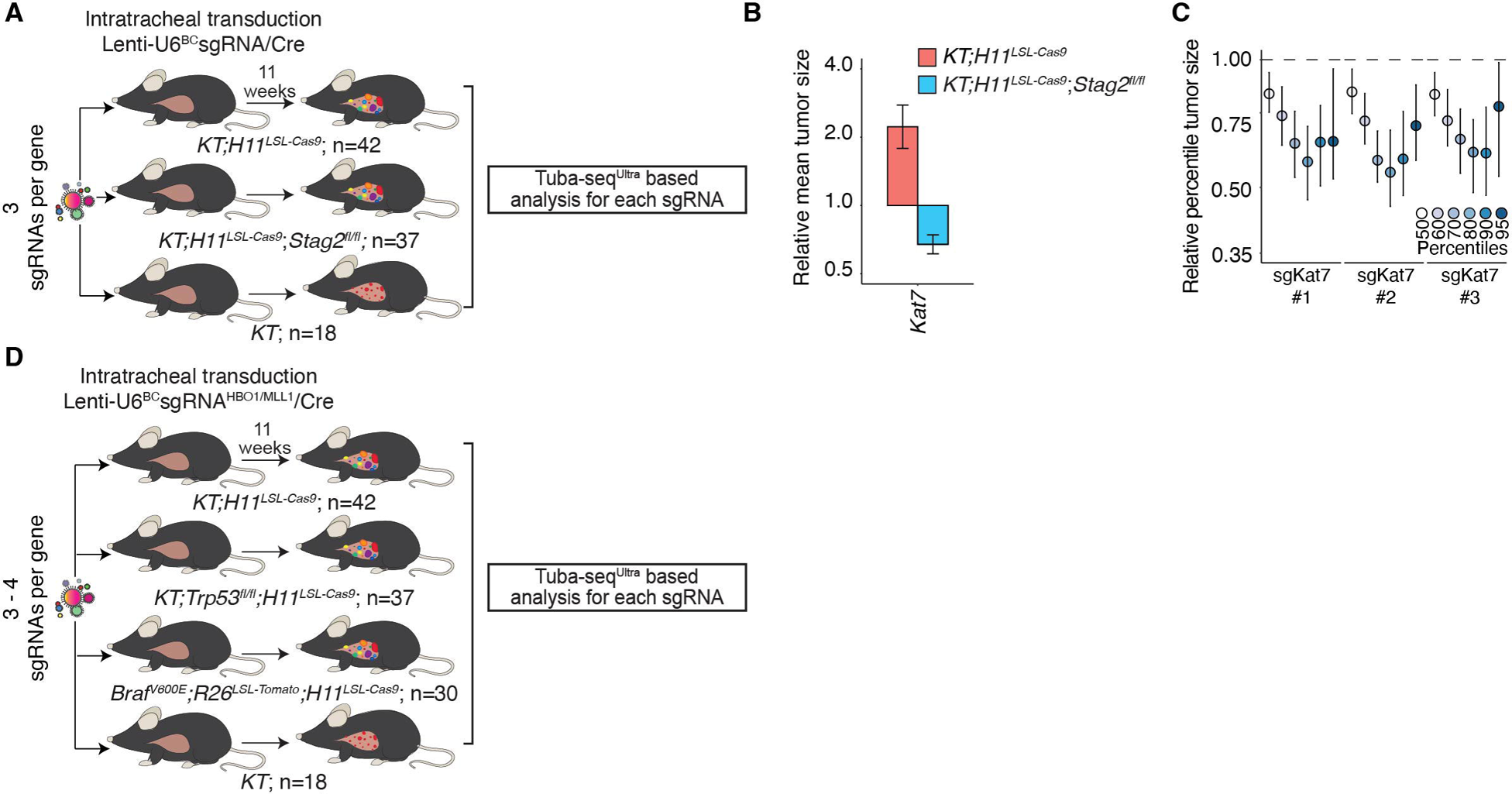
Lung tumor growth phenotypes resulting from perturbing the HBO1^JADE2-ING5^ or MLL1 complex in multiple genetic driver contexts. **A.** Intratracheal transduction of sgRNAs targeting *Kat7* to initiate tumors in *KT;H11^LSL-Cas9^*, *KT;H11^LSL-Cas9^;Stag2^fl/fl^* and *KT* mice. **B.** The mean tumor size relative to inert sgRNAs when targeting *Kat7* in *KT;H11^LSL-Cas9^* and *KT;H11^LSL-Cas9^;Stag2^fl/fl^* mice. **C.** Tumor sizes at the indicated percentiles for sgRNAs targeting *Kat7* in *KT;H11^LSL-Cas9^;Stag2^fl/fl^* mice. Error bars are 95% confidence intervals. The dotted line indicates no effect. **D.** Intratracheal transduction of sgRNAs targeting genes in HBO1^JADE2-ING5^ or MLL1 complex to initiate tumors in *KT;H11^LSL-Cas9^*, *KT;Trp53^fl/fl^;H11^LSL-Cas9^*, *Braf^V600E^;R26^LSL-Tomato^;H11^LSL-Cas9^*, and *KT* mice.

**Supplemental Figure 16.**
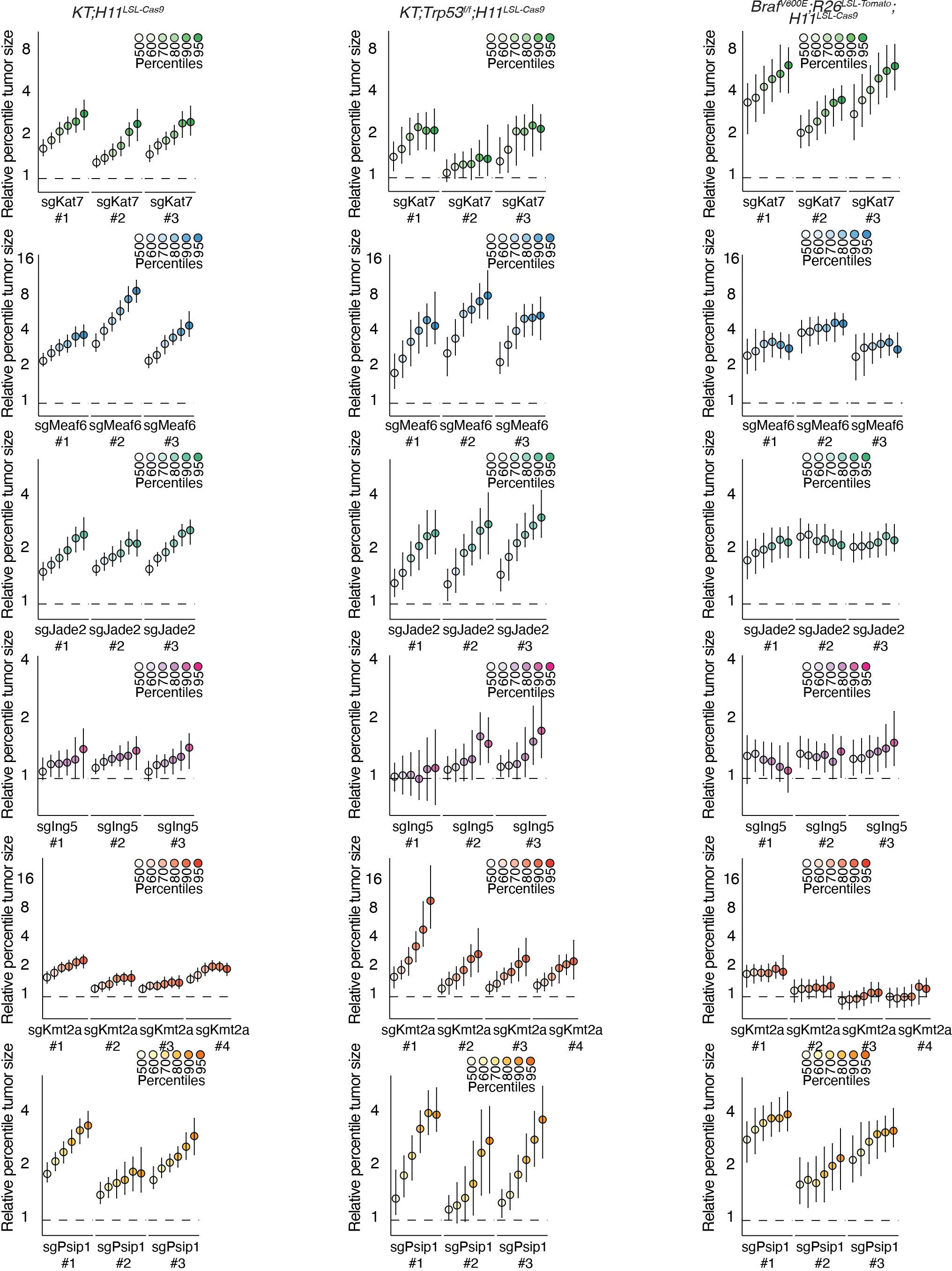
Tumor sizes at different percentiles upon perturbing the HBO1^JADE2-ING5^ or MLL1 complex in multiple genetic driver contexts. Tumor sizes at the indicated percentiles of sgRNAs targeting genes in the HBO1^JADE2-ING5^ or MLL1 complexes in *KT;H11^LSL-Cas9^*, *KT;Trp53^fl/fl^;H11^LSL-Cas9^*, and *Braf^V600E^;R26^LSL-Tomato^;H11^LSL-Cas9^* mice.

